# Umbilical cord blood-derived cell therapy for perinatal brain injury: A systematic review & meta-analysis of preclinical studies - Part B

**DOI:** 10.1101/2022.10.20.513105

**Authors:** Elisha Purcell, Timothy Nguyen, Madeleine Smith, Tayla Penny, Madison CB Paton, Lindsay Zhou, Graham Jenkin, Suzanne L Miller, Courtney A McDonald, Atul Malhotra

## Abstract

**Introduction:** We previously described preclinical literature, which supports umbilical cord blood-derived cell (UCBC) therapy use for perinatal brain injury. However, efficacy of UCBCs may be influenced by different patient populations and intervention characteristics.

**Objectives:** To systematically review effects of UCBCs on brain outcomes in animal models of perinatal brain injury across subgroups to better understand contribution of model type (preterm versus term), brain injury type, UCB cell type, route of administration, timing of intervention, cell dosage and number of doses.

**Methods:** A systematic search of MEDLINE and Embase databases was performed to identify studies using UCBC therapy in animal models of perinatal brain injury. Subgroup differences were measured by chi^2^ test where possible.

**Results:** Differential benefits of UCBCs were seen in a number of subgroup analyses including intraventricular haemorrhage (IVH) vs. hypoxia ischaemia (HI) model (apoptosis white matter (WM): chi^2^=4.07; P=0.04, neuroinflammation-TNF-α: chi^2^=5.99; P=0.01), UCB-derived mesenchymal stromal cells (MSCs) vs. UCB-derived mononuclear cells (MNCs) (oligodendrocyte WM: chi^2^=5.01; P=0.03, neuroinflammation-TNF-α: chi^2^=3.93; P=0.05, apoptosis grey matter (GM), astrogliosis WM) and intraventricular/intrathecal vs. systemic routes of administration (microglial activation GM: chi^2^=7.51; P=0.02, astrogliosis WM: chi^2^=12.44; P=0.002). We identified a serious risk of bias and overall low certainty of evidence.

**Conclusions:** Preclinical evidence suggests greater efficacy for UCBCs in IVH compared to HI injury model, use of UCB-MSCs compared to UCB-MNCs, and use of local administrative routes compared to systemic routes in animal models of perinatal brain injury. Further research is needed to improve certainty of evidence found and address knowledge gaps.

**SIGNIFICANCE STATEMENT:** In neonatal medicine there is a clear need for the development of new therapies that can provide neuroregenerative benefits for infants with brain injuries. This review offers a unique and comprehensive resource to inform the development of future preclinical and clinical studies. In part A of this review, we systematically reviewed the preclinical literature surrounding UCBCs as a therapy for perinatal brain injury. In part B of this review, we investigated the effect variables, such as UCB cell type, timing of administration and dosage, have on the efficacy of UCB-derived cell therapy in animal models of perinatal brain injury. We identified UCBCs to show greater efficacy in the brain injury model of IVH compared to HI, the use of UCB-derived MSCs compared to MNCs and the use of local administrative routes compared to systemic routes. In addition to this, we identified knowledge gaps such as the limited preclinical literature surrounding the effect of dose number and sex.

## 1. INTRODUCTION

Perinatal brain injury continues to be a major cause of neonatal mortality and life-long neurological disability in both premature and term infants. The term ‘perinatal brain injury’, understood as brain injury occurring during pregnancy or around the time of birth, encompasses a wide range of neuropathologies.[1] These include conditions such as hypoxic ischaemic encephalopathy (HIE), intraventricular haemorrhage (IVH), periventricular leukomalacia and ischaemic stroke.[1-3] Perinatal brain injury is common across both developed and low to middle income nations, with certain forms such as HIE having an incidence as high as 2-4 per 1,000 live births.[4-5] Moreover, perinatal brain injuries are significant contributors to the development of a range of serious neurological sequelae including cerebral palsy (CP), which remains the most common physical disability in childhood.[6-8] The high prevalence and morbidity associated with perinatal brain injury highlights the pressing need for developing safe therapies that can effectively reduce and repair brain injuries in infants.

Despite advances in perinatal care which have markedly improved the survival rate of newborns, the available therapies offered for infants born with encephalopathy remains largely supportive.[7] The only neuroprotective option available for term born infants with HIE is therapeutic hypothermia.[9-10] However, this intervention is only shown to reduce neonatal mortality and major morbidity if started within the first 6 hours of life for a period of 72 hours and deviation from this protocol has shown to worsen neurological recovery.[11] For preterm infants with perinatal brain injury, no current intervention exists except neurosurgical intervention for worsening ventricular dilatation or hydrocephalus following IVH. In order to see further clinical improvements, new neuroprotective interventions are needed.[9-12] Current preclinical interventions under investigation include creatine, melatonin, erythropoietin, xenon, microRNAs, insulin-like growth factors and stem cell therapies.[9,13-15] Umbilical cord blood (UCB)-derived cell therapy is one of the most prominent emerging interventions in this area of research and has received a large amount of attention in both preclinical and clinical studies.[16-19]

In part A of this review, we demonstrated UCB-derived cells (UCBCs) were effective in improving both neuropathological and behavioural outcomes in preclinical models. The specific outcomes investigated were apoptosis, astrogliosis, infarct size, microglial activation, oligodendrocyte number, neuroinflammation and motor function. Importantly, when we applied the Grading of Recommendations Assessment, Development and Evaluation (GRADE) tool adapted for preclinical studies, the certainty of this evidence was deemed low.[20] In part B of this review, we investigated the heterogeneity and variation that existed within the 55 preclinical studies analysed in Part A. We aimed to systematically compare the efficacy of UCB-derived cell therapy on brain outcomes across types of perinatal brain injuries, UCB cell types, routes of intervention, timing of intervention, dosage and number of cell doses.

## 2. METHODS

This systematic review and meta-analysis was conducted according to the Preferred Reporting Items for Systematic Reviews and Meta-Analyses (PRISMA) guidelines and subgroup analysis was performed using a protocol based on The Cochrane Handbook.[21, 22] The research protocol was registered on PROSPERO (CRD42022275764).

### 2.1 SELECTION CRITERIA

Published preclinical studies of any design investigating UCBC therapy for perinatal brain injury were assessed for eligibility. A detailed summary of inclusion and exclusion criteria was outlined in Part A. Briefly, inclusion criteria consisted of animal models of perinatal brain injury, an intervention arm that used any UCB cell or subtype, comparator of no intervention or placebo and assessed structural or functional brain outcomes. Review articles, conference abstracts, studies where full text was not available and studies unable to be retrieved in English were excluded.

### 2.2 PRIMARY OUTCOMES

The primary brain outcomes of apoptosis, astrogliosis, infarct size, microglial activation, neuron number, oligodendrocyte number, neuroinflammation and motor function were outlined and investigated in Part A.

### 2.3 SEARCH STRATEGY

MEDLINE and Embase databases were searched via Ovid using a combined search strategy conducted by EP and TN. To ensure recent studies were not missed, the search strategy was conducted on June 24, 2021, April 19, 2022 and additional citation search performed in August 2022. The advanced search strategy is presented in **Supplemental 1**.

### 2.4 STUDY SELECTION PROCESS

All studies were exported into Covidence Systematic Review Software (Veritas Health Innovation, Melbourne, Australia, available at www.covidence.org). Duplicates were automatically removed using Covidence in conjunction with manual deduplication (EP, TN). Title and abstract screening and full text screening was independently performed by two reviewers (EP, TN). Disagreements were resolved via discussion with a third reviewer.

### 2.5 DATA EXTRACTION

Relevant data specified in Part A of the review was independently extracted by two review authors (EP, TN). When data was published in a figure without tables or text to ascertain values, PlotDigitizer (version 2.6.9) was used to quantify the data. For papers with missing data, specifically standardised mean difference (SMD), n number and standard deviation (SD) or standard error (SE), corresponding authors were contacted a total of three times. If authors did not respond, the paper was excluded from the meta-analysis for that particular outcome.

### 2.6 DATA SYNTHESIS

Data was synthesised using Review Manager Software for meta-analysis (RevMan, version 5.4). Due to the expected heterogeneity across continuous data measurements, we used a random-effects, inverse variance model to calculate the standardised mean difference (SMD) and 95% confidence interval (CI). The *I*^*2*^ statistic was used to measure heterogeneity, with 25% considered low, 50% considered moderate and 75% considered high heterogeneity.[21]

### 2.7 SUBGROUP ANALYSIS

We aimed to investigate if the intervention effect varied with different patient population and intervention characteristics. For each brain outcome assessed in part A, we planned to undertake a subgroup analysis of the following pre-specified variables:

⍰ Model type
⍰ Brain injury type
⍰ UCB cell type
⍰ Timing of cell administration
⍰ Route of cell administration
⍰ Cell dosage
⍰ Number of cell doses

For each subgroup analysis we considered the criteria of (i) whether a statistically significant subgroup difference was detected (ii) the covariate distribution (iii) the plausibility of the treatment effect (iv) the importance of the treatment effect and (v) the possibility of confounding.[22] Subgroup differences were measured using the chi^2^ test, which tested the difference between the pooled effect estimate (i.e. SMD) between subgroups. As recommended by The Cochrane Handbook we planned to not compare within-subgroup statistics such as SMDs.[21] We defined a statistically significant subgroup effect as one where the covariate considered in the subgroup analysis modified the treatment effect by a p-value less than 0.1 as recommended by The Cochrane Handbook.[21] The covariate distribution was taken into account by considering the number of studies and study entries included in each subgroup analysis. The plausibility of the treatment effect was evaluated by considering whether evidence currently existed for the observed treatment effect in different studies of similar interventions. The importance of the treatment effect was considered by acknowledging the size of the measured subgroup difference in the context of the limitations of the review. Finally, the possibility of confounding was also considered.

As advised by The Cochrane Handbook at least ten study entries were required to be included in the meta-analysis for subgroup analyses to be eligible.[21] Additionally, a covariate was defined as a subgroup characteristic which included a minimum of four study entries. The covariates included in subgroup analyses were informed from results of Part A and are detailed below.

#### Model type

Covariates were preterm and term models. Insufficient reporting of preterm and term models were found in studies which used mouse or rat models. Thus, after discussion with review authors, rat preterm was defined as injury induction less than post-natal day (PND) seven and mouse preterm was defined as injury induction less than PND nine.

#### Brain injury type

All brain injury models were extracted from included studies (chorioamnionitis, excitotoxic brain injury, HI, ischaemic stroke, IVH, meningitis, hyperoxia and FGR). The brain injury model covariates of HI and IVH included a sufficient number of studies for subgroup analysis.

#### UCB cell type

All UCB cell types were extracted from included studies (EPCs, CD34+ cells, CD34-cells, MNCs, monocytes, MSCs, Tregs and unrestricted somatic stem cells). The UCB cell types of MNCs and MSCs included enough studies to be included as covariates in subgroup analysis.

#### Timing of cell administration

The times of cell administration post injury induction extracted were grouped as ‘less than 24 hours’, ‘24 to 72 hours’ and ‘greater than 72 hours’. The covariates of ‘less than 24 hours’ and ‘24 to 72 hours’ included a sufficient number of studies for subgroup analysis. Before commencement of the review, we considered how the timing of ‘early’, ‘moderate’ and ‘late’ administration times in relation to humans varied across animal species. However, lack of published literature outlining how these differences vary across animals resulted in the team deciding upon the above time ranges across all species. Subsequently, caution should be taken when evaluating the results yielded from timing of cell administration.

#### Route of cell administration

All routes of cell administration were extracted from included studies (arterial, intracerebral, intranasal, intraperitoneal, intrathecal, intratracheal, intraventricular and intravenous). The routes of arterial, intraperitoneal, intrathecal, intraventricular, intravenous underwent subgroup analysis. The following routes of cell administration were combined as covariates to allow for comparison between systemic and local routes of delivery; arterial and intravenous (systemic circulation) as well as intraventricular and intrathecal (local).

#### Cell dosage

To allow for comparison of cell dosage between animal models, cell dose amounts were extracted as cells per kilogram (kg). If studies did not report this unit, reported animal weights for the specific aged animal were used to calculate this cell dose amount. Studies were divided into the three covariates of ‘25 million cells per kg’, ‘25 to 100 million cells per kg’ and ‘greater than 100 million cells per kg’. Before commencement of the review, we considered how cell dosage in relation to humans varied across animal species. For example, we considered how a ‘low dose’ varied in a rat when compared to a sheep. However, the lack of published literature investigating these differences resulted in the aforementioned dose ranges being employed across species. Subsequently, caution should be taken when evaluating the results produced from cell dosage subgroup analyses.

#### Number of cell doses

Covariates included single and multiple cell doses. Insufficient studies used a multiple cell dose regimen to allow for subgroup analysis to be performed.

### 2.8 QUALITY ASSESSMENT

Two reviewers (EP, TN) independently assessed the risk of bias of included studies using the Systematic Review Centre for Laboratory Animal Experimentation (SYRCLE) risk of bias tool.[23] Disagreements between reviewers were resolved through discussion with additional authors. Funnel plot analysis in conjunction with Egger’s test was performed to assess the presence of publication bias using MedCalc for Windows, v20.115 (MedCalc Software, Ostend, Belgium). The certainty of evidence was assessed using the GRADE tool adapted for preclinical studies.[20]

## 3. RESULTS

### 3.1 SEARCH RESULTS

In part A of our review, a PRISMA flowchart was presented. In summary, 1082 citations were identified. After the process of deduplication, 714 papers underwent title and abstract screening using predefined selection criteria. Seventy-two papers underwent full-text screening. Nineteen of these papers were excluded for incorrect population (n = 9), intervention (n = 9) and study design (n = 1). Two additional studies were identified through manual citation searching. After the screening process, a final number of 55 papers were included in this systematic review.

### 3.2 CHARACTERISTICS OF INCLUDED STUDIES

The characteristics of included studies are summarised in **Table 1**. Studies included preterm (31%) and term (69%) animal models of rats (65%), mice (16%), sheep (13%) and rabbits (6%). The models of brain injury included HI (74%), IVH (13%), ischaemic stroke (2%), chorioamnionitis (3%), meningitis (2%), FGR (2%), hyperoxia (2%) and excitotoxic brain lesions (2%). The route of cell administration included systemic circulation (arterial and intravenous) (37%), intraventricular and intrathecal (27%), intraperitoneal (27%), intranasal (3.5%), intracerebral (3.5%) and intratracheal (2%). The timing of brain injury ranged from in utero to PND14. The timing of UCB-derived cell therapy ranged from 1 hour to 7 days post injury induction. UCB cell types included MNC (60%), MSC (17%), CD34+ (10%), EPC (3%), unrestricted somatic stem cells (3%) and others (Tregs, monocytes, CD34- and CD133+). The cell dosage ranged from 0.5 million cells/kg to 800 million cells/kg. Two out of the 55 studies used multiple cell doses of UCBCs.

**Table 1.**
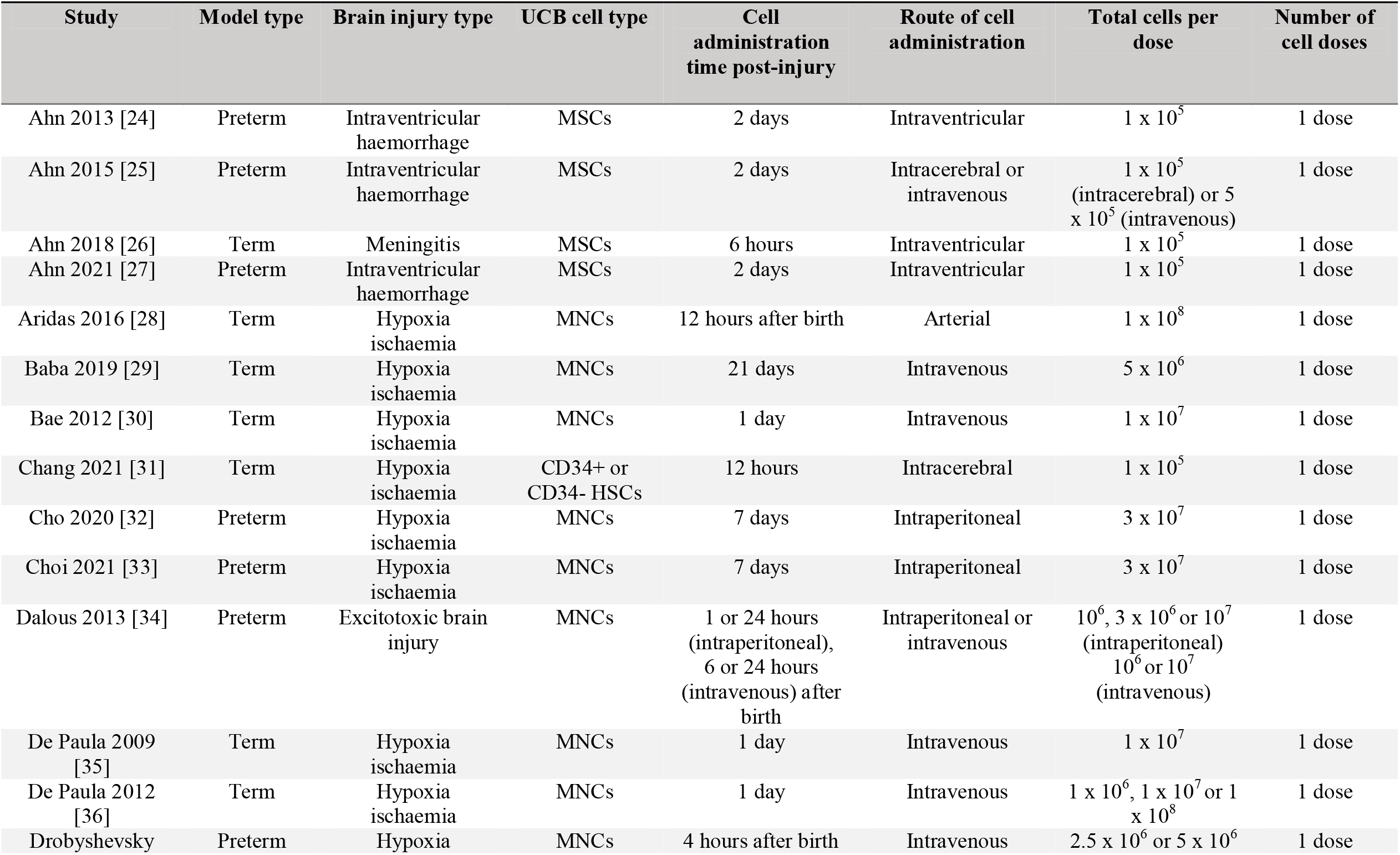
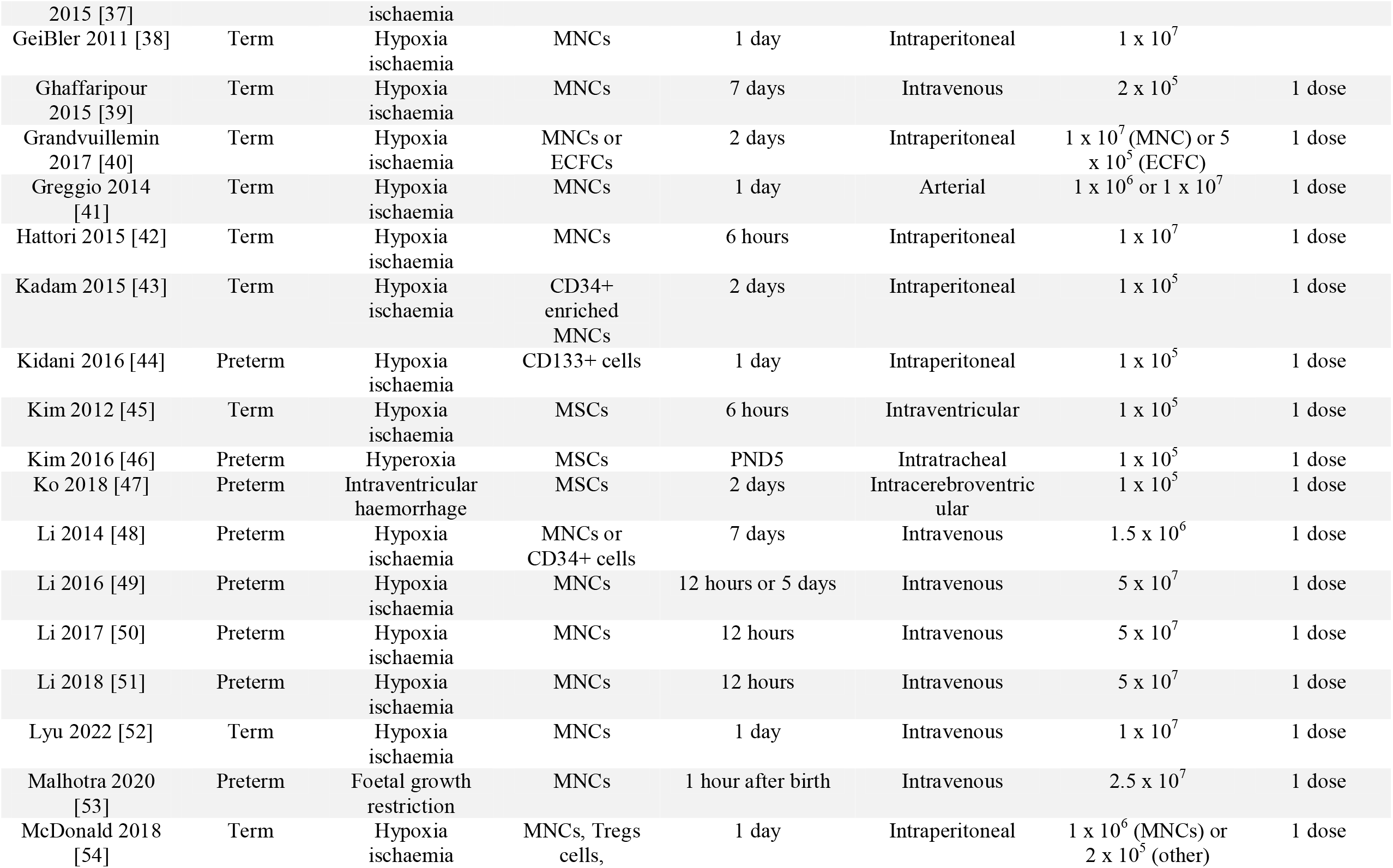
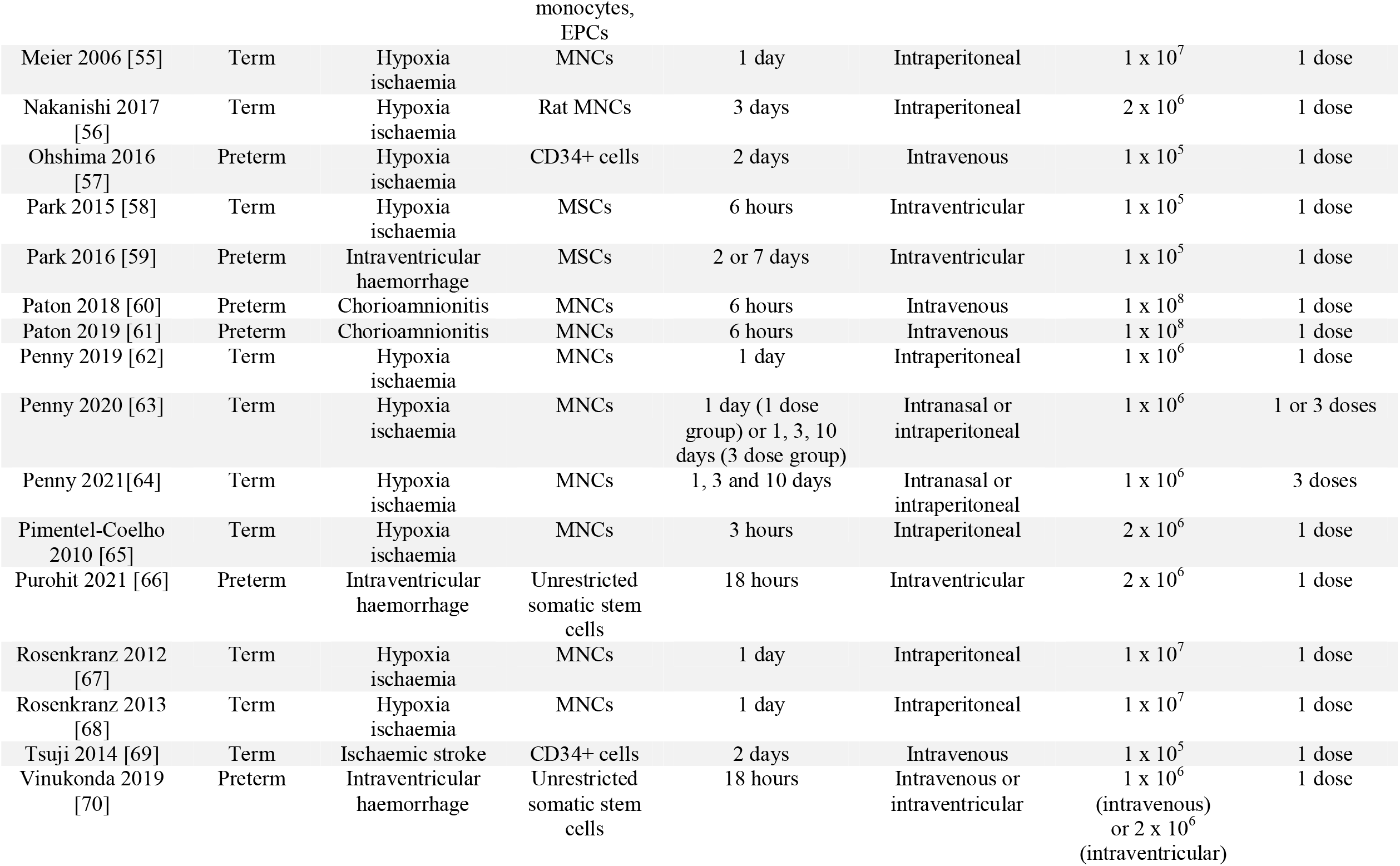
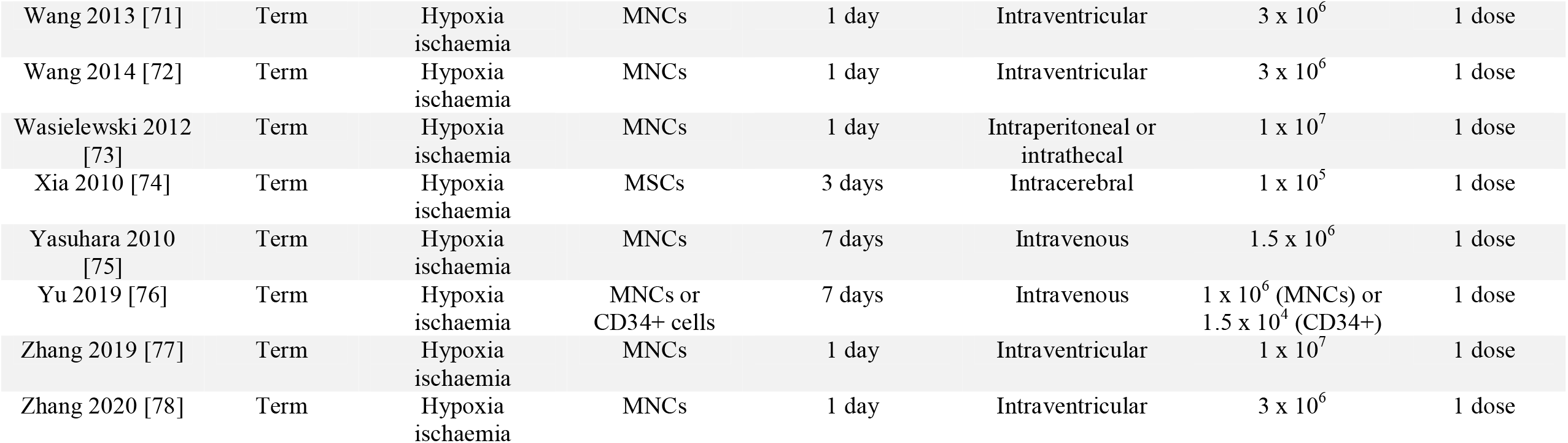
Characteristics of included studies. Abbreviations: ECFC, endothelial colony forming cells; EPCs, endothelial progenitor cells; HSCs, haematopoietic stem cells; MNCs, mononuclear cells; MSCs, mesenchymal stromal cells; PND, post-natal day; Tregs, T regulatory cells.

### 3.3 EFFECT OF UCB-DERIVED CELL THERAPY ON PRIMARY BRAIN OUTCOMES

Part A of this systematic review demonstrated that administration of UCBCs was associated with a statistically significant improvement in all evaluated brain outcomes. In summary, UCB-derived cell therapy significantly decreased infarct size, apoptosis in grey matter (GM) and white matter (WM), astrogliosis in GM and WM, microglial activation in GM and WM and neuroinflammation. UCB-derived cell therapy also improved neuron number, oligodendrocyte number in GM and WM as well as motor function. The overall certainty of results was determined as low according to the GRADE tool adapted for preclinical studies.[20]

### 3.4 EFFECT OF PRETERM AND TERM MODELS ON EFFICACY OF UCB-DERIVED CELL THERAPY

Four of eight outcomes which underwent subgroup analysis demonstrated a statistically significant difference in the efficacy of UCB-derived cell therapy between preterm and term models as summarised in **Table 2**. As shown in **Figure 1A**, microglial activation measured in GM, the test for subgroup differences detected a statistically significant subgroup effect in favour of term models (chi^2^ = 3.11, P = 0.08). Five studies (seven study entries) evaluated preterm models and eleven studies (sixteen study entries) evaluated term models. Thus, the covariate distribution was not concerning for this subgroup analysis. As shown in **Supplemental 2B**, astrogliosis in GM and **Supplemental 2D**, infarct size, statistically significant subgroup differences in favour of term models were also detected. However, in both subgroup analyses the covariate was unevenly distributed as seven of nine study entries were associated with one study in the preterm subgroup. Thus, conclusions should not be drawn from these subgroup analyses. In contrast, as shown in **Figure 1B**, oligodendrocyte number in WM, a statistically significant difference between preterm and term models was detected in favour of preterm models (chi^2^ = 14.37, P = 0.0002). Six studies (eight study entries) assessed preterm models and three studies (four study entries) assessed term models. Thus, the covariate distribution was of minimal concern for this analysis. The remaining four outcomes did not show statistically significant differences in the efficacy of UCB-derived cell therapy between preterm and term models (**Supplemental 2**).

**Table 2.** Summary of subgroup analyses. Abbreviations: * = 0.01 ≤ P < 0.1; ** = 0.001 ≤ P < 0.01; *** P < 0.001; -, subgroup analysis not performed due to insufficient number of studies; GM, grey matter; HI, hypoxia ischaemia; IVH, intraventricular haemorrhage; M/kg, million cells per kilogram; NSD, no significant difference; vs, versus; WM, white matter.

| **Outcome** | **Model type**<br>(preterm vs term) | **Brain injury type**<br>(HI vs IVH) | **UCB cell type**<br>(MNC vs MSC) | **Timing of cell administration**<br>(<24hrs vs 24-72hrs) | **Route of cell administration**<br>(intraperitoneal vs intraventricular/<br>intrathecal vs systemic) | **Cell dosage**<br>(<25M/kg vs 25-100M/kg vs >100M/kg) | **Number of cell doses**<br>(single vs multiple) |
| --- | --- | --- | --- | --- | --- | --- | --- |
| Apoptosis (GM) | NSD | - | * | NSD | NSD | NSD | - |
| Apoptosis (WM) | - | * | NSD | * | NSD | * | - |
| Astrogliosis (GM) | * | - | - | NSD | NSD | NSD | - |
| Astrogliosis (WM) | NSD | - | ** | * | ** | NSD | - |
| Infarct size | * | - | - | NSD | NSD | NSD | - |
| Microglial activation (GM) | * | - | NSD | NSD | * | NSD | - |
| Microglial activation (WM) | - | - | - | - | - | - | - |
| Neuron number | - | - | - | - | - | NSD | - |
| Oligodendrocyte number (GM) | - | - | - | - | - | - | - |
| Oligodendrocyte number (WM) | *** | - | * | - | - | NSD | - |
| Neuroinflammation (TNF-α) | NSD | * | * | * | NSD | - | - |
| Neuroinflammation (IL-6) | - | - | NSD | - | - | - | - |
| Neuroinflammation (IL-1β) | NSD | - | NSD | * | - | NSD | - |
| Neuroinflammation (IL-10) | - | - | - | - | - | - | - |
| Motor function – cylinder test | - | - | - | - | * | NSD | - |
| Motor function – rotarod test | - | - | - | - | - | - | - |

**Figure 1.**
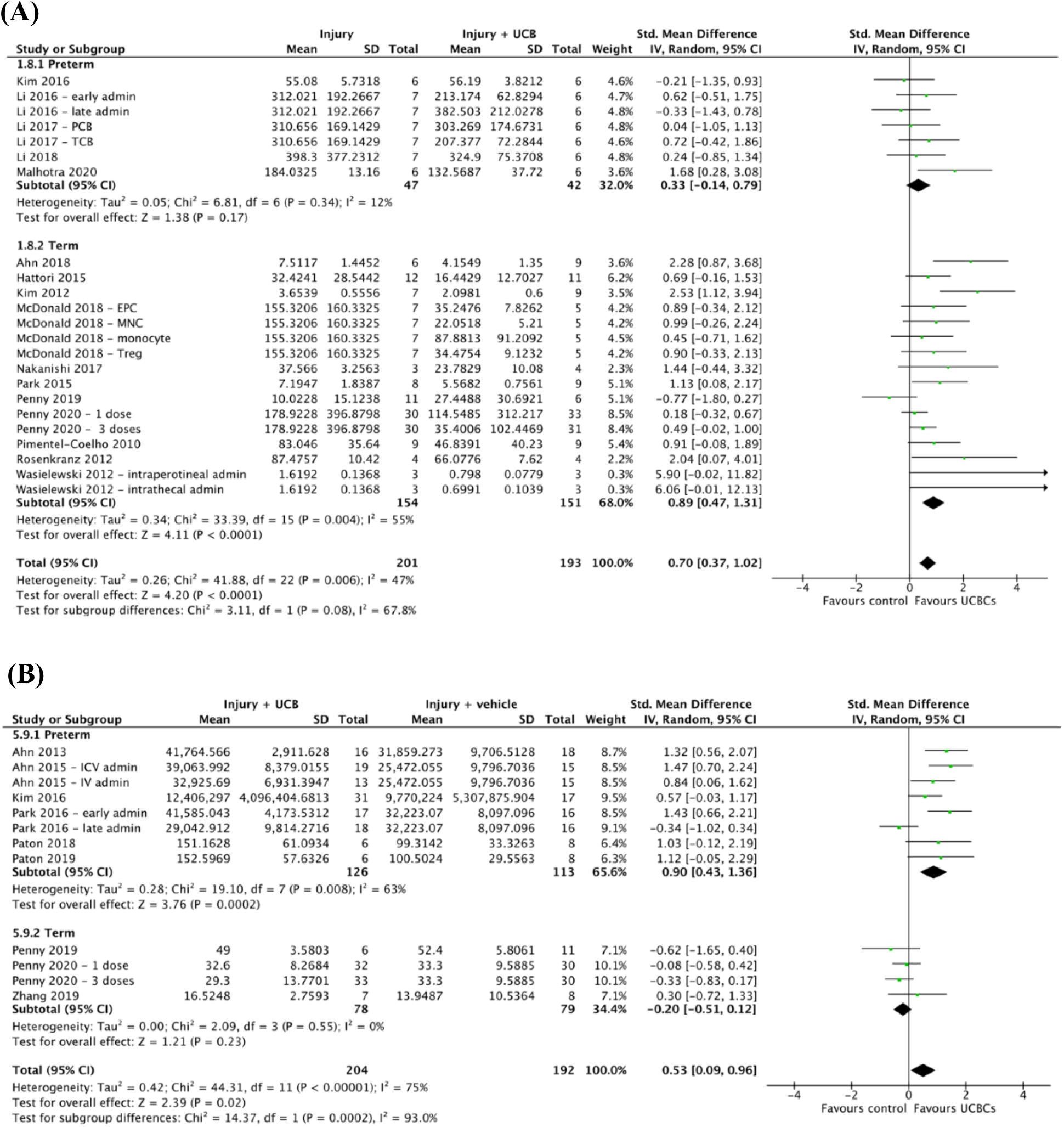
Forest plots demonstrating the effect of preterm and term models on brain outcomes of **(A)** Microglial activation – grey matter; **(B)** Oligodendrocyte number - white matter. Abbreviations: admin, administration; EPC, endothelial progenitor cell; ICV; intracerebroventricular; IV, intravenous; MNC, mononuclear cell; PCB, preterm cord blood; TCB, term cord blood; Treg, T regulatory cell.

### 3.5 EFFECT OF TYPE OF BRAIN INJURY ON EFFICACY OF UCB-DERIVED CELL THERAPY

As summarised in **Table 2**, two of two outcomes which underwent subgroup analysis of brain injury type were associated with a statistically significant subgroup difference in the efficacy of UCB-derived cell therapy. As evident in **Figure 2A**, apoptosis in WM, a statistically significant difference between HI and IVH injury models was detected in favour of IVH models (chi^2^ = 4.07, P = 0.04). The covariate distribution was not concerning for this analysis as three studies (four study entries) assessed HI and three studies (five study entries) assessed IVH. In a similar fashion, as presented in **Figure 2B**, a statistically significant difference in the efficacy of UCB-derived cell therapy on neuroinflammation as measured by TNF-α was found in favour of IVH over HI injury models (chi^2^ = 5.99, P = 0.01). Six studies (seven study entries) assessed HI models and five studies (seven study entries) assessed IVH models.

**Figure 2.**
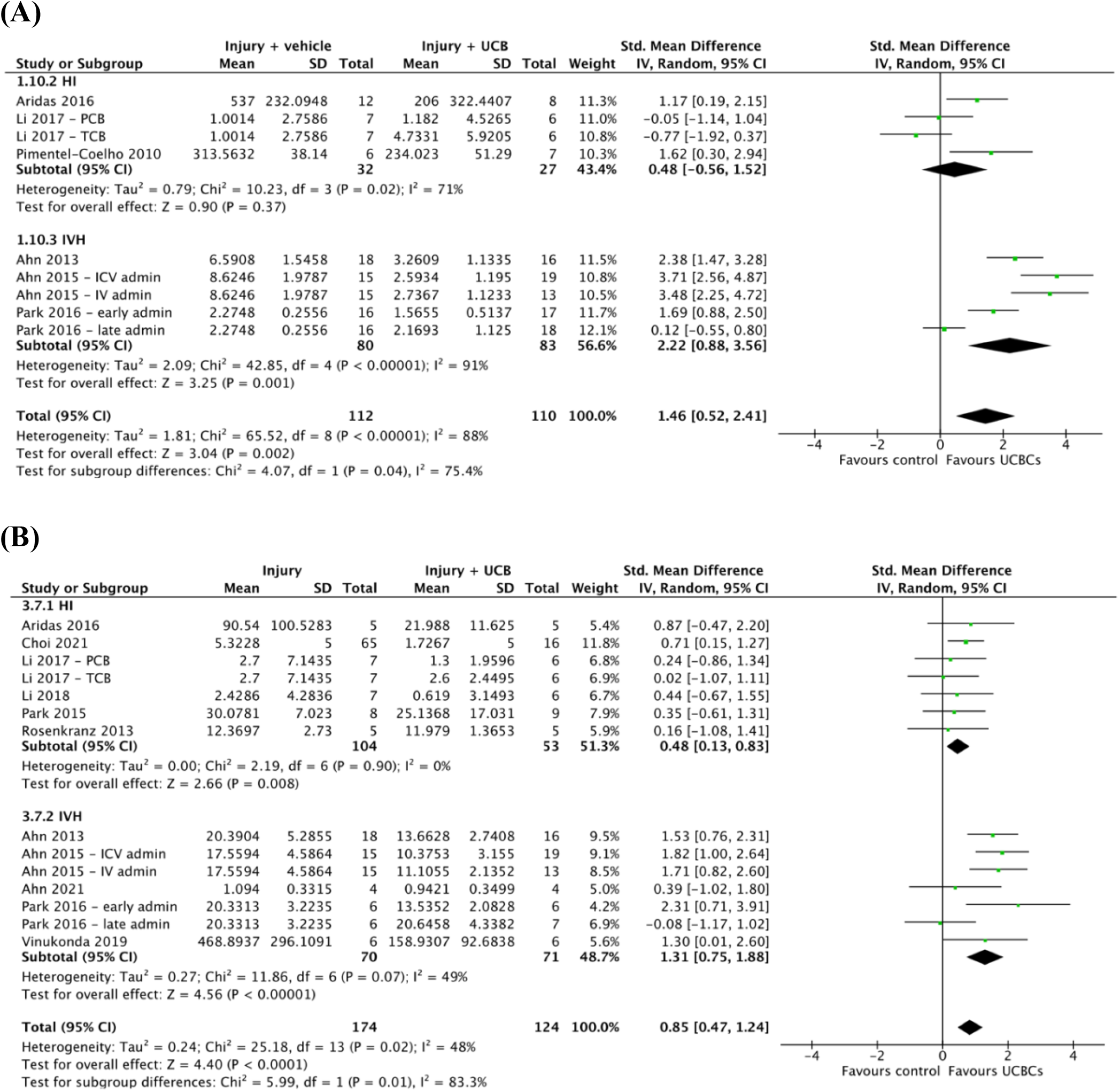
Forest plots demonstrating the effect of brain injury type on brain outcomes of **(A)** Apoptosis – white matter; **(B)** Neuroinflammation - TNF-α. Abbreviations: admin, administration; ICV, intracerebroventricular; IV, intravenous; PCB, preterm cord blood; TCB, term cord blood.

### 3.6 EFFECT OF UCB CELL TYPE ON EFFICACY OF UCB-DERIVED CELL THERAPY

Four of eight outcomes demonstrated a statistically significant difference in the efficacy of UCB-derived cell therapy between UCB cell types as detailed in **Table 2**. As shown in **Figure 3A**, a statistically significant subgroup difference between the efficacy of MNCs compared to MSCs when evaluating the outcome of oligodendrocyte number in WM was observed (chi^2^ = 5.01, P = 0.03). This modification of the treatment effect was in favour of MSCs. Five studies (six study entries) evaluated the efficacy of MNCs and four studies (six study entries) evaluated the efficacy of MSCs. Similarly, as evident in **Figure 3B**, neuroinflammation as measured by TNF-α, a statistically significant subgroup difference was detected in favour of MSCs over MNCs (chi^2^ = 3.93, P = 0.05). The covariate was evenly distributed with six studies (seven study entries) investigating MNCs and seven studies (nine study entries) investigating MSCs. Additional subgroup differences were detected in apoptosis in GM (**Supplemental 4A**) and astrogliosis in WM (**Supplemental 4C)**. Both of these subgroup analyses also detected statistically significant differences in favour of MSCs over MNCs. The remaining four outcomes demonstrated no statistically significant differences in the efficacy of UCB-derived cell therapy between MNCs and MSCs (**Supplemental 4**).

**Figure 3.**
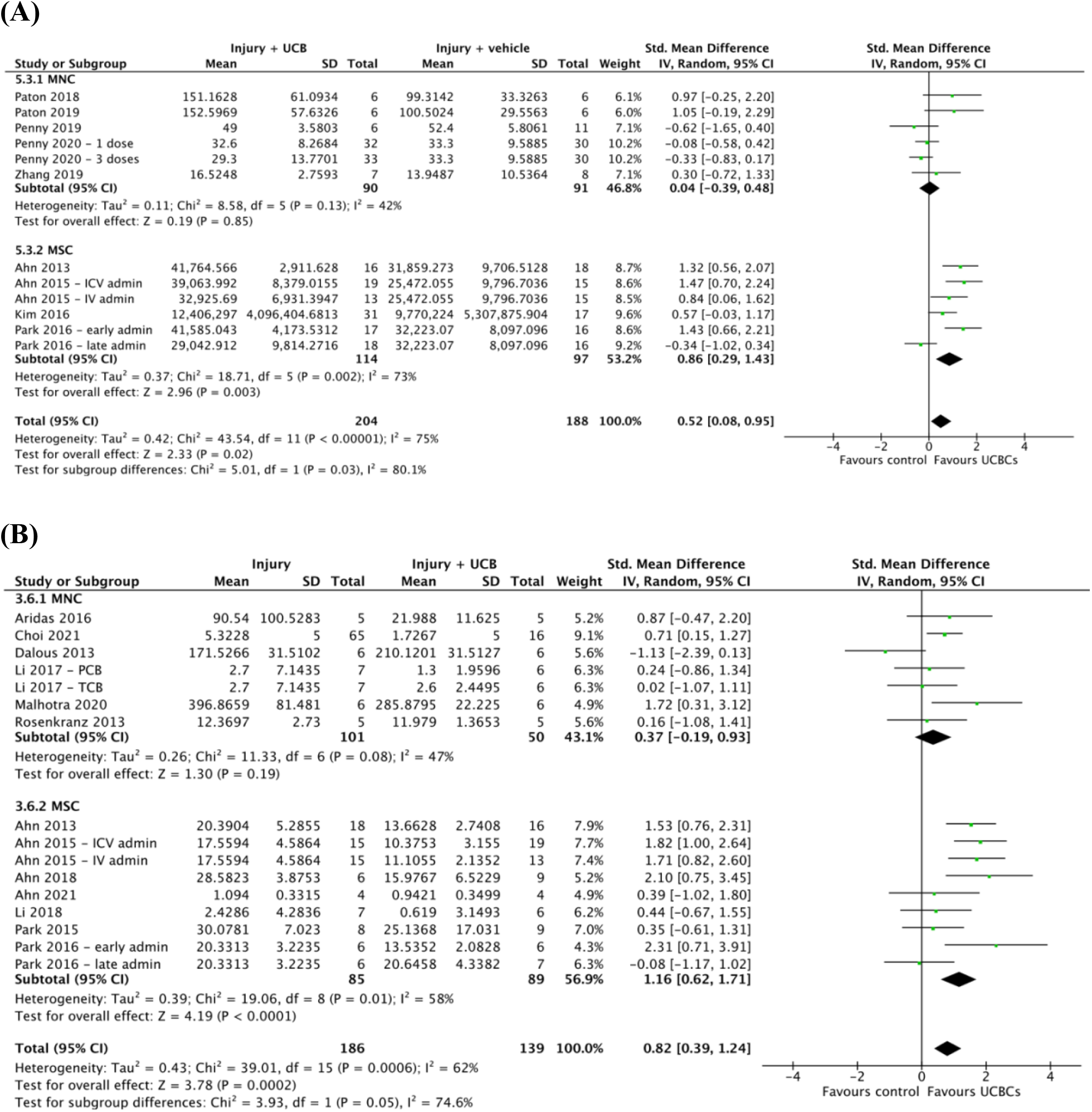
Forest plots demonstrating the effect of UCB cell type on brain outcomes of **(A)** Oligodendrocyte number - white matter; **(B)** Neuroinflammation - TNF-α. Abbreviations; admin, administration; ICV, intracerebroventricular; IV, intravenous; MNC, mononuclear cell; MSC, mesenchymal stromal cell.

### 3.7 EFFECT OF TIMING OF CELL ADMINISTRATION ON EFFICACY OF UCB-DERIVED CELL THERAPY

As summarised in **Table 2**, four of eight outcomes showed a statistically significant difference in the efficacy of UCB-derived cell therapy across different times of cell administration. As shown in **Figure 4A**, apoptosis WM, a statistically significant modification in treatment effect was seen in favour of cell administration timing of ‘24 to 72 hours’ post injury induction when compared to ‘less than 24 hours’(chi^2^ = 4.72, P = 0.03). The covariate distribution was of moderate concern for this analysis as six studies (seven study entries) formed the ‘less than 24 hours’ subgroup and three studies (four study entries) formed the ‘24 to 72 hours’ subgroup. As evident in Figure **4B**, neuroinflammation as measured by IL-1/1*β*, a statistically significant difference in subgroups of ‘less than 24 hours’ and ‘24 to 72 hours’ post injury induction was also seen in favour of ‘24 to 72 hours’ (chi^2^ = 3.31, P = 0.07). The covariate distribution was not concerning for this subgroup analysis. A similar pattern of UCB-derived cell therapy favouring ‘24 to 72 hours’ post injury induction over ‘less than 24 hours’ was also found in astrogliosis in WM and neuroinflammation as measured by TNF-α. These are presented in **Supplemental 5**. The remaining four outcomes analysed showed no statistically significant differences in the efficacy of UCB-derived cell therapy between different intervention administration times post injury induction. (**Supplemental 5**).

**Figure 4.**
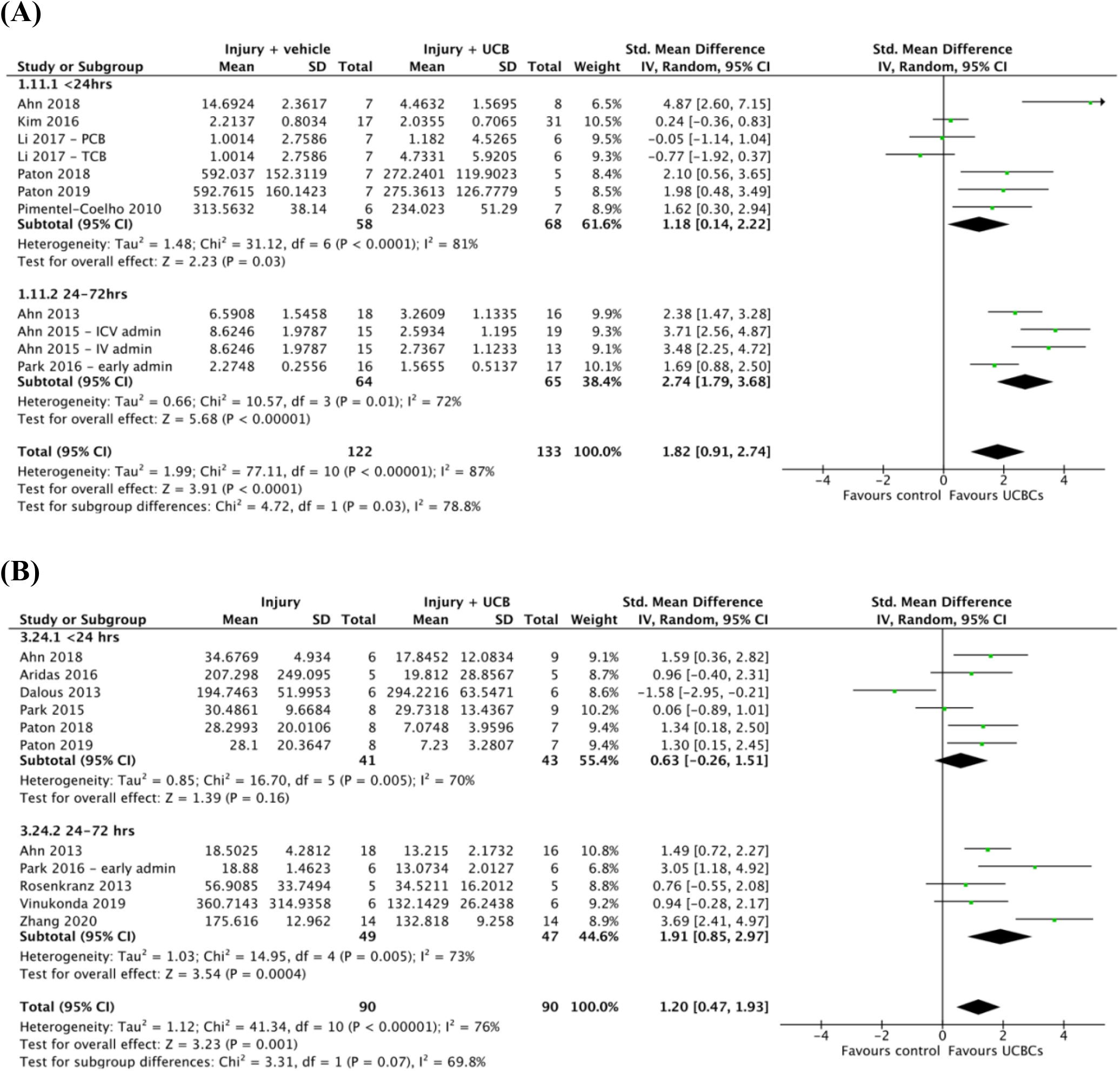
Forest plots demonstrating the effect of timing of intervention post injury induction on brain outcomes **(A)** Apoptosis - white matter; **(B)** Neuroinflammation - IL-1*β*. Abbreviations: admin, administration; ICV, intracerebroventricular; IV, intravenous; PCB, preterm cord blood; TCB, term cord blood.

### 3.8 EFFECT OF CELL ADMINISTRATION ROUTE ON EFFICACY OF UCB-DERIVED CELL THERAPY

Three of eight outcomes were shown to have a statistically significant difference in the efficacy of UCB-derived cell therapy across varying routes of cell administration as summarised in **Table 2**. As seen in **Figure 5A**, microglial activation in GM, the test for subgroup differences detected a statistically significant subgroup effect in favour of intraventricular/intrathecal administration over systemic circulation (chi^2^ = 7.51, P = 0.02). A sufficient number of trials were included in the subgroup analysis with seven studies (ten study entries) contributing to intraperitoneal route of administration, four studies (four study entries) contributing to intraventricular/intrathecal route of administration and four studies (six study entries) contributing to systemic circulation. A similar subgroup effect favouring intraventricular/intrathecal administration over intraperitoneal administration was also detected in astrogliosis in WM (**Figure 5B**, chi^2^ = 12.44, P = 0.002). In contrast, as shown in **Supplemental 6H**, motor function measured by cylinder test, a subgroup difference favouring intraperitoneal route of cell administration over intraventricular/intrathecal route was detected (chi^2^ = 6.50, P = 0.01). The covariate distribution was not concerning for this subgroup analysis as six studies (six study entries) examined intraperitoneal administration and four studies (five study entries) examined intraventricular/intrathecal route of cell administration. The remaining five outcomes demonstrated no statistically significant differences in the efficacy of UCB-derived cell therapy between cell administration routes. (**Supplemental 6**).

**Figure 5.**
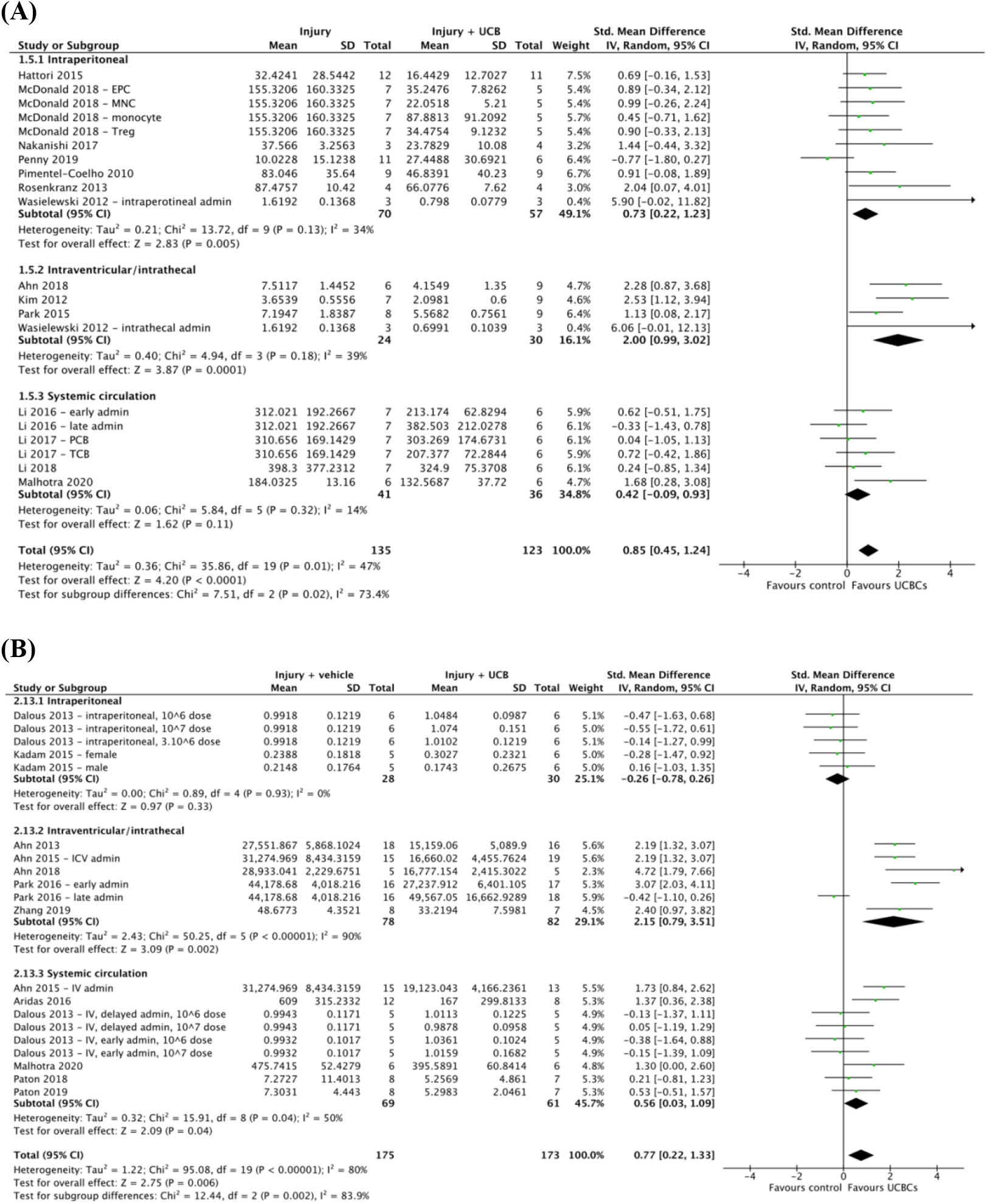
Forest plot demonstrating the effect of route of cell administration on brain outcomes **(A)** Microglial activation – grey matter; **(B)** Astrogliosis – white matter. Abbreviations: admin, administration; EPC, endothelial progenitor cell; ICV, intracerebroventricular; IV, intravenous; MNC, mononuclear cell; PCB, preterm cord blood; TCB, term cord blood; Treg, T regulatory cell.

### 3.9 EFFECT OF CELL DOSAGE ON EFFICACY OF UCB-DERIVED CELL THERAPY

As summarised in **Table 2**, one of ten outcomes demonstrated a statistically significant difference in the efficacy of UCB-derived cell therapy with different cell dosages. As shown in **Supplemental 7B**, apoptosis in WM, a statistically significant subgroup effect was found between ‘less than 25 million cells per kg’ and ’25 to 100 million cells per kg’ (chi^2^ = 5.63, P = 0.02). This modification in treatment effect favoured ‘less than 25 million cells per kg’. The covariate distribution was not concerning for this analysis as a sufficient number of studies were included in each subgroup. The remaining nine outcomes which underwent subgroup analysis of dose amount found no statistically significant differences in the efficacy of UCB-derived cell therapy between different doses (**Supplemental 7**).

### 3.10 QUALITY ASSESSMENT

Quality assessment of included studies was reported in part A of this review. The risk of bias of included studies was assessed using the SYRCLE risk of bias tool and is presented in **Supplemental 8**.[23] In summary, most biases assessed were judged ‘unclear’ due to lack of sufficient reporting. Additionally, through further assessment via the generation of funnel plots and egger’s test, publication bias was assessed as high across brain outcomes. The certainty of results was assessed using the GRADE tool adapted for preclinical studies.[20] As detailed in part A, after the assessment of risk of bias, inconsistency, imprecision, publication bias, indirectness and upgrading, the overall certainty of evidence for our findings was rated as low.

## 4. DISCUSSION

In part A of our review, we concluded UCB-derived cell therapy is an efficacious treatment in preclinical models of perinatal brain injury, with benefits seen across both neuropathological and functional outcomes. However, findings were limited by a low certainty of evidence. In this paper, we demonstrated for the first time that variations in study features and design, specifically IVH brain injury, use of UCB-MSCs and local route (around the site of injury) of administration play a statistically significant role in modifying the treatment effect seen with administration of UCB-derived cell therapy for perinatal brain injury.

### 4.1 MODEL OF BRAIN INJURY

One reason stem cell therapies receive such widespread attention in neonatal research is their unique potential to improve multiple disease states.[79] Despite this, our systematic review identified a heavy focus on HI brain injury, with 41 out of 55 studies investigating HI. Due to the limited studies investigating other forms of brain injury, we were only powered to compare the brain injury models of HI and IVH by subgroup analysis. The outcomes which demonstrated a statistically significant difference between models of brain injury were apoptosis in WM and neuroinflammation measured by TNF-α. The data from our meta-analysis suggests that UCBCs may potentially offer more significant neuropathological improvements in IVH injury when compared to the HI model of brain injury. However, it is important to emphasise this insight is heavily limited by the overall low certainty of our evidence. Additionally, high heterogeneity across studies also introduces the possibility of confounding. Nonetheless, our paper highlights the need for further studies to investigate the potential of UCBCs as therapy for brain injury models other than HI. Of the seven studies we identified that assessed the effect of UCBCs on IVH, none were performed in large animal models. Furthermore, there is no current preclinical literature which directly compares the efficacy of UCB derived cell therapy across different models of brain injury. Thus, further investigation comparing perinatal brain injuries is warranted and may offer insights into the underlying mechanisms of UCBCs as well as provide essential evidence-based data to inform the development of future clinical trials.

### 4.2 UCB CELL TYPE

UCB refers to blood within the umbilical cord and blood vessels surrounding the fetal component of the placenta.[17] Numerous cell types comprise UCB including HSCs, MSCs, Tregs, monocytes and EPCs.[81] In our review, all four outcomes which demonstrated statistically significant differences between UCB cell types were associated with a favoured modification of treatment effect in MSCs over MNCs. Our review provides further neuropathological support for UCB-MSCs as a potential therapeutic option for infants with perinatal brain injury.[13, 61, 82] To the best of our knowledge, in current literature no study is yet to directly compare UCB-MNCs to UCB-MSCs. However, Paton et al., (2019) has investigated UCB-MNCs cells to UC-MSCs and found the cell types had differential effects on WM in the preterm brain.[61] The results of our review are consistent with the reasoning for this differential effect being that MSCs comprise <0.1% of the total MNCs in UCB.[61] The beneficial effects seen in our review are also consistent with a recent systematic review performed by Lehnerer et al., (2022) which found that administration of MSCs (sourced from bone marrow, UCB, placenta, Wharton’s jelly and adipose tissue) significantly favoured sensorimotor and cognitive performance in perinatal arterial ischaemic stroke injured animals.[82] Despite our review findings supporting the use of UCB-MSCs over UCB-MNCs, particularly in the context of WM microstructure, it is important to highlight other UCB cell types, such as Tregs and monocytes, did not have sufficient studies to be included in subgroup analysis. Moreover, it is essential to understand our review findings in the context of our quality assessment, which found that overall certainty of our results was low, primarily due to the high heterogeneity between studies.

### 4.3 ROUTE OF CELL ADMINISTRATION

A range of UCB-derived cell therapy delivery routes have been investigated in preclinical literature. UCBCs can be delivered locally around the site of injury (intracerebral, intraventricular, intrathecal, intranasal) or systemically (intravenous, intraarterial, intraperitoneal).[7] In our review, three outcomes showed statistically significant differences between the method of delivery. Two of these outcomes (astrogliosis in WM and microglial activation in GM) favoured intraventricular/intrathecal administration over systemic routes and the third outcome (motor function measured by cylinder test) favoured intraperitoneal route of cell administration over local routes. This data is suggestive that UCB-derived cell therapy may potentially be more effective on neuropathological outcomes when UCBCs are administered locally to the injured site where they have been shown to have effects via cell-to-cell contact in addition to paracrine mechanisms.[9, 82, 83] In the preclinical space, only three studies have directly compared the routes of intracerebral, intraventricular or intrathecal administration to another route of cell administration. Wasielewski et al., (2012) compared the routes of intrathecal to intraperitoneal, Ahn et al., (2015) compared intracerebral to intravenous and Vinukonda et al., (2019) compared intraventricular to intravenous routes.[25, 70, 73] Further studies comparing intraventricular or intracerebral routes of delivery to other less invasive local routes such as intranasal delivery and systemic routes would be valuable additions to the current preclinical literature. Additionally, in the clinical setting, the majority of trials have implemented intravenous routes of administration.[16] To the best of our knowledge, there has been one phase one trial using intraventricular transplantation and this was shown to be safe and feasible in extremely premature infants with severe IVH.[84] In comparing this trial to clinical trials performed in children with cerebral palsy, intrathecal and intraventricular delivery of stem cells have also been shown to have no inferior safety profile to systemic routes in early phase trials.[85, 86] Thus, further research is needed to be done investigating the safety profile, feasibility and efficacy of local administration routes of UCBCs.

### 4.4 LIMITATIONS

We acknowledge there are limitations of this review. Of most importance is the high heterogeneity within the studies investigated. Studies included in the review varied across animal species, brain injury models, UCB cell types, administration routes, cell dosage, measurement tools and animal sex. Although such heterogeneity enabled subgroup analyses to be performed, the substantial heterogeneity significantly reduced the overall certainty and validity of the evidence. For example, across different animal species an early cell administration time point and high cell dose amount in relation to humans vary considerably. Subsequently, caution should be taken when evaluating the results yielded from timing of cell administration and cell dosage. Moreover, with high heterogeneity between studies, significant limitations arise from analysing subgroups in isolation. For instance, when comparing MSCs to MNCs, the dose range, animal species and injury type varies across studies and thus the possibility of confounding variables is a significant limitation. Additionally, despite our best efforts to retrieve missing data from respective authors, a number of studies were excluded from respective meta-analyses due to missing data. Thus, as discussed in Part A of this review, through GRADE analysis the overall certainty of evidence is considered low due to factors such as heterogeneity and serious risk of bias seen across studies.

Furthermore, our review included distinct treatment groups of the same study as individual study entries. Although this method has been implemented across several past reviews, when a limited number of study entries exist in a subgroup, the effect seen in one particular study can substantially influence the overall SMD seen for that subgroup.[21] Similarly, when evaluating the results of this review caution should be taken when subgroups included a limited number of studies. In addition to this, the size of subgroup differences detected should also be noted. Fifteen of the eighteen subgroup analyses which detected a statistically significant subgroup difference were measured as a p value between 0.001 to 0.1. By incorporating a larger number of studies and minimising heterogeneity across studies, our results and findings may have altered.

Another limitation was additional subgroup analyses were unable to be performed. Recent literature has shown administration of multiple cell doses is an important factor in the efficacy of UCBCs.[63] However, due to the limited number of studies which implemented a multiple dose regimen, we were not powered to undertake a subgroup analysis of single versus multiple cell doses. Other subgroup analyses such as the impact of physical sex were also not considered for this review. Additionally, functional outcomes assessed were limited to motor function as measured by rotarod and cylinder tests due to the wide variation in tests used across studies. Standardisation of such clinically important outcomes will provide greater power for future meta-analyses and thus allow for more robust preclinical evidence. Finally, in this review we limited our focus to UCBCs. It is important to note there are other sources of cells which have shown potentially neuroregenerative effects such as cells derived from umbilical cord tissue, bone marrow, amnion and placental tissue.[8, 12]

### 4.5 FUTURE DIRECTIONS

With the increasing number of clinical trials showing beneficial results, the use of UCB-derived cell therapy in the treatment of infants with brain injuries is an exciting possibility. However, this review has demonstrated that further preclinical research is warranted to progress UCB-derived cell therapy along the research pipeline. We recommend continued research of UCB-derived cell therapy in the context of preterm versus term models, physical sex, brain injury models other than HI such as IVH, cell types other than MNCs, timing of administration particularly greater than 72 hours post injury, effect of local routes of administration such as intranasal compared to other local and systemic routes, the effect of cell dosage and the use of multiple cell doses. Additionally, further research should be performed in large animal models where feasible. To improve the quality of preclinical evidence, we recommend future studies to pre-register study protocol, adopt standardised tests for measuring functional outcomes, report methodology in greater detail such as use of blinding, randomisation, animal sex, survival rate, dosage in cell/kg, sample numbers and specify error bars as SD or SEM. In addition to this, future research should investigate how across species we define a preterm or term model, low to high cell dosages and early to late timing of interventions. Forming standardised definitions of such characteristics across animal species will greatly improve the power of future systematic reviews and yield further needed evidence. In summary, further preclinical research into UCB-derived cell therapy for perinatal brain injury is needed to determine and confirm optimal cell type, timing of administration, route of administration, cell dosage and dose number across varying brain injuries.

## 5. CONCLUSIONS

This systematic review and meta-analysis of 55 preclinical studies identified UCBCs to show greater efficacy in the brain injury model of IVH compared to HI, the use of UCB-derived MSCs compared to MNCs and the use of local administrative routes compared to systemic routes. Additional preclinical research, particularly in large animal models, are required so that we can further identify and confirm differences in the efficacy of UCB-derived cell therapy across all investigated variables in addition to dose number and sex. Research in such areas is crucial to aid in the translation of UCB-derived cell therapy to the clinical setting.

## Contributions and Acknowledgements

E.P. and T.N.: conception and design, literature searching, collection and/or assembly of data, data analysis and interpretation, risk of bias assessment, manuscript writing; M.S.: conception and design, data analysis and interpretation, risk of bias assessment, manuscript editing; T.P., M.P., L.Z., G.J and S.M.: conception and design, manuscript editing; C.M. and A.M.: conception and design, literature searching, data analysis and interpretation, risk of bias assessment, manuscript editing, supervision

## Abbreviations and symbols

ECFC: endothelial colony forming cell
EPC: endothelial progenitor cell
FGR: fetal growth restriction
HI: Hypoxia Ischemia
HIE: Hypoxic Ischemic Encephalopathy
HSC: haematopoietic stem cell
IV: intravenous
MNC: mononuclear cell
MSC: mesenchymal stromal cell
PBS: phosphate buffered saline
PND: post-natal day
Treg: T regulatory cell
UCB: Umbilical Cord Blood
UCBC: umbilical cord blood-derived cell.

## REFERENCES

1. Novak, C.M., Ozen, M., and. Burd, I., Perinatal Brain Injury. Clin Perinatol, 2018. 45(2): p. 357–375.

2. Hagberg, H., et al., The role of inflammation in perinatal brain injury. Nat Rev, Neurol, 2015. 11: p. 192–208.

3. McNally, M.A., and Soul, J.S., Pharmacological Prevention and Treatment of Neonatal Brain Injury. Clin Perinatol, 2019. 46(2): p. 311–325.

4. Hagberg, H., Edwards, A.D., and Groenendaal, F. Perinatal brain damage: The term infant. Neurobiol Dis, 2016. 92(Pt A): p. 1022–112.

5. Placha, K., et al., Neonatal brain injury as a consequence of insufficient cerebral oxygenation. Neuro Endocrinol Lett., 2016. 37(2): p. 79–96.

6. Larroque, B., et al., Neurodevelopmental disabilities and special care of 5-year-old children born before 33 weeks of gestation (the EPIPAGE study): a longitudinal cohort study. Lancet, 2008. 371(9615): p. 813–820.

7. Peng, X., et al., Umbilical cord blood stem cell therapy in premature brain injury: Opportunities and challenges. J Neurosci Res, 2020. 98: p. 815–825.

8. McDonald, C.A., et al., Umbilical cord blood cells for treatment of cerebral palsy; timing and treatment options. Paediatr Res, 2018. 83: p. 333–344.

9. Davidson, J.O., et al., Perinatal brain injury: mechanisms and therapeutic approaches. Front Biomed Technol, 2018. 23(12): p. 2204–2226.

10. Saw, C.L., et al., Current Practice of Therapeutic Hypothermia for Mild Hypoxic Ischaemic Encephalopathy. J Child Neurol, 2019. 34(7): p. 402–409.

11. Davies, A., et al., Can we further optimize therapeutic hypothermia for hypoxic-ischemic encephalopathy? Neural Regen Res, 2019. 14(10): p. 1678–1683.

12. Wagenaar, N., Nikboer, C.H., and van Bel, F., Repair of neonatal brain injury: bringing stem cell-based therapy into clinical practice. Dev Med Child Neurol, 2017. 59(10): p. 997–1003.

13. Cho, K.H.T., et al., Emerging Roles of miRNAs in Brain Development and Perinatal Brain Injury. Front Physiol, 2019. 10: p. 227.

14. Razak, A., and Hussain, A., Erythropoietin in perinatal hypoxic-ischemic encephalopathy: a systematic review and meta-analysis. J Perinat Med, 2019. 47(4): p. 478–489.

15. Alonso-Alconada, D., et al., Neuroprotective Effect of Melatonin: A Novel Therapy against Perinatal Hypoxic-Ischaemia. Int J Mol Sci, 2013. 14(5): p. 9379–9395

16. Zhou, L., et al., Umbilical Cord Blood and Cord Tissue-Derived Cell Therapies for Neonatal Morbidities: Current Status and Future Challenges. Stem Cells Transl Med, 2022. 11(2): p. 135–145.

17. Xi, Y., et al., Human umbilical cord blood mononuclear cells transplantation for perinatal brain injury. Stem Cell Res Ther, 2022. 13: p. 458.

18. Tsuji, M., et al., Autologous cord blood cell therapy for neonatal hypoxic-ischaemic encephalopathy: a pilot study for feasibility and safety. Sci Rep, 2020. 10: p. 4603.

19. Tsuji, M., Sizonenko, S.V., and Baud, O., Editorial: Preventing Developmental Brain Injury – From Animal Models to Clinical Trials. Front. Neurol., 2019. 10: p. 775.

20. Hooijmans, C.R., et al., Facilitating healthcare decisions by assessing the certainty in the evidence from preclinical animal studies. PLoS One, 2018. 13(1): p. e0187271.

21. Deeks, J.J., Higgins, J.P.T., and Altman, D.G., Chapter 10: Analysing data and undertaking meta-analyses. In: Higgins, J.P.T., et al., Cochrane Handbook for Systematic Reviews of Interventions version 6.3 (updated February 2022). Cochrane, 2022. Available from: www.training.cochrane.org/handbook.

22. Richardson, M., Garner, P., and Donegan, S., Interpretation of subgroup analyses in systematic reviews: A tutorial. Clin Epidemiology Glob Health, 2019. 7: p. 192–198.

23. Hooijmans, C.R., et al., SYRCLE’s risk of bias tool for animal studies. BMC Med Res Methodol, 2014. 14(1): p. 43–43.

24. Ahn, S.Y., et al., Mesenchymal Stem Cells Prevent Hydrocephalus After Severe Intraventricular Hemorrhage. Stroke, 2013. 44(2): p. 497–504.

25. Ahn, S.Y., et al., Optimal Route for Mesenchymal Stem Cells Transplantation after Severe Intraventricular Hemorrhage in Newborn Rats. PLoS One, 2015. 10(7): p. e0132919–e0132919.

26. Ahn, S.Y., et al., Mesenchymal stem cells transplantation attenuates brain injury and enhances bacterial clearance in Escherichia coli meningitis in newborn rats. Pediatr Res, 2018. 84(5): p. 778–785.

27. Ahn, S.Y., et al., Stem cell restores thalamocortical plasticity to rescue cognitive deficit in neonatal intraventricular hemorrhage. Exp Neurol, 2021. 342: p. 113736–113736.

28. Aridas, J.D.S., et al., Cord blood mononuclear cells prevent neuronal apoptosis in response to perinatal asphyxia in the newborn lamb: Umbilical cord blood cells for treatment of perinatal asphyxia. The Journal of physiology, 2016. 594(5): p. 1421–1435.

29. Baba, N., et al., Induction of regional chemokine expression in response to human umbilical cord blood cell infusion in the neonatal mouse ischemia-reperfusion brain injury model. PloS one, 2019. 14(9): p. e0221111–e0221111.

30. Bae, S.-H., et al., Long-Lasting Paracrine Effects of Human Cord Blood Cells on Damaged Neocortex in an Animal Model of Cerebral Palsy. Cell Transplant, 2012. 21(11): p. 2497–2515.

31. Chang, Y., et al., Umbilical cord blood CD34+ cells administration improved neurobehavioral status and alleviated brain injury in a mouse model of cerebral palsy. Childs Nerv Syst, 2021. 37(7): p. 2197–2205.

32. Cho, K.H., et al., Therapeutic mechanism of cord blood mononuclear cells via the IL-8-mediated angiogenic pathway in neonatal hypoxic-ischaemic brain injury. Sci Rep, 2020. 10(1): p. 4446–4446.

33. Choi, J.I., et al., Synergistic effect in neurological recovery via anti-apoptotic akt signaling in umbilical cord blood and erythropoietin combination therapy for neonatal hypoxic-ischemic brain injury. Int J Mol Sci, 2021. 22(21): p. 11995.

34. Dalous, J., et al., Use of Human Umbilical Cord Blood Mononuclear Cells to Prevent Perinatal Brain Injury: A Preclinical Study. Stem Cells Dev, 2013. 22(1): p. 169–179.

35. De Paula, S., et al., Hemispheric Brain Injury and Behavioral Deficits Induced by Severe Neonatal Hypoxia-Ischemia in Rats Are Not Attenuated by Intravenous Administration of Human Umbilical Cord Blood Cells. Pediatr Res, 2009. 65(6): p. 631–635.

36. De Paula, S., et al., The dose-response effect of acute intravenous transplantation of human umbilical cord blood cells on brain damage and spatial memory deficits in neonatal hypoxia-ischemia. Neuroscience, 2012. 210: p. 431–441.

37. Drobyshevsky, A., et al., Human Umbilical Cord Blood Cells Ameliorate Motor Deficits in Rabbits in a Cerebral Palsy Model. Dev Neurosci, 2015. 37(4-5): p. 349–362.

38. GeiBler, M., et al., Human umbilical cord blood cells restore brain damage induced changes in rat somatosensory cortex. PLoS One, 2011. 6(6): p. e20194–e20194.

39. Ghaffaripour, H.A., et al., Neuronal cell reconstruction with umbilical cord blood cells in the brain hypoxia-ischemia. Iran Biomed J, 2015. 19(1): p. 29–34.

40. Grandvuillemin, I., et al., Long□Term Recovery After Endothelial Colony□Forming Cells or Human Umbilical Cord Blood Cells Administration in a Rat Model of Neonatal Hypoxic□Ischemic Encephalopathy. Stem Cells Transl Med, 2017. 6(11): p. 1987–1996.

41. Greggio, S., et al., Intra-arterial transplantation of human umbilical cord blood mononuclear cells in neonatal hypoxic–ischemic rats. Life Sci, 2014. 96(1-2): p. 33–39.

42. Hattori, T., et al., Administration of Umbilical Cord Blood Cells Transiently Decreased Hypoxic-Ischemic Brain Injury in Neonatal Rats. Dev Neurosci, 2015. 37(2): p. 95–104.

43. Kadam, S.D., et al., Systemic Injection of CD34+-Enriched Human Cord Blood Cells Modulates Poststroke Neural and Glial Response in a Sex-Dependent Manner in CD1 Mice. Stem Cells Dev, 2015. 24(1): p. 51–66.

44. Kidani, Y., et al., The therapeutic effect of CD133+ cells derived from human umbilical cord blood on neonatal mouse hypoxic-ischemic encephalopathy model. Life Sci, 2016. 157: p. 108–115.

45. Kim, E.S., et al., Human umbilical cord blood-derived mesenchymal stem cell transplantation attenuates severe brain injury by permanent middle cerebral artery occlusion in newborn rats. Pediatr Res, 2012. 72(3): p. 277–284.

46. Kim, Y.E., et al., Intratracheal transplantation of mesenchymal stem cells simultaneously attenuates both lung and brain injuries in hyperoxic newborn rats. Pediatr Res, 2016. 80(3): p. 415–424.

47. Ko, H.R., et al., Human UCB-MSCs treatment upon intraventricular hemorrhage contributes to attenuate hippocampal neuron loss and circuit damage through BDNF-CREB signaling. Stem Cell Res Ther, 2018. 9(1): p. 326–326.

48. Li, X., Q. Shang, and L. Zhang, Comparison of the Efficacy of Cord Blood Mononuclear Cells (MNCs) and CD34+ Cells for the Treatment of Neonatal Mice with Cerebral Palsy. Cell Biochem Biophys, 2014. 70(3): p. 1539–1544.

49. Li, J., et al., Preterm white matter brain injury is prevented by early administration of umbilical cord blood cells. Exp Neurol, 2016. 283: p. 179–187.

50. Li, J., et al., Term vs. preterm cord blood cells for the prevention of preterm brain injury. Pediatr Res, 2017. 82(6): p. 1030–1038.

51. Li, J., et al., Preterm umbilical cord blood derived mesenchymal stem/stromal cells protect preterm white matter brain development against hypoxia-ischemia. Exp Neurol, 2018. 308: p. 120–131.

52. Lyu, H., et al., Umbilical Cord Blood Mononuclear Cell Treatment for Neonatal Rats With Hypoxic Ischemia. Front Cell Neurosci, 2022. 16: p. 823320–823320.

53. Malhotra, A., et al., Neurovascular effects of umbilical cord blood-derived stem cells in growth-restricted newborn lambs. Stem cell research & therapy, 2020. 11(1): p. 1–14.

54. McDonald, C.A., et al., Effects of umbilical cord blood cells, and subtypes, to reduce neuroinflammation following perinatal hypoxic-ischemic brain injury. J Neuroinflammation, 2018. 15(1): p. 1–14.

55. Meier, C., et al., Spastic paresis after perinatal brain damage in rats is reduced by human cord blood mononuclear cells. Pediatr Res, 2006. 59(2): p. 244–249.

56. Nakanishi, K., et al., Rat umbilical cord blood cells attenuate hypoxic-ischemic brain injury in neonatal rats. Sci Rep, 2017. 7(1): p. 44111–44111.

57. Ohshima, M., et al., Evaluations of Intravenous Administration of CD34+ Human Umbilical Cord Blood Cells in a Mouse Model of Neonatal Hypoxic-Ischemic Encephalopathy. Dev Neurosci, 2016. 38(5): p. 331–341.

58. Park, W.S., et al., Hypothermia augments neuroprotective activity of mesenchymal stem cells for neonatal hypoxic-ischemic encephalopathy. PLoS One, 2015. 10(3): p. e0120893–e0120893.

59. Park, W.S., et al., Optimal Timing of Mesenchymal Stem Cell Therapy for Neonatal Intraventricular Hemorrhage. Cell Transplant, 2016. 25(6): p. 1131–1144.

60. Paton, Madison C.B., et al., Human Umbilical Cord Blood Therapy Protects Cerebral White Matter from Systemic LPS Exposure in Preterm Fetal Sheep. Dev Neurosci, 2018. 40(3): p. 258–270.

61. Paton, M.C.B., et al., Umbilical cord blood versus mesenchymal stem cells for inflammation-induced preterm brain injury in fetal sheep. Pediatr Res, 2019. 86(2): p. 165–173.

62. Penny, T.R., et al., Human Umbilical Cord Therapy Improves Long-Term Behavioral Outcomes Following Neonatal Hypoxic Ischemic Brain Injury. Front Physiol, 2019. 10: p. 283–283.

63. Penny, T., et al., Multiple Doses of Umbilical Cord Blood Cells Improve Long□Term Perinatal Brain Injury. Stem cells translational medicine, 2020. 9(S1): p. S3–S3.

64. Penny, T.R., et al., Umbilical cord blood therapy modulates neonatal hypoxic ischemic brain injury in both females and males. Scientific reports, 2021. 11(1): p. 15788–15788.

65. Pimentel-Coelho, P.M., et al., Human cord blood transplantation in a neonatal rat model of hypoxic-ischemic brain damage: Functional outcome related to neuroprotection in the striatum. Stem Cells Dev, 2010. 19(3): p. 351–358.

66. Purohit, D., et al., Human Cord Blood Derived Unrestricted Somatic Stem Cells Restore Aquaporin Channel Expression, Reduce Inflammation and Inhibit the Development of Hydrocephalus After Experimentally Induced Perinatal Intraventricular Hemorrhage. Front Cell Neurosci, 2021. 15: p. 633185–633185.

67. Rosenkranz, K., et al., Transplantation of human umbilical cord blood cells mediated beneficial effects on apoptosis, angiogenesis and neuronal survival after hypoxic-ischemic brain injury in rats. Cell Tissue Res, 2012. 348(3): p. 429–438.

68. Rosenkranz, K., et al., Changes in Interleukin-1 alpha serum levels after transplantation of umbilical cord blood cells in a model of perinatal hypoxic-ischemic brain damage. Ann Anat, 2013. 195(2): p. 122–127.

69. Tsuji, M., et al., Effects of intravenous administration of umbilical cord blood CD34+ cells in a mouse model of neonatal stroke. Neuroscience, 2014. 263: p. 148–158.

70. Vinukonda, G., et al., Human Cord Blood□Derived Unrestricted Somatic Stem Cell Infusion Improves Neurobehavioral Outcome in a Rabbit Model of Intraventricular Hemorrhage. Stem Cells Transl Med, 2019. 8(11): p. 1157–1169.

71. Wang, X.-L., et al., Umbilical cord blood cells regulate endogenous neural stem cell proliferation via hedgehog signaling in hypoxic ischemic neonatal rats. Brain Res, 2013. 1518: p. 26–35.

72. Wang, X., Y. Zhao, and X. Wang, Umbilical cord blood cells regulate the differentiation of endogenous neural stem cells in hypoxic ischemic neonatal rats via the hedgehog signaling pathway. Brain Res, 2014. 1560: p. 18–26.

73. Wasielewski, B., et al., Neuroglial activation and Cx43 expression are reduced upon transplantation of human umbilical cord blood cells after perinatal hypoxic-ischemic injury. Brain Res, 2012. 1487: p. 39–53.

74. Xia, G., et al., Intracerebral transplantation of mesenchymal stem cells derived from human umbilical cord blood alleviates hypoxic ischemic brain injury in rat neonates. Journal of Perinatal Medicine, 2010. 38(2): p. 215–221.

75. Yasuhara, T., et al., Mannitol facilitates neurotrophic factor up□regulation and behavioural recovery in neonatal hypoxic□ischaemic rats with human umbilical cord blood grafts. J Cell Mol Med, 2010. 14(4): p. 914–921.

76. Yu, Y., et al., Effects of human umbilical cord blood CD34+ cell transplantation in neonatal hypoxic-ischemia rat model. Brain Dev, 2019. 41(2): p. 173–181.

77. Zhang, J., et al., Umbilical cord mesenchymal stem cells and umbilical cord blood mononuclear cells improve neonatal rat memory after hypoxia-ischemia. Behav Brain Res, 2019. 362: p. 56–63.

78. Zhang, M.-B., et al., Transplantation of umbilical cord blood mononuclear cells attenuates the expression of IL-1β via the TLR4/NF-κB pathway in hypoxic-ischemic neonatal rats. J Neurorestoratology, 2020. 8(2): p. 122–130.

79. Trounson, A., and McDonald, C., Stem Cell Therapies in Clinical Trials: Progress and Challenges. Cell Stem Cell, 2015. 17(1): p. 11–22.

80. Broxmeyer, H.E., Biology of cord blood cells and future prospects for enhances clinical benefit. Cytotherapy, 2005. 7(3): p. 209–218.

81. Hoang, D.M., et al., Stem cell-based therapy for human diseases. Signal Transduct Target Ther, 2022. 7: p. 272.

82. Lehnerer, V., et al., Mesenchymal stem cell therapy in perinatal arterial ischaemic stroke: systematic review of preclinical studies. Pediatr Res, 2022. p. 1–16.

83. Jiao, Y., Xiao-yan, L., and Liu, J., A New Approach to Cerebral Palsy Treatment: Discussion of the Effective Components of Umbilical Cord Blood and its Mechanisms of Action. Cell Transplant, 2019. 28(5): p. 497–509.

84. Ahn, S.Y., et al., Mesenchymal Stem Cell for Sever Intraventricular Hemorrhage in Preterm Infants: Phase 1 Dose-Escalation Clinical Trial. Stem Cells Transl Med, 2018. 7(12): p. 847–856.

85. Eggenberger, S., et al., Stem cell treatment and cerebral palsy: Systematic review and meta-analysis. World J Stem Cells, 2019. 11(10): p. 891–903.

86. Sun, J.M., and Kurtzberg, J., The Effects of Umbilical Cord Blood and Cord Tissue Cell Therapies in Animal and Human Models of Cerebral Palsy. Cerebral Palsy, 2020: p. 97–110.

