## Supplementary files for "Umbilical cord blood-derived cell therapy for perinatal brain injury: A systematic review & meta-analysis of preclinical studies - Part B"

**Supplemental 1.** Search strategy and search terms.

| **#** | **Query** |
| --- | --- |
| 1 | (perinatal or neonat* or newborn or infant or f?et* or prematur*).tw. |
| 2 | exp Animals/ or exp Rats/ or exp Mice/ or exp Sheep/ or exp Swine/ or exp Rabbits/ or exp Disease Models, Animal/ |
| 3 | 1 and 2 |
| 4 | (brain or cerebral or encephalopath* or neuro*).tw. |
| 5 | (injury or damage or insult or trauma or hypox* or anox* or oxygen deprivation or isch?em* or h?emorrhag* or stroke or thrombosis or hypoperfusion or underperfusion or asphyxia or growth restriction or periventricular leukomalacia or inflamm* or chorioamnionitis or infection or sepsis or toxin or hypoglyc?em* or placental insufficiency or placental abnormality or placental lesion or placental abruption or uterine rupture or maternal obesity or metabolic or stress or distress or vascular dysfunction or low birth weight or cord prolapse or eclampsia or preeclampsia).tw. |
| 6 | 4 and 5 |
| 7 | (umbilical cord and cell).tw. |
| 8 | umbilical cord blood.tw. |
| 9 | (cell and cord blood).tw. |
| 10 | (h?ematopoietic and cord blood).tw. |
| 11 | (mesenchymal and cord blood).tw |
| 12 | (stromal and cord blood).tw. |
| 13 | (endothelial progenitor and cord blood).tw. |
| 14 | (mononuclear and cord blood).tw. |
| 15 | (regulatory T and cord blood).tw. |
| 16 | 7 or 8 or 9 or 10 or 11 or 12 or 13 or 14 or 15 |
| 17 | 3 and 6 and 16 |

**Supplemental 2.** Forest plots demonstrating the effect of preterm and term models on brain outcomes of **(A)** Apoptosis – grey matter; **(B)** Astrogliosis- grey matter; **(C)** Astrogliosis – white matter; **(D)** Infarct size; **(E)** Microglial activation – grey matter; **(F)** Oligodendrocyte number – white matter; **(G)** Neuroinflammation - TNF-α; **(H)** Neuroinflammation - IL-1$\beta$. Abbreviations: admin, administration; ECFC, endothelial colony-forming cell; EPC, endothelial progenitor cell; ICV, intracerebroventricular; IV, intraventricular; MNC, mononuclear cell; PCB, preterm cord blood; TCB, term cord blood; Treg, T regulatory cell.

**(A)**

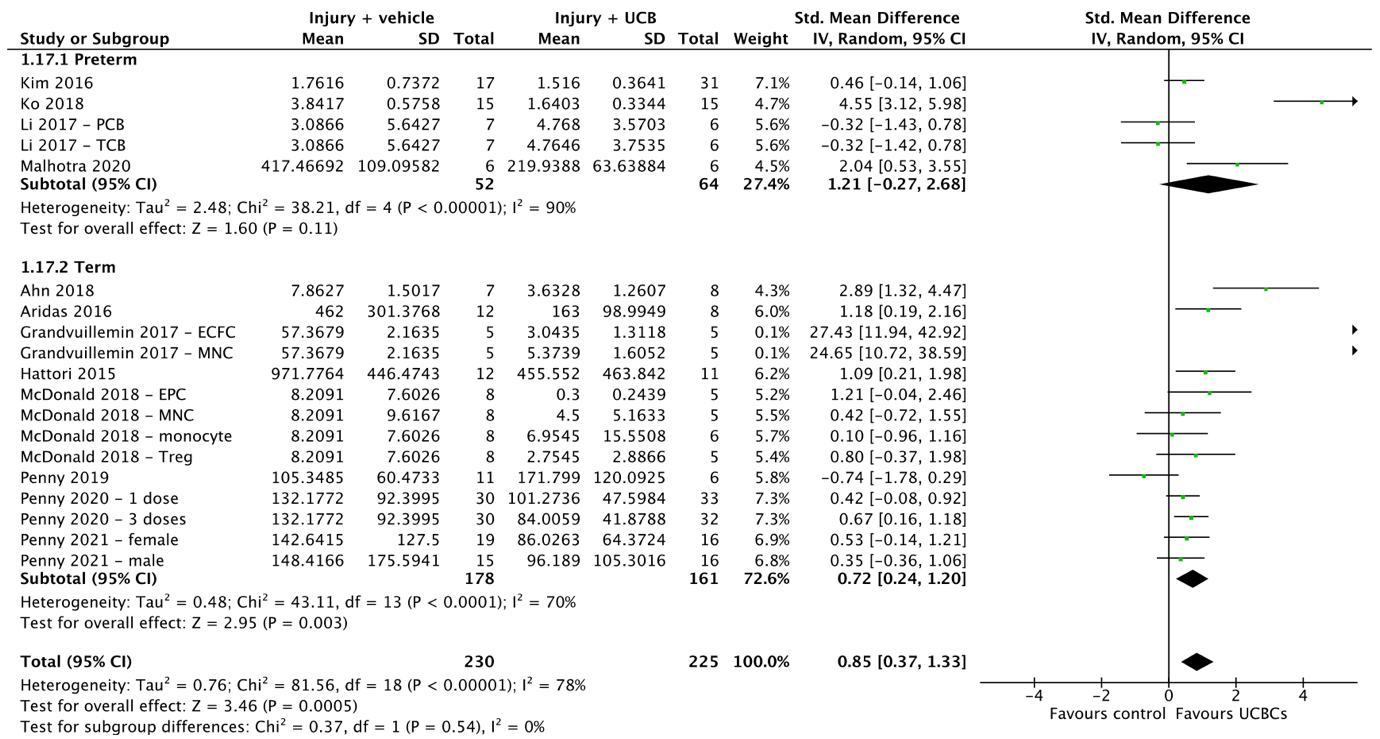

**(B)**

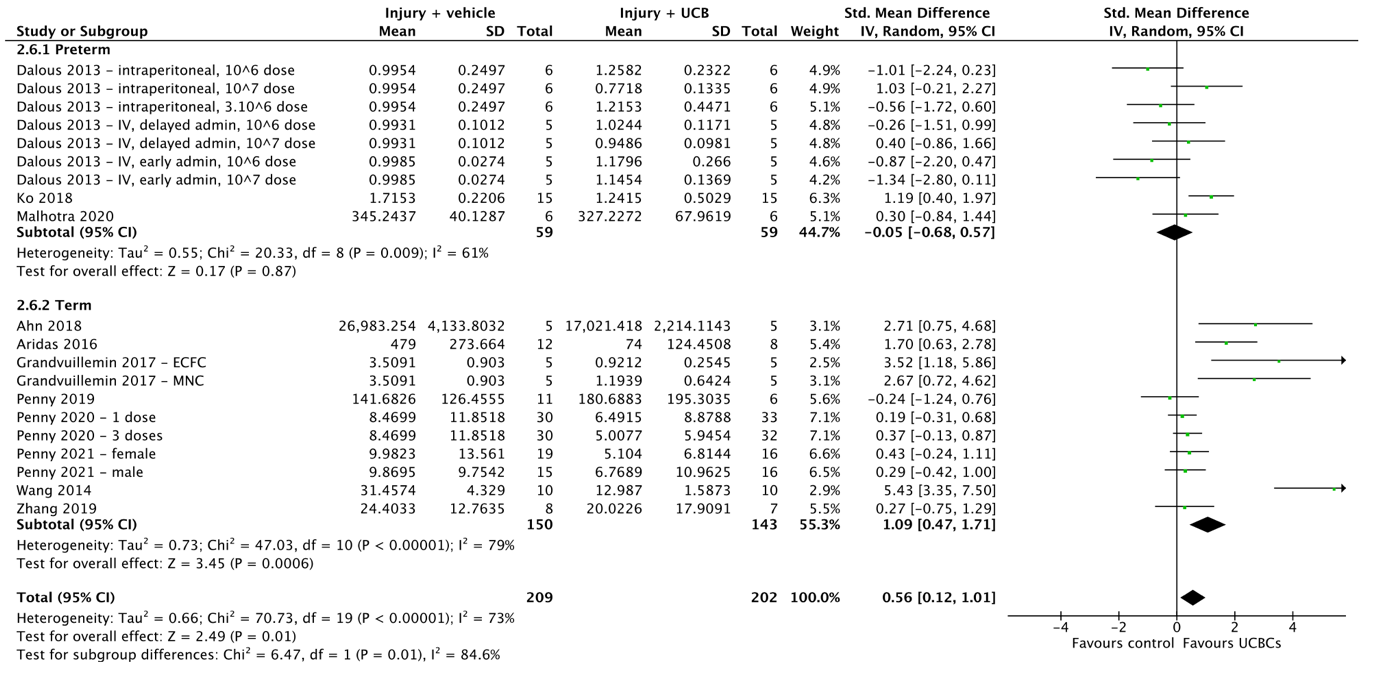

**(C)**

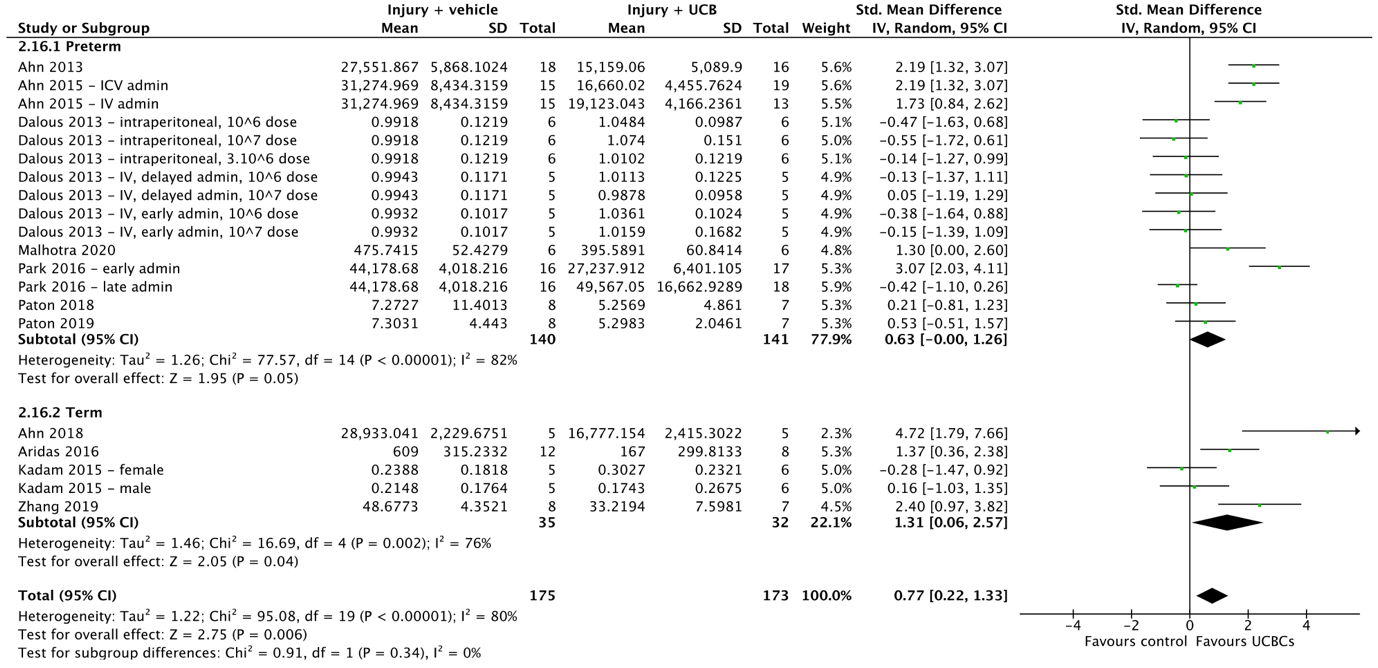

**(D)**

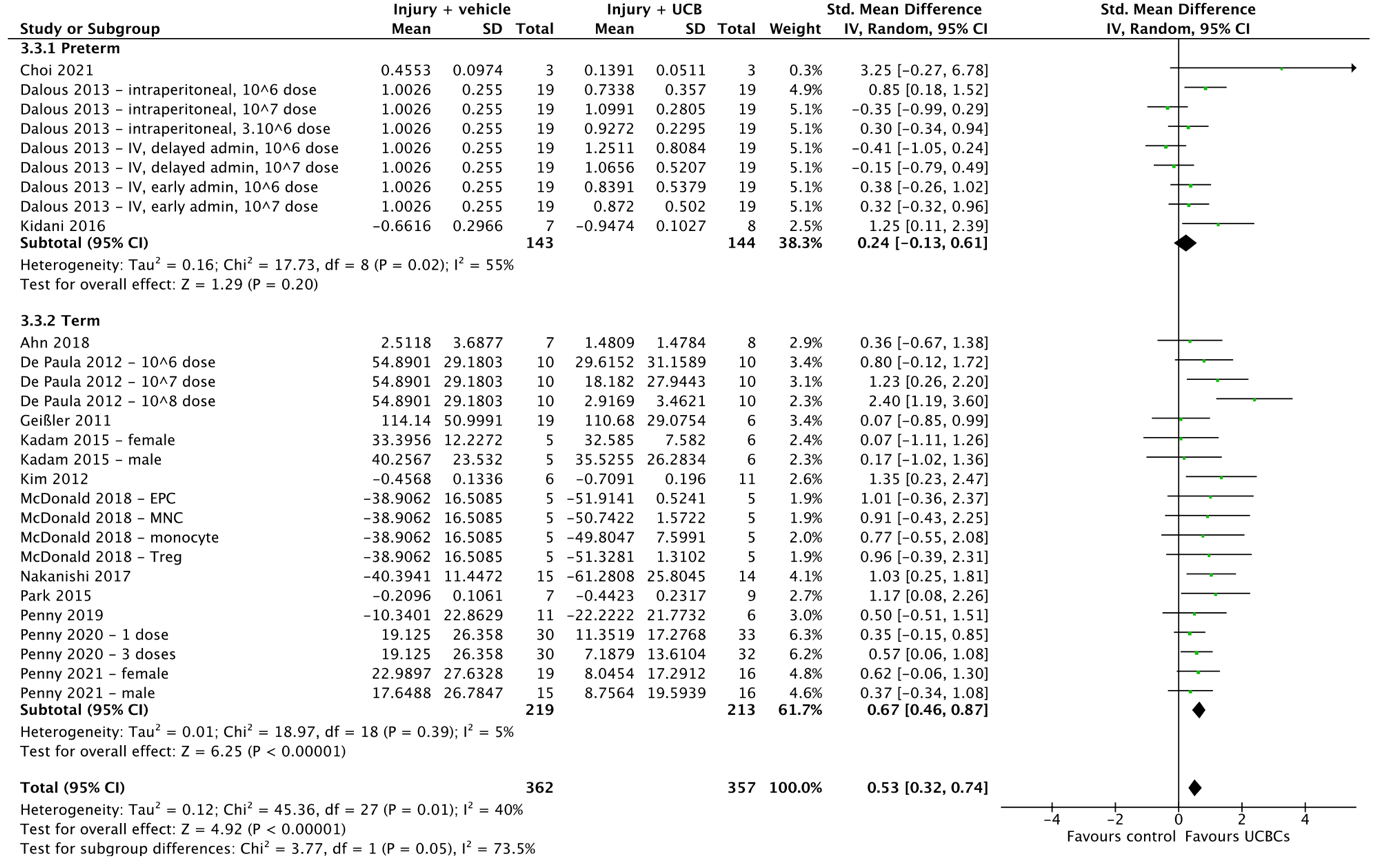

**(E)**

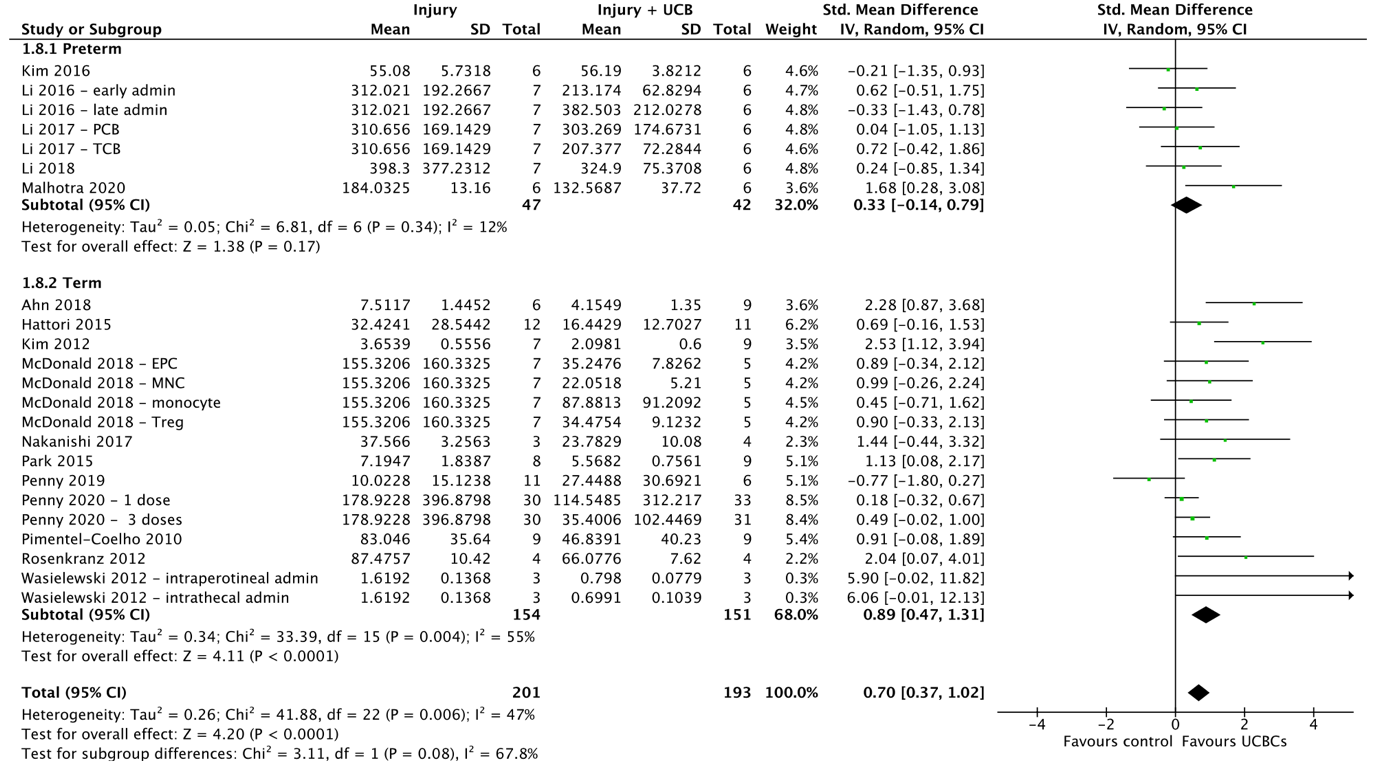

**(F)**

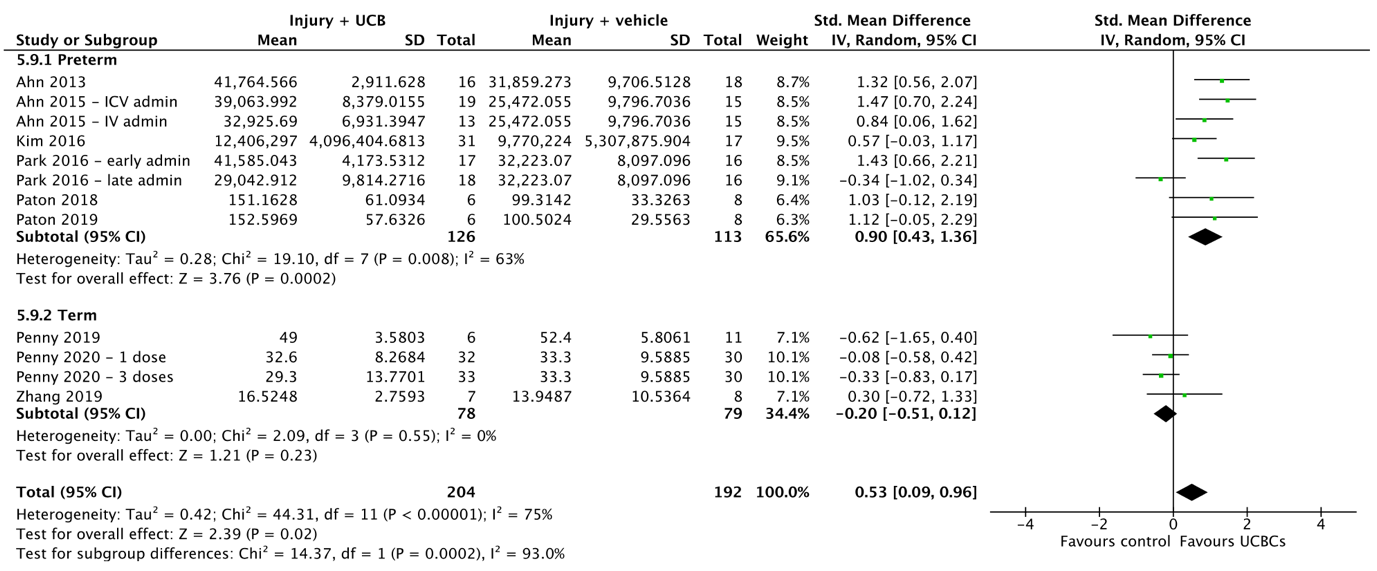

**(G)**

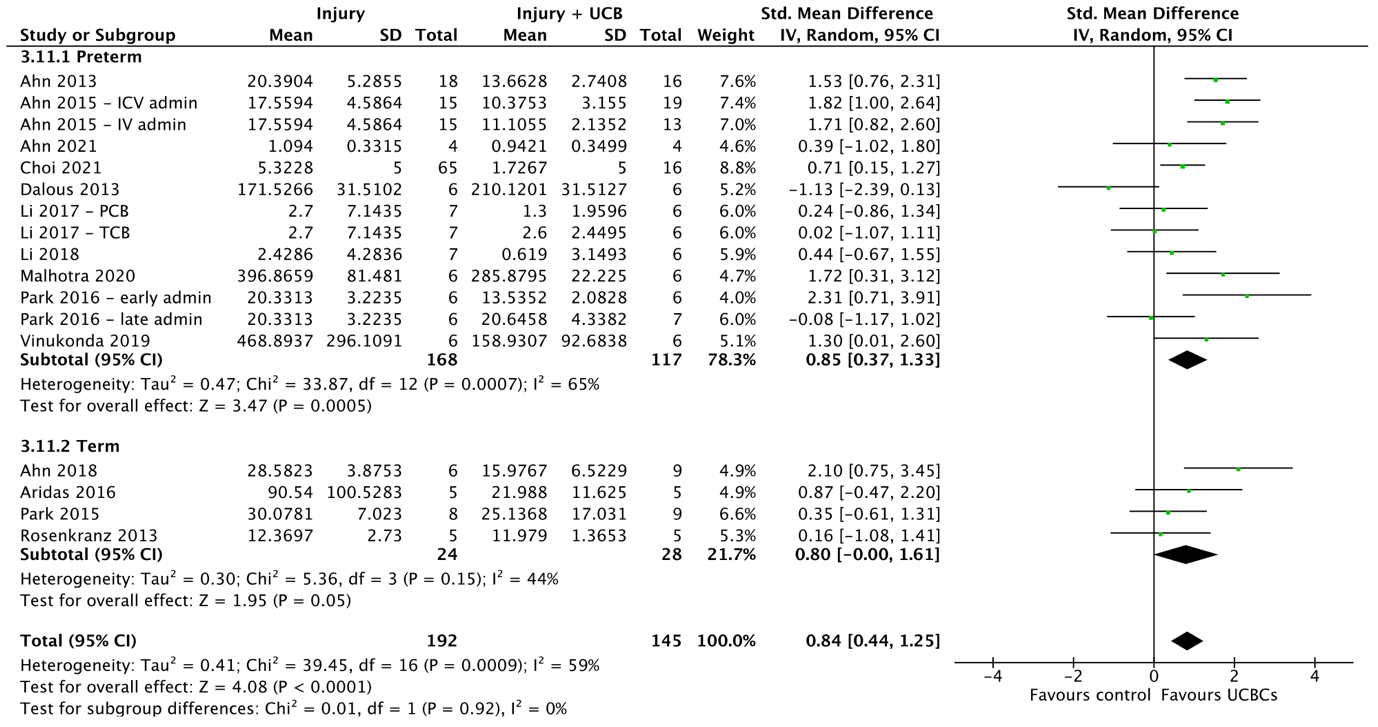

**(H)**

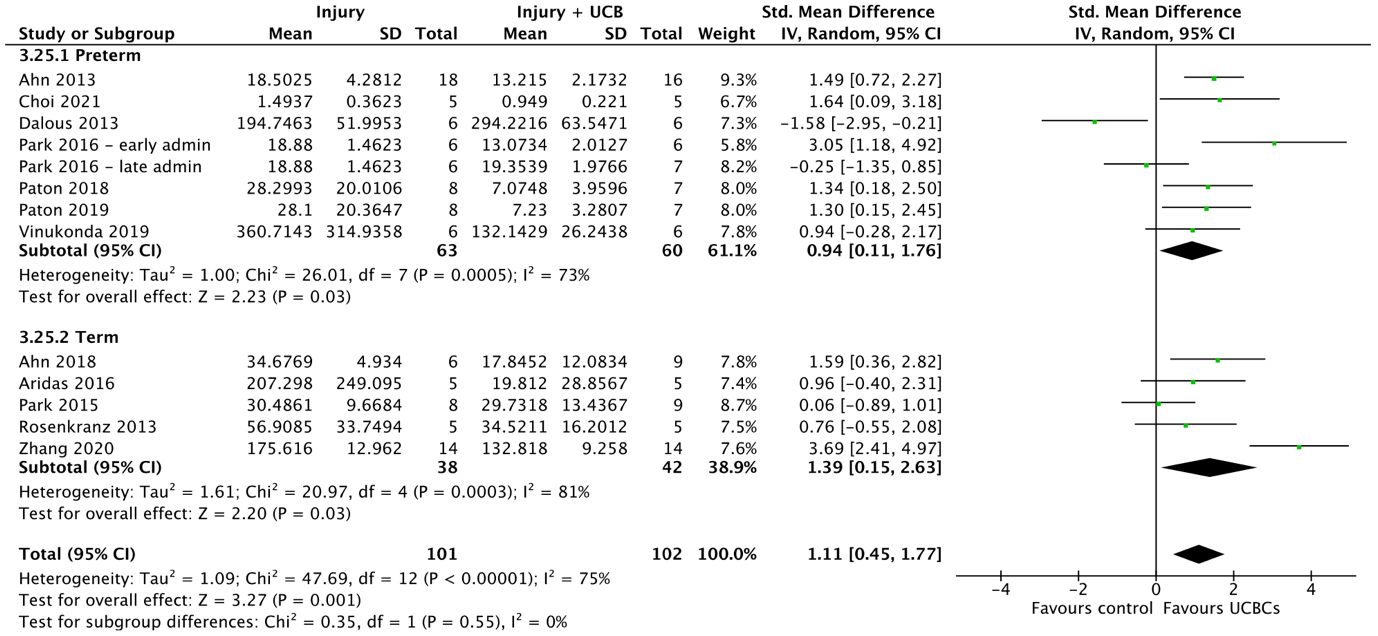

**Supplemental 3.** Forest plots demonstrating the effect of brain injury model on brain outcomes of **(A)** Apoptosis – white matter; **(B)** Neuroinflammation - TNF-α. Abbreviations: admin, administration; HI, hypoxia ischemia; ICV, intracerebroventricular; IV, intraventricular; IVH, intraventricular haemorrhage; PCB, preterm cord blood; TCB, term cord blood.

**(A)**

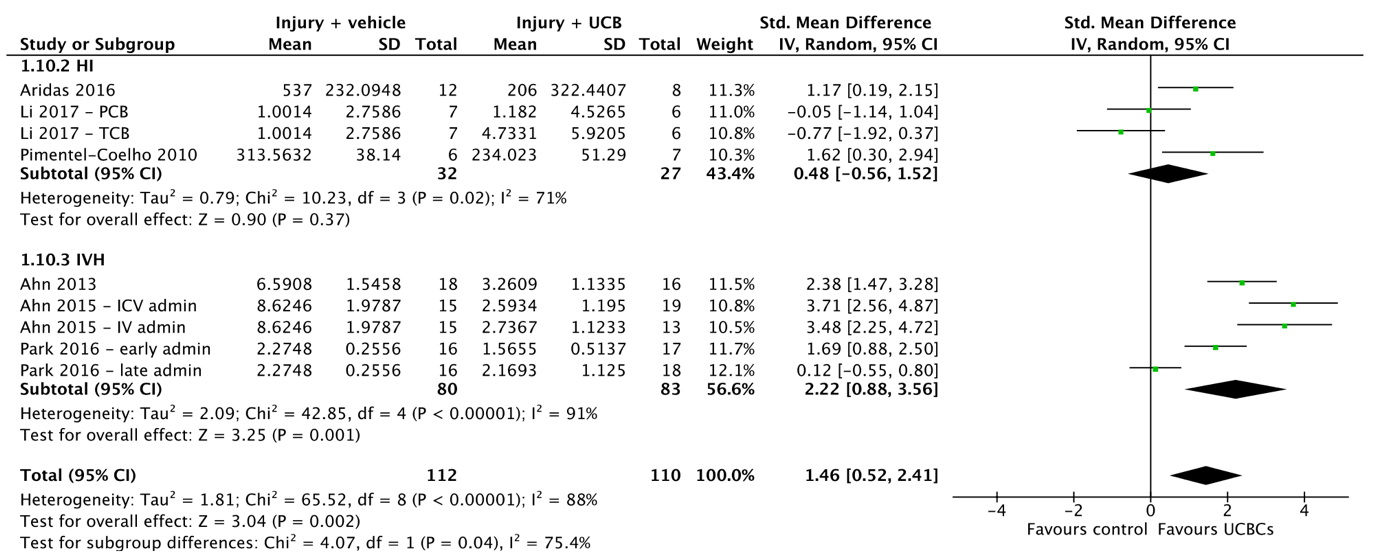

**(B)**

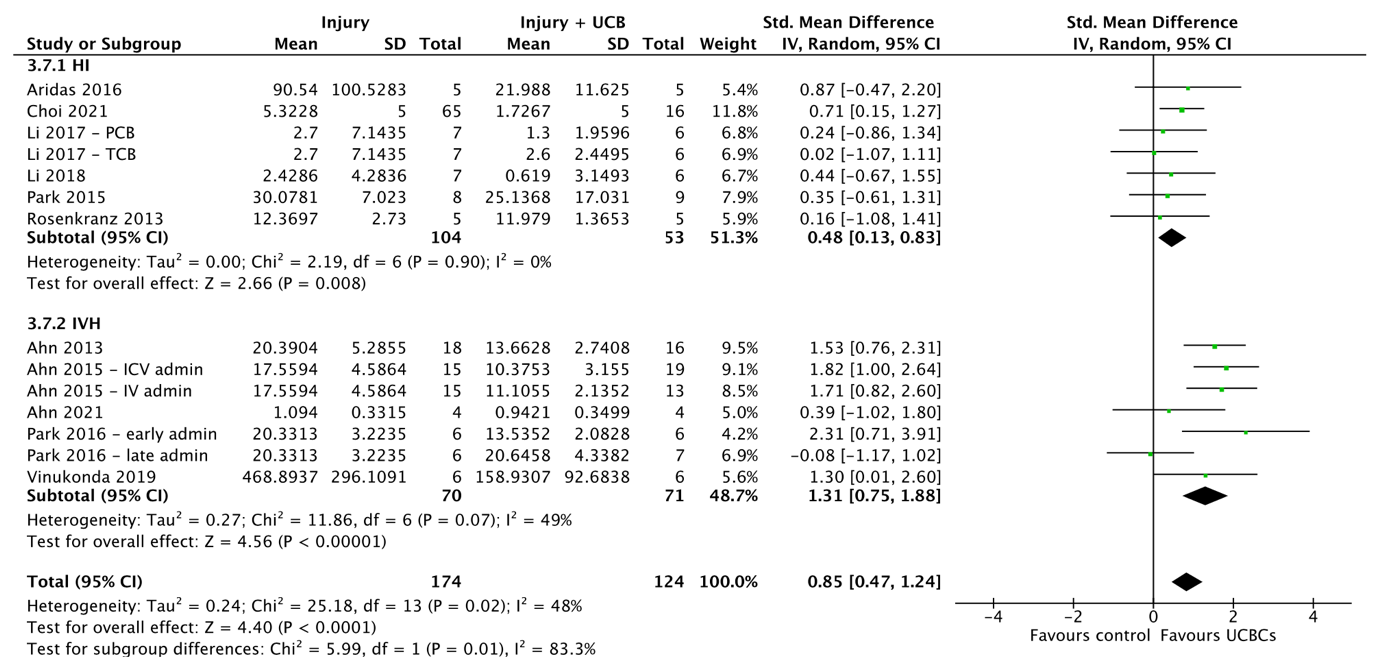

**Supplemental 4**. Forest plots demonstrating the effect of cell type on brain outcomes of **(A)** Apoptosis - grey matter; **(B)** Apoptosis - white matter; **(C)** Astrogliosis - white matter; **(D)** Microglial activation - grey matter; **(E)** Oligodendrocyte number – white matter; **(F)** Neuroinflammation - TNF-α; **(G)** Neuroinflammation - IL-6. **(H)** Neuroinflammation - IL-1$\beta$. Abbreviations: admin, administration; ICV, intracerebroventricular; IV, intraventricular; MNC, mononuclear cell; MSC, mesenchymal stromal cell; PCB, preterm cord blood; TCB, term cord blood.

**(A)**

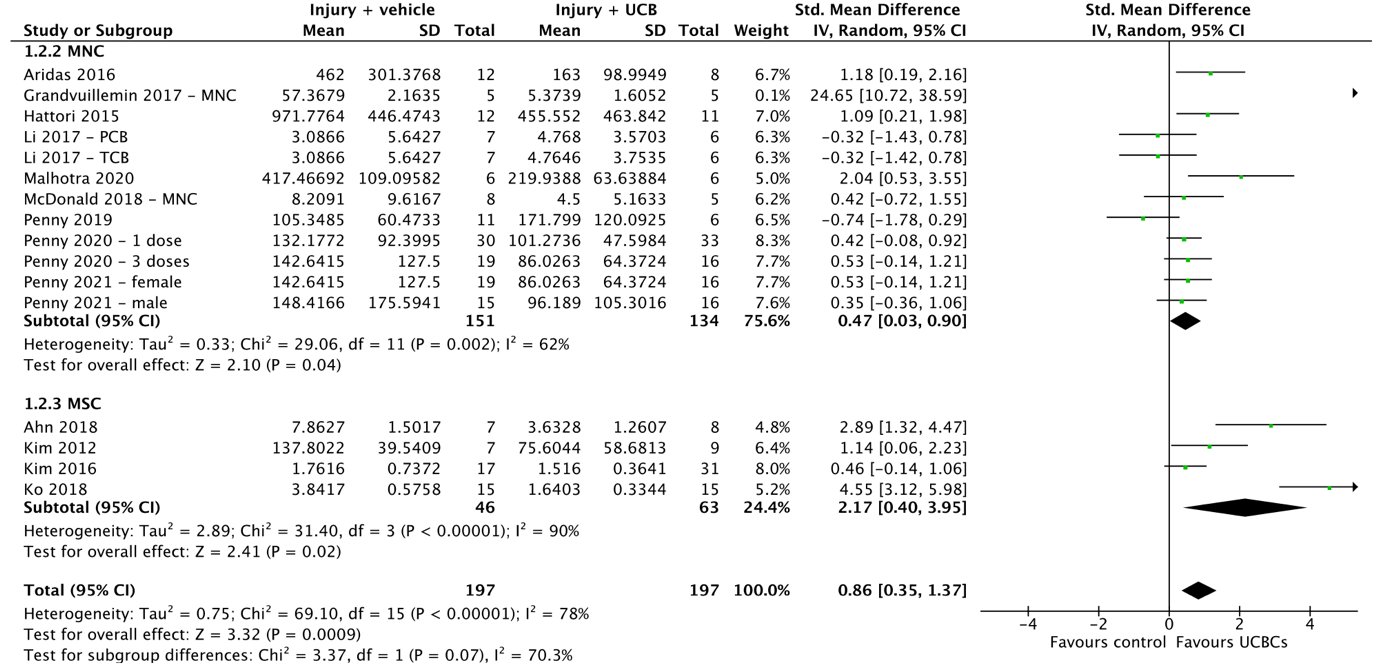

**(B)**

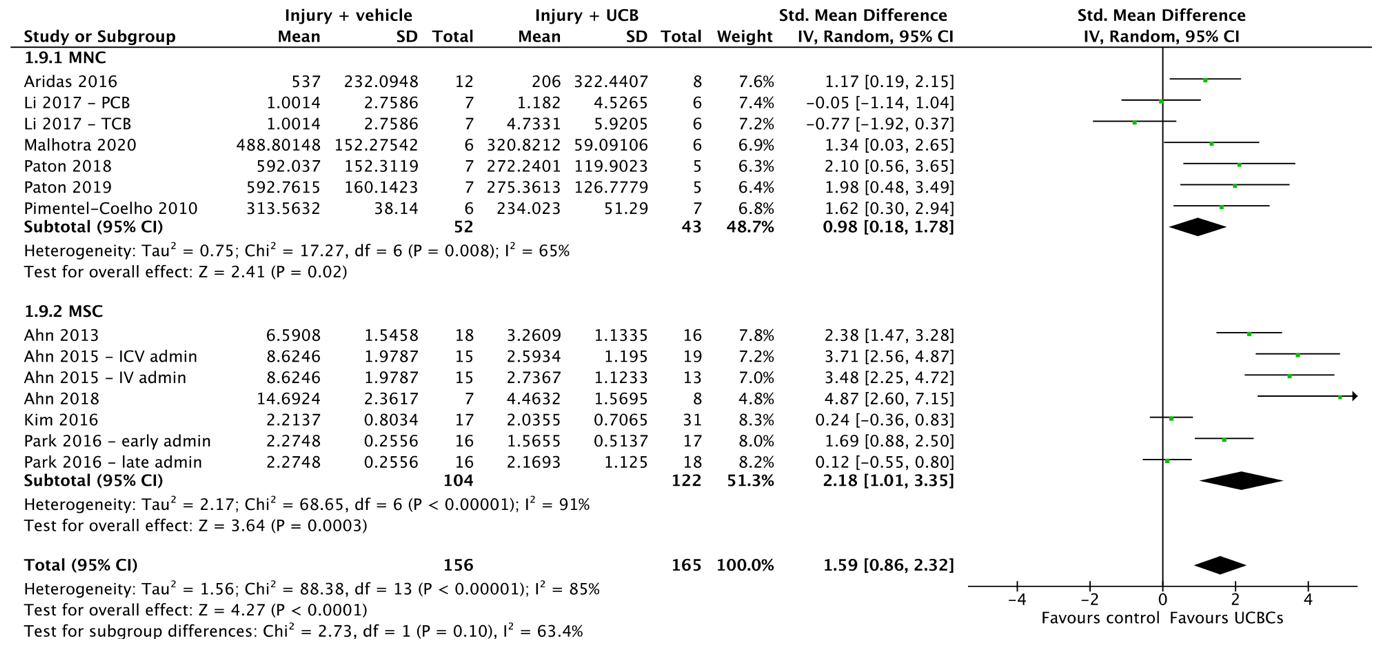

**(C)**

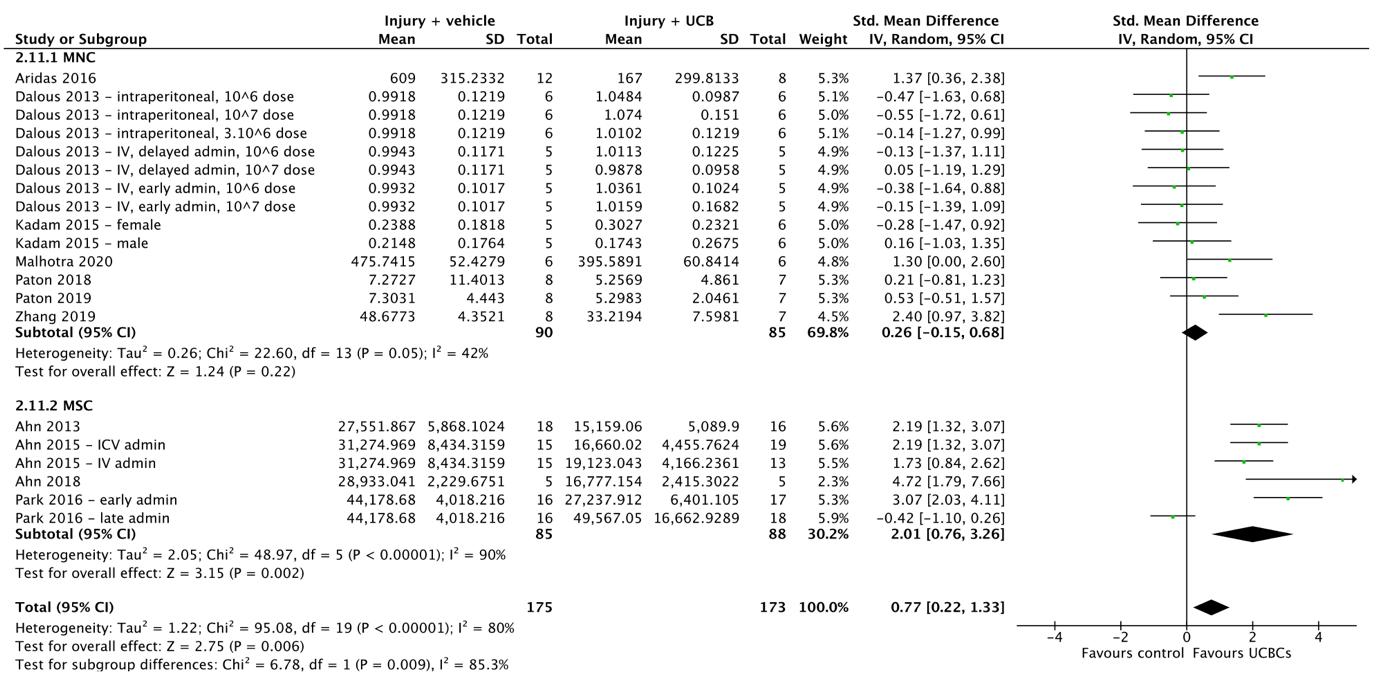

**(D)**

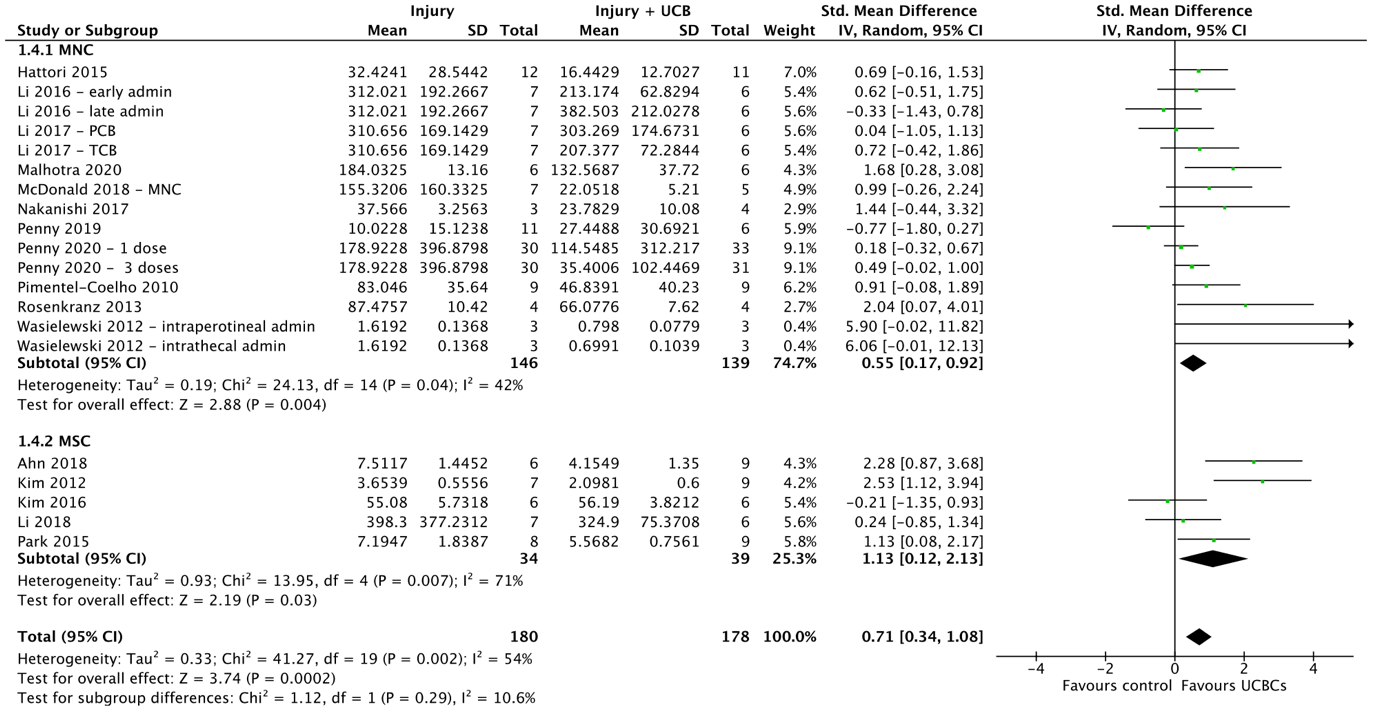

**(E)**

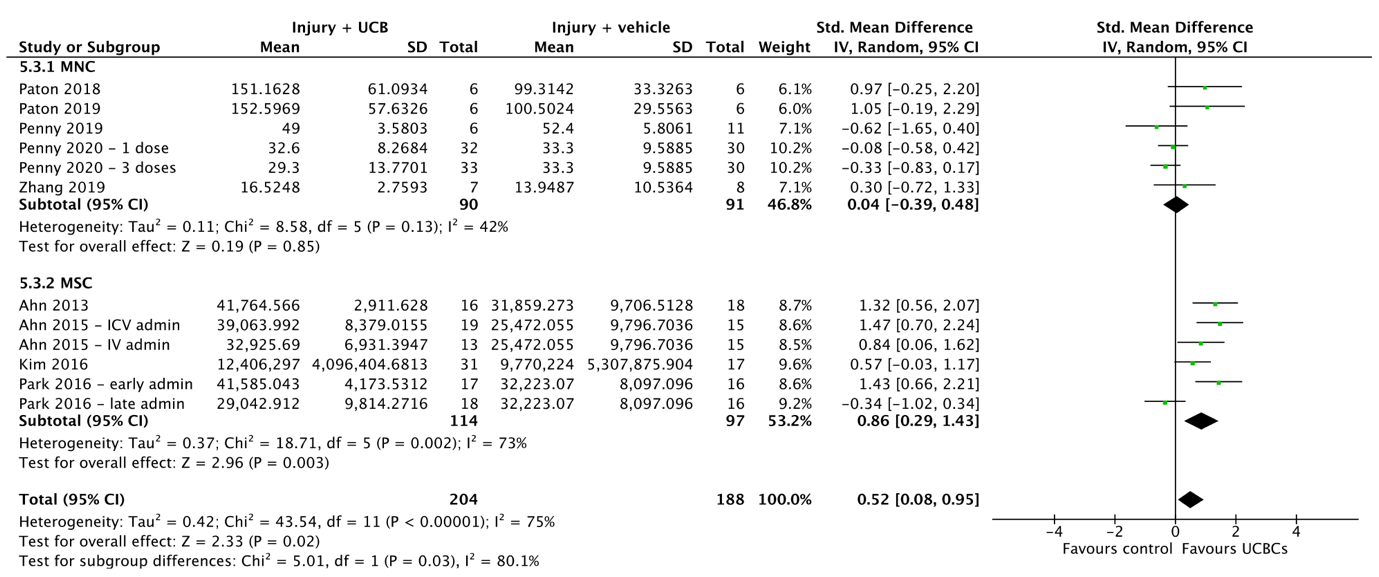

**(F)**

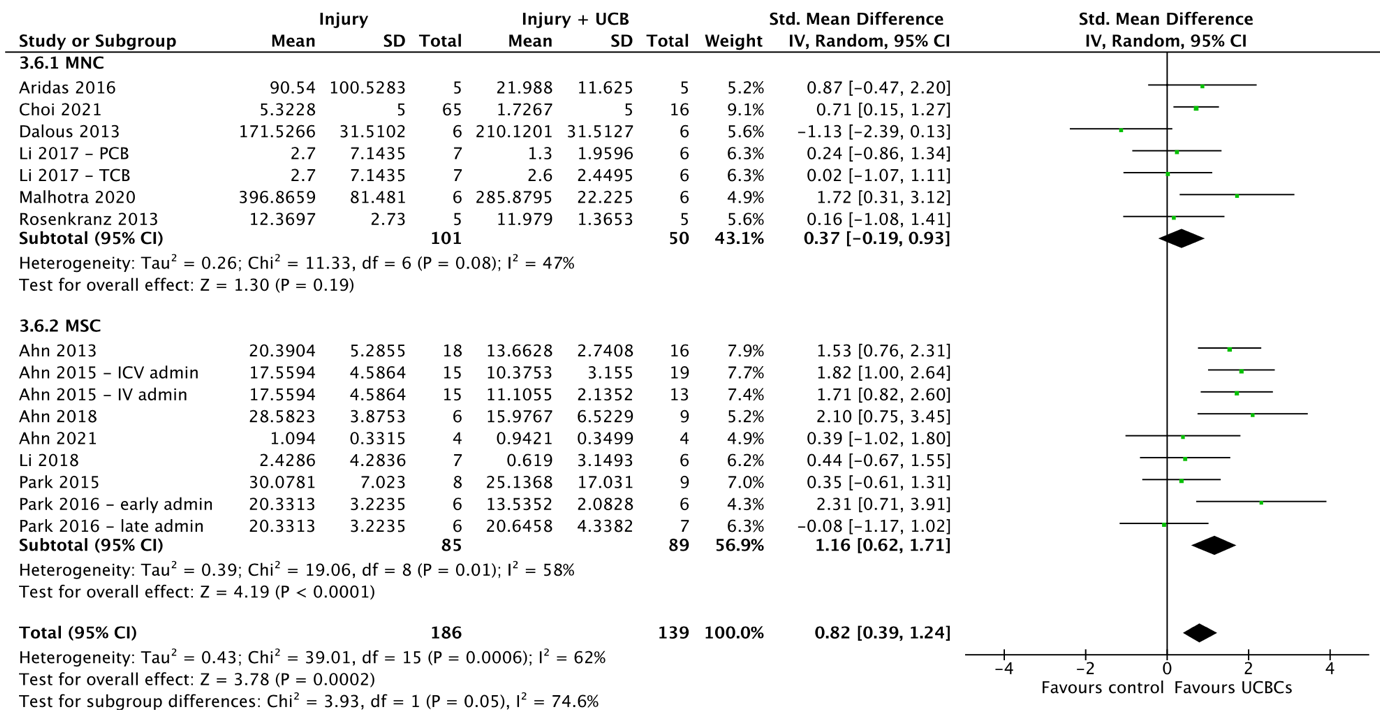

**(G)**

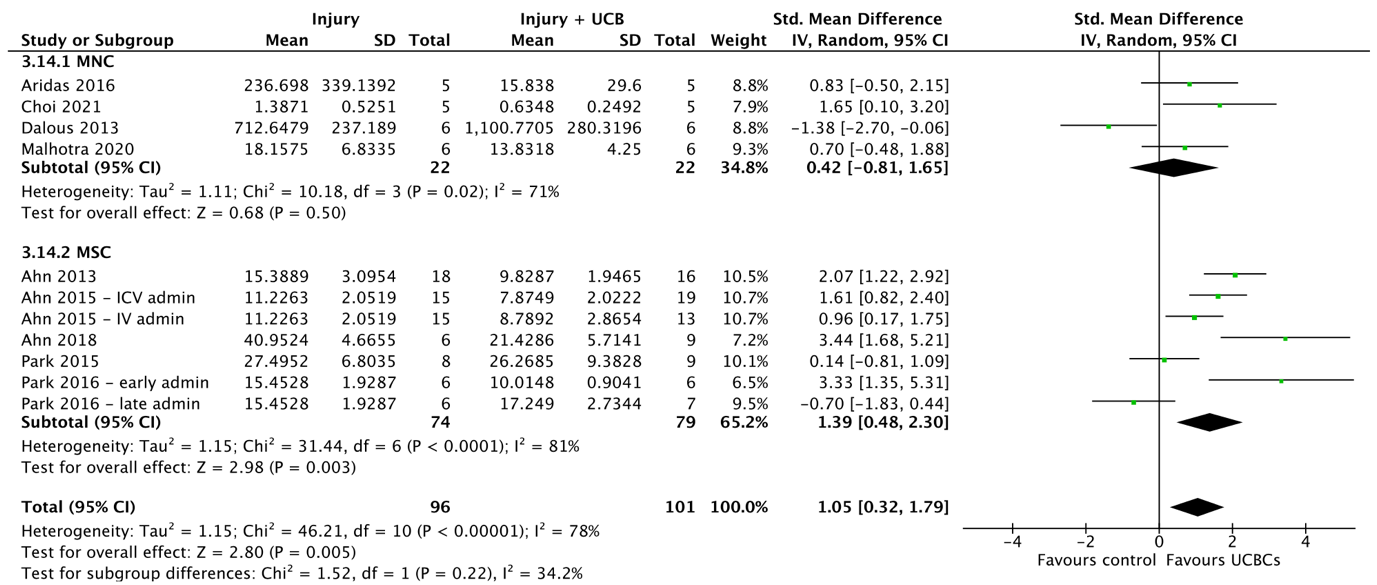

**(H)**
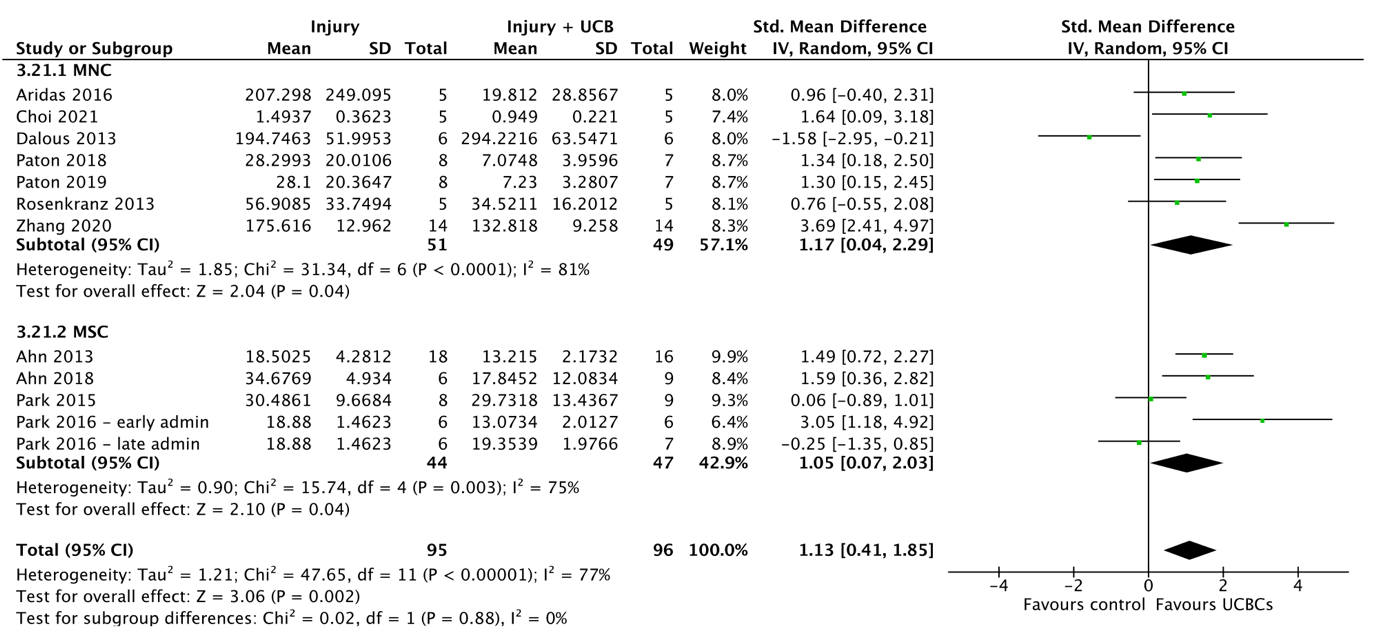

**Supplemental 5**. Forest plots demonstrating the effect of timing of UCB-derived cell therapy post injury induction on brain outcomes of **(A)** Apoptosis - grey matter; **(B)** Apoptosis - white matter; **(C)** Astrogliosis - grey matter; **(D)** Astrogliosis - white matter; **(E)** Infarct size; **(F)** Microglial activation - grey matter; **(G)** Neuroinflammation - TNF-α; **(H)** Neuroinflammation - IL-1$\beta$. Abbreviations: admin, administration; ECFC, endothelial colony-forming cell; EPC, endothelial progenitor cell; ICV, intracerebroventricular; IV, intraventricular; MNC, mononuclear cell; PCB, preterm cord blood; TCB, term cord blood; Treg, T regulatory cell.

**(A)**
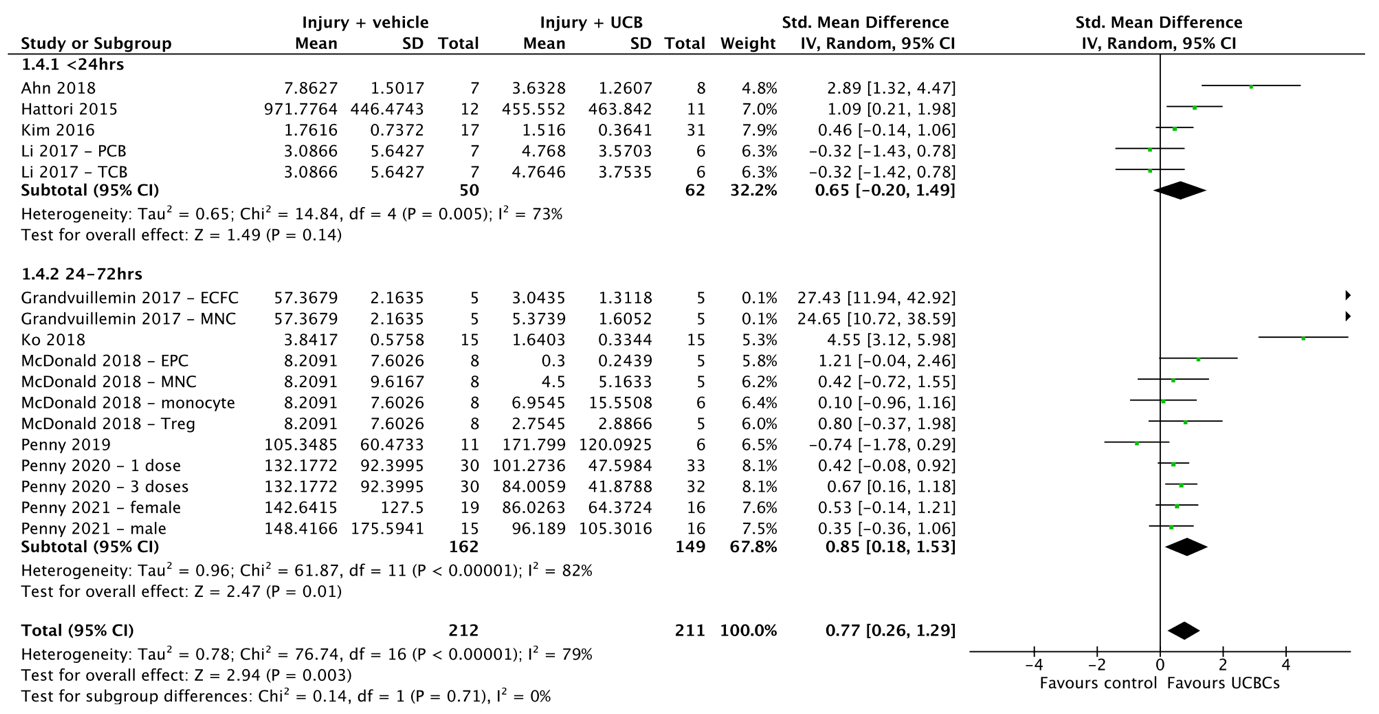

**(B)**
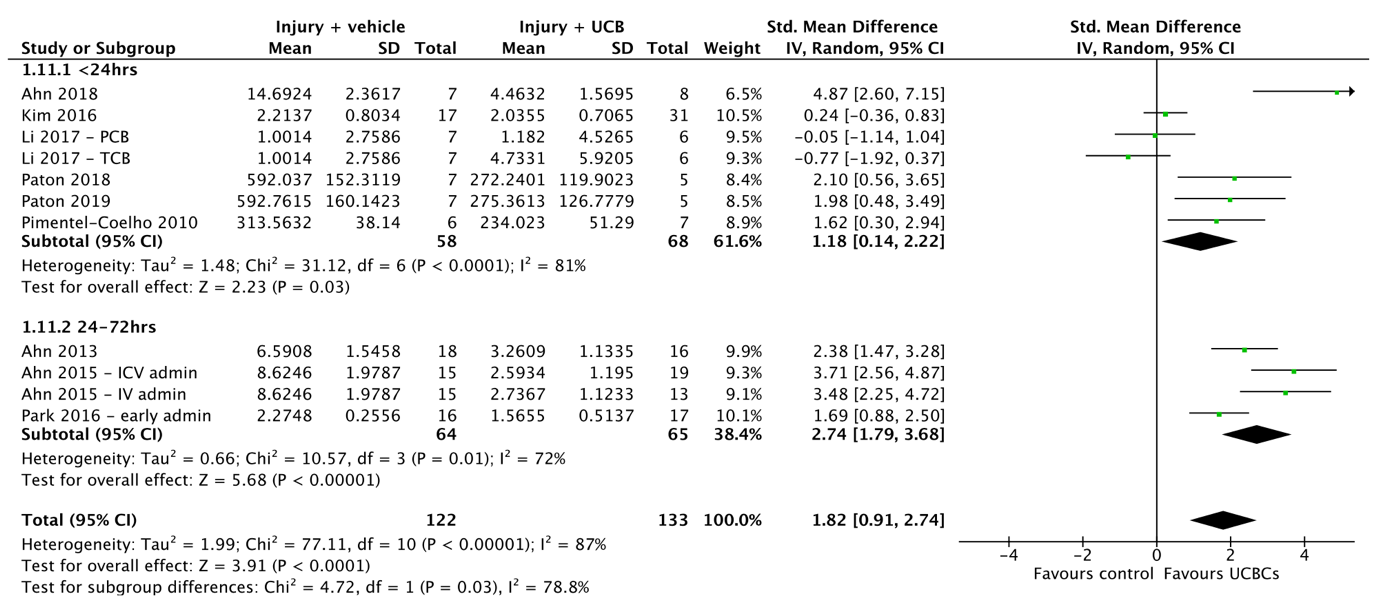

**(C)**
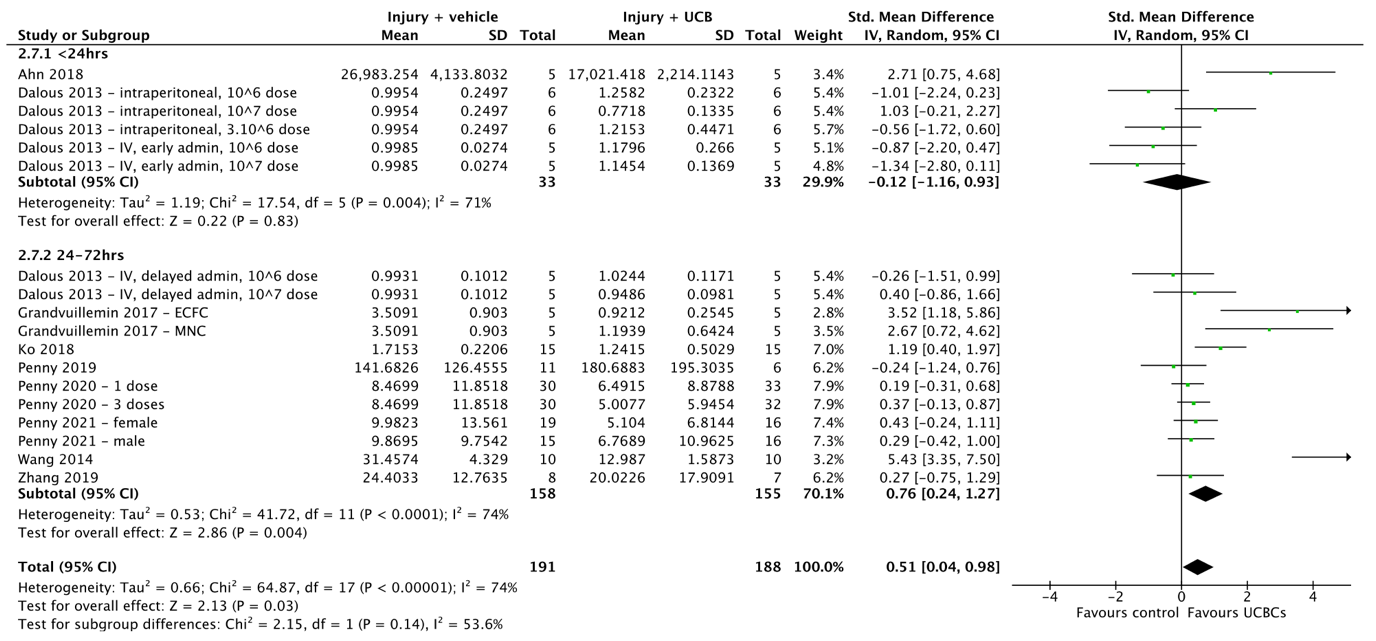

**(D)**
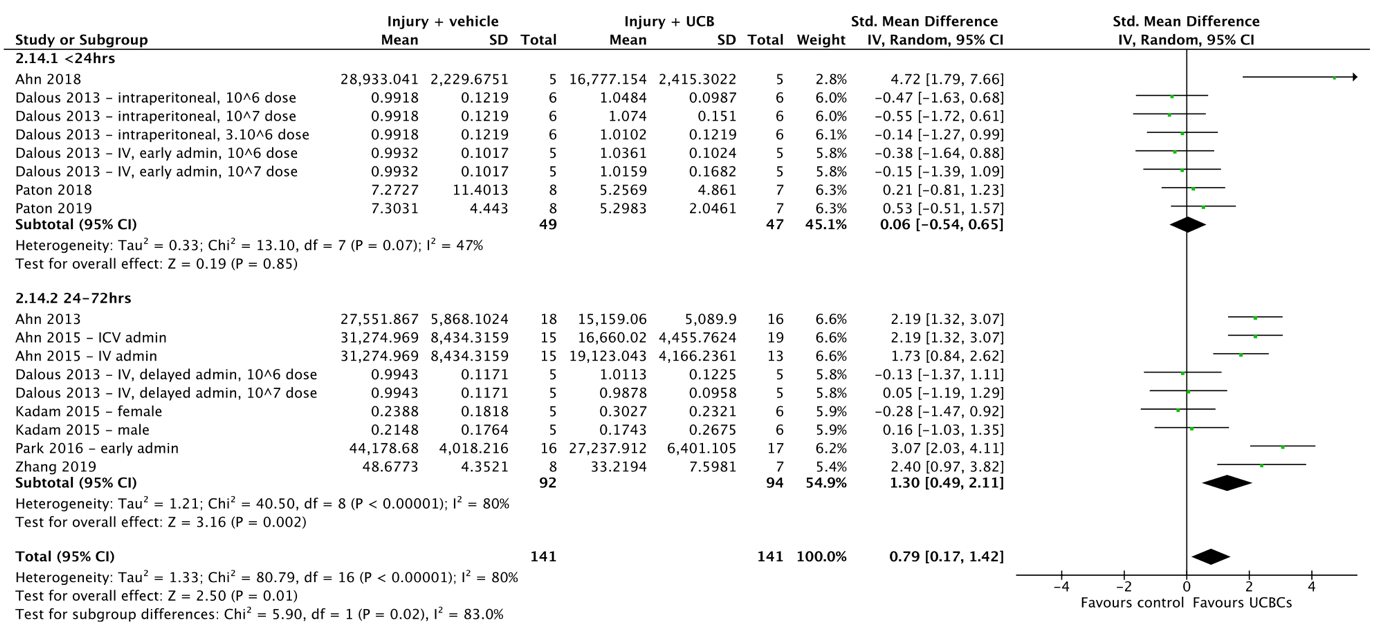

**(E)**

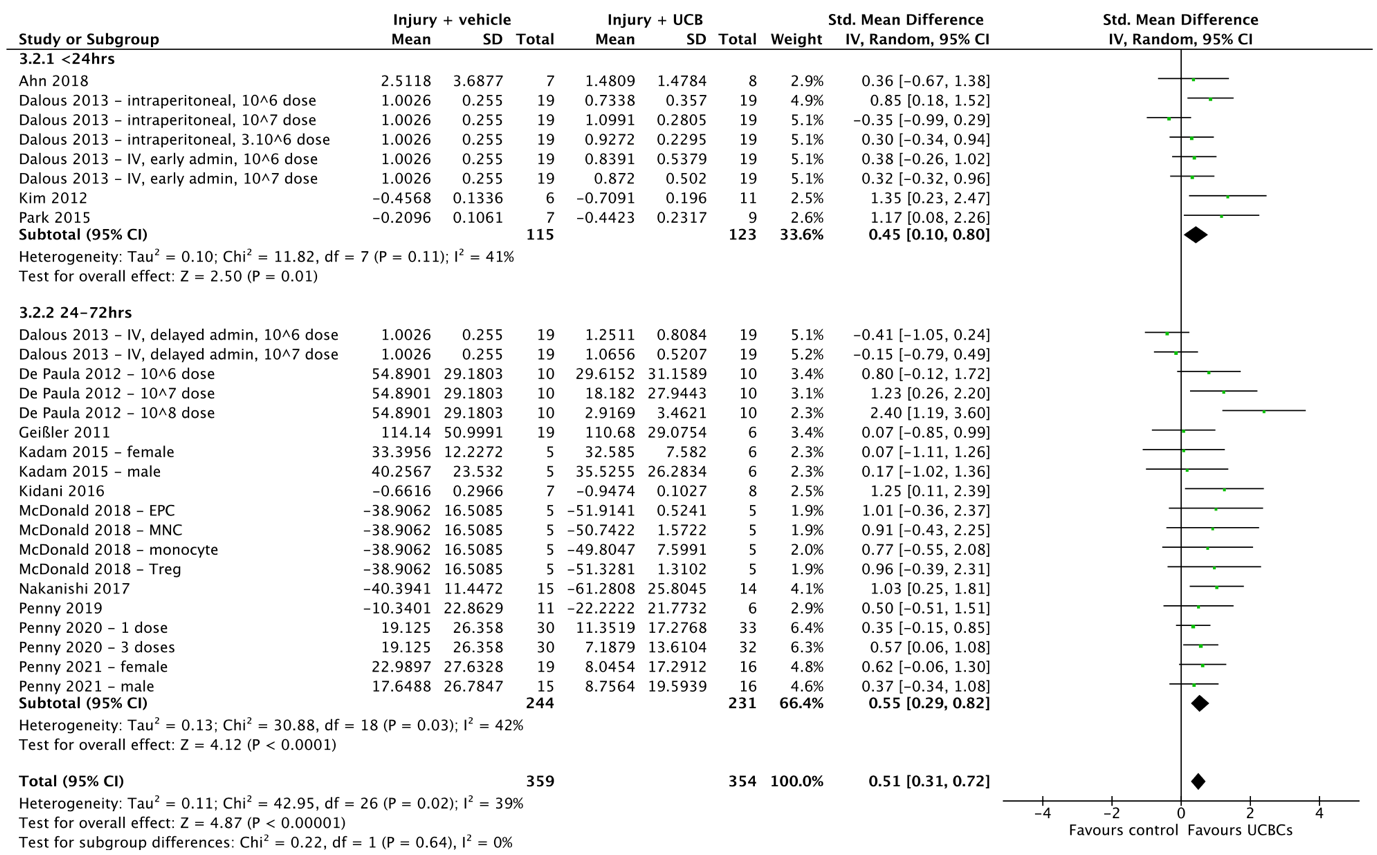

**(F)**

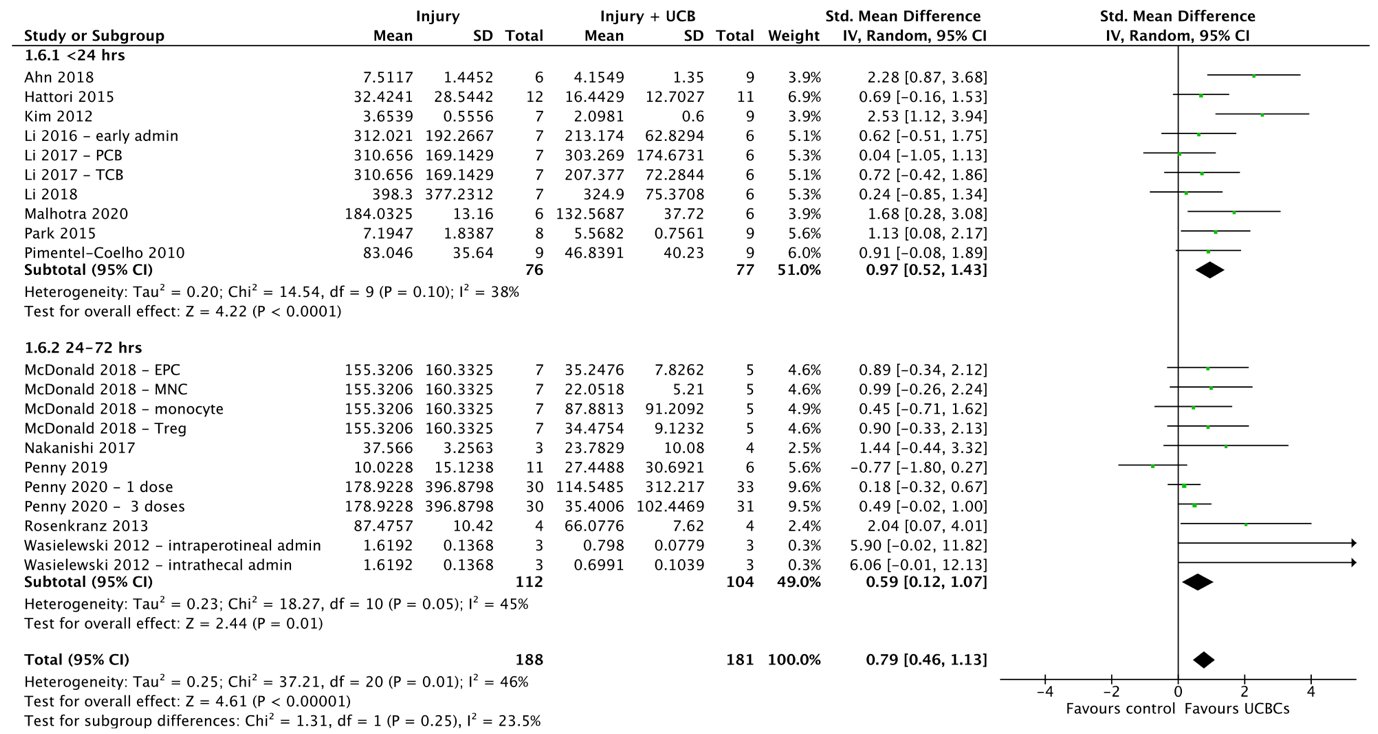

**(G)**
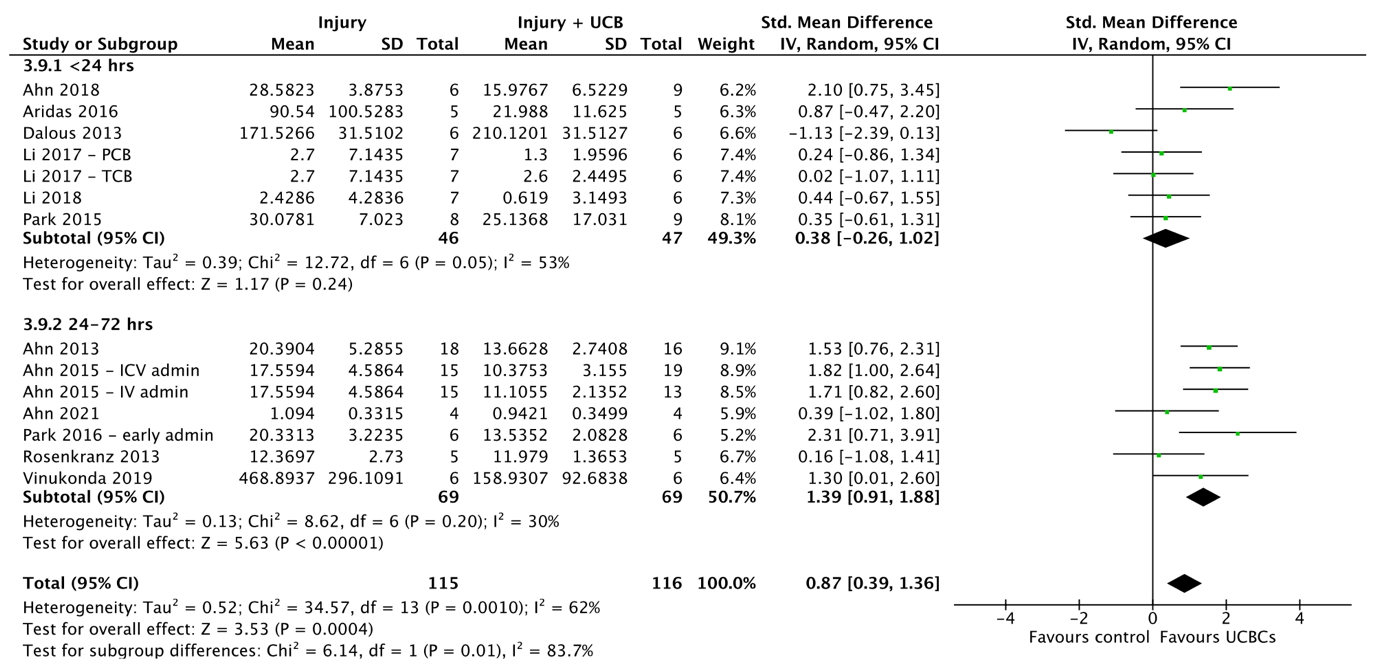

**(H)**

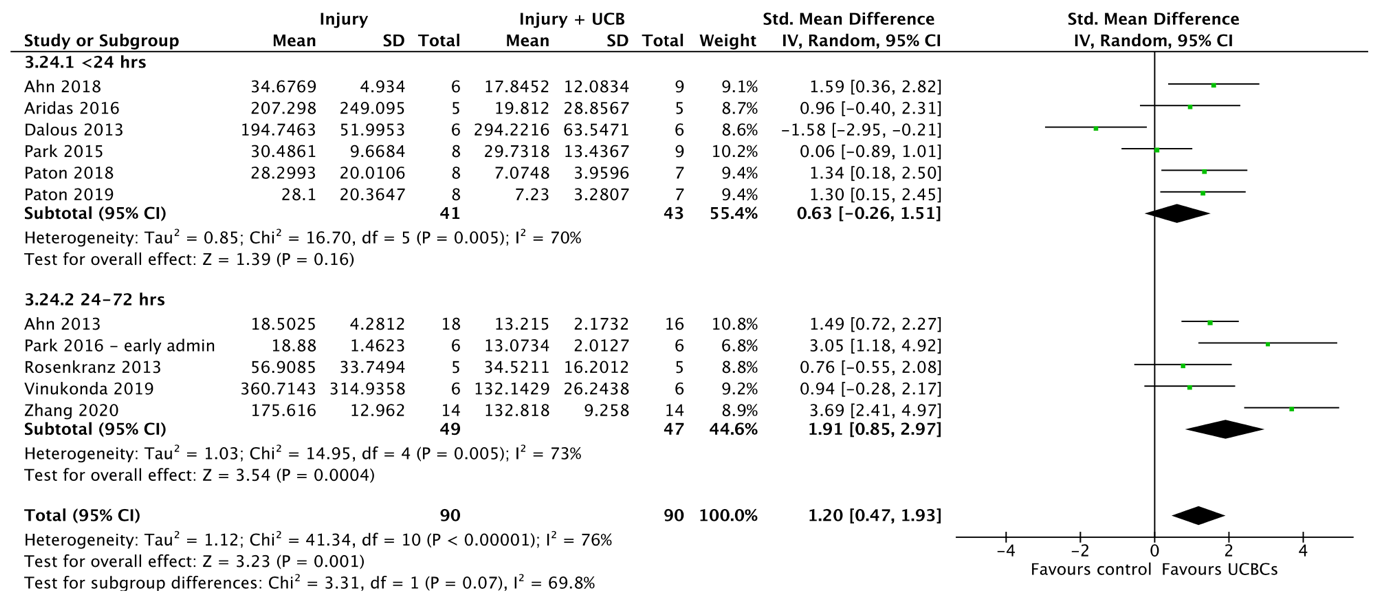

**Supplemental 6**. Forest plots demonstrating the effect of the route of cell administration on brain outcomes of **(A)** Apoptosis - grey matter; **(B)** Apoptosis - white matter; **(C)** Astrogliosis - grey matter; **(D)** Astrogliosis - white matter; **(E)** Infarct size; **(F)** Microglial activation - grey matter; **(G)**Neuroinflammation - TNF-α; **(H)** Motor function – cylinder test. Abbreviations: admin, administration; ECFC, endothelial colony-forming cell; EPC, endothelial progenitor cell; ICV, intracerebroventricular; IV, intraventricular; MNC, mononuclear cell; PCB, preterm cord blood; TCB, term cord blood; Treg, T regulatory cell.

**(A)**
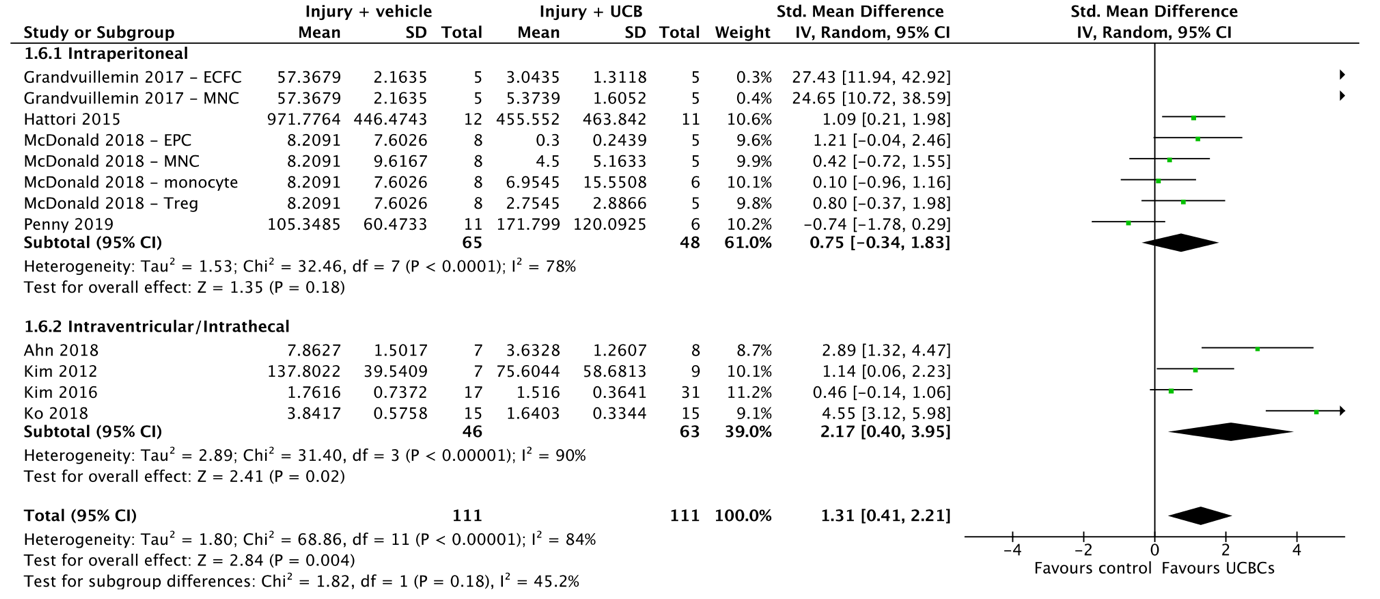

**(B)**
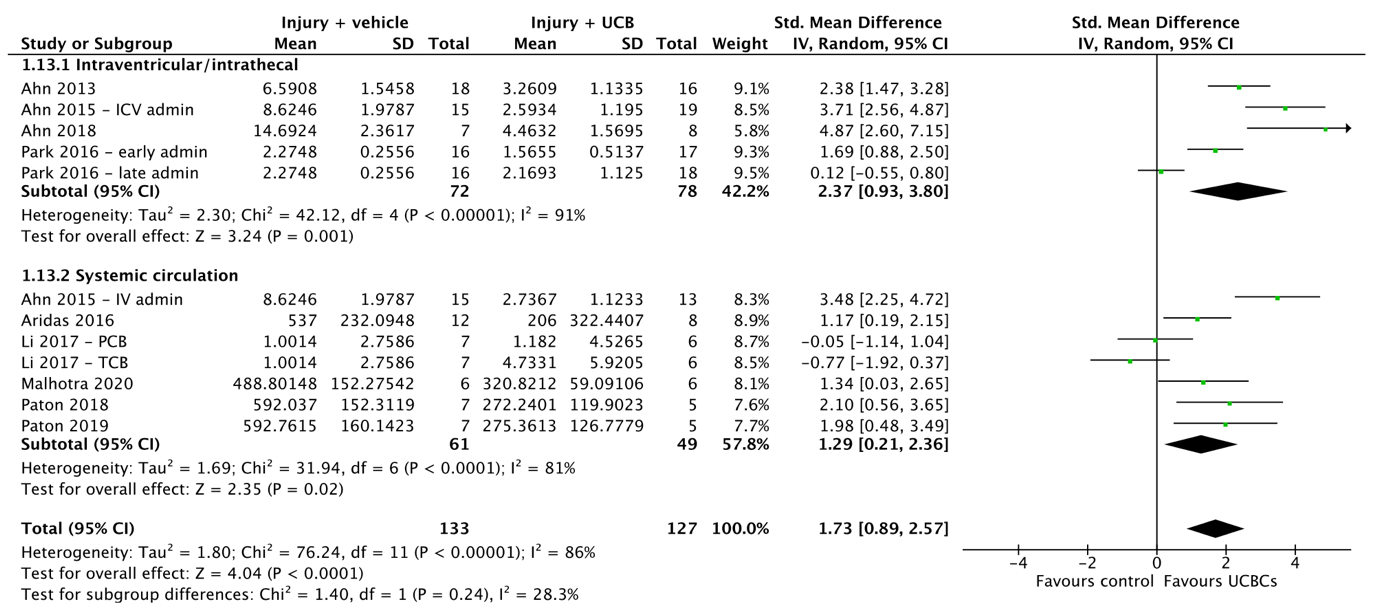

**(C)**
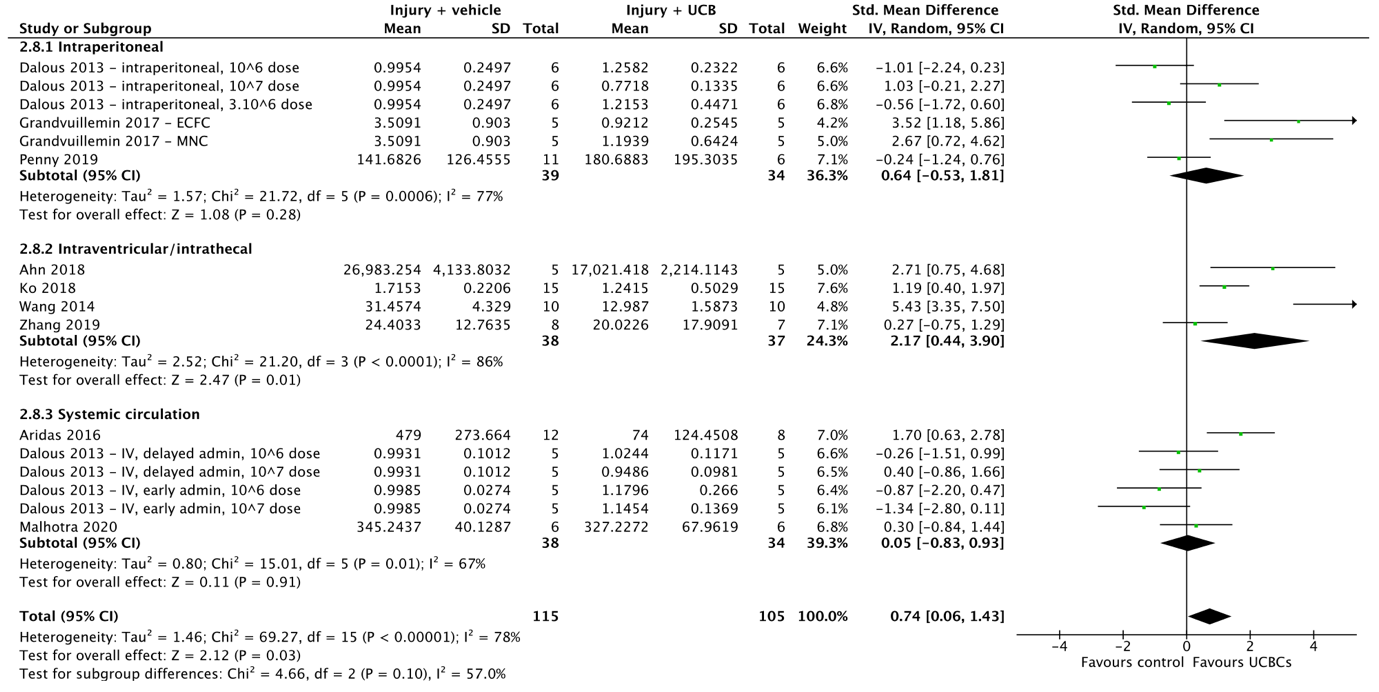

**(D)**

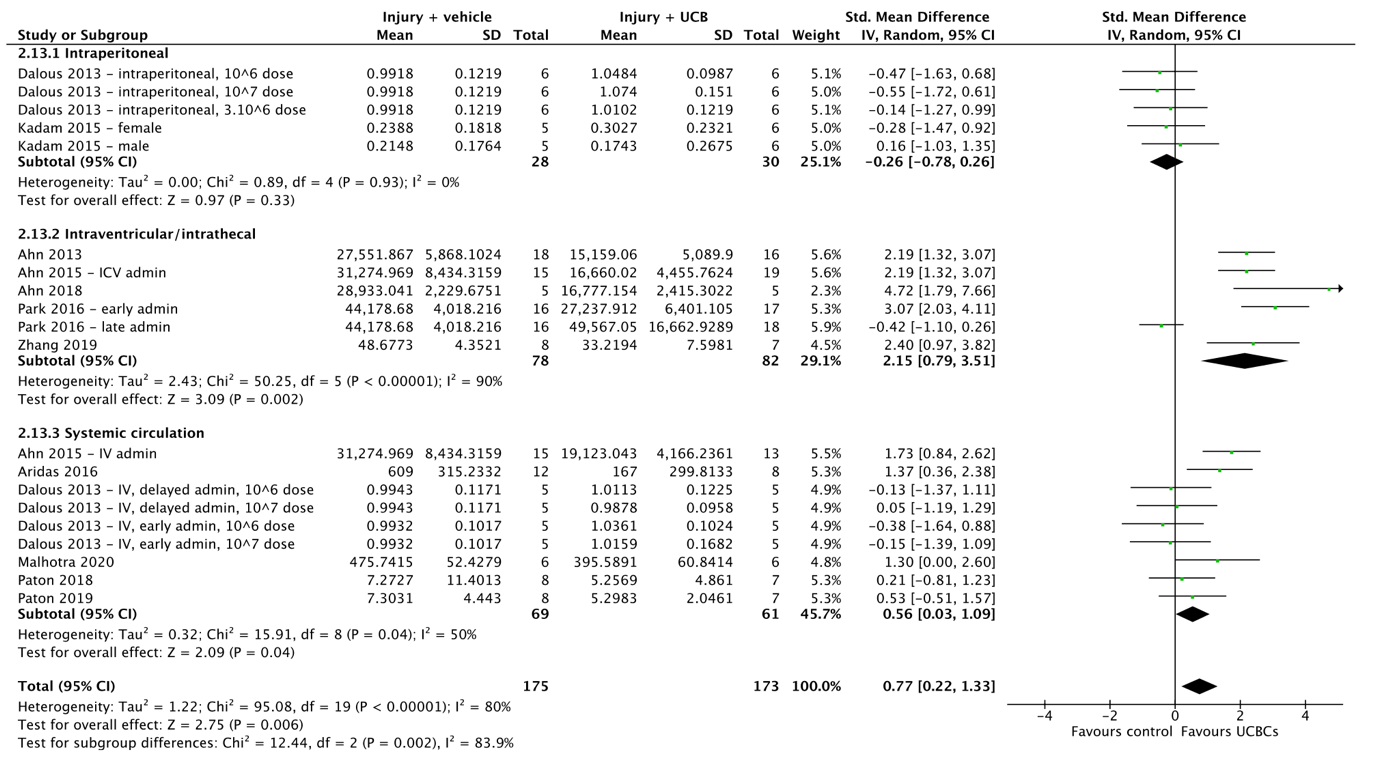

**(E)**

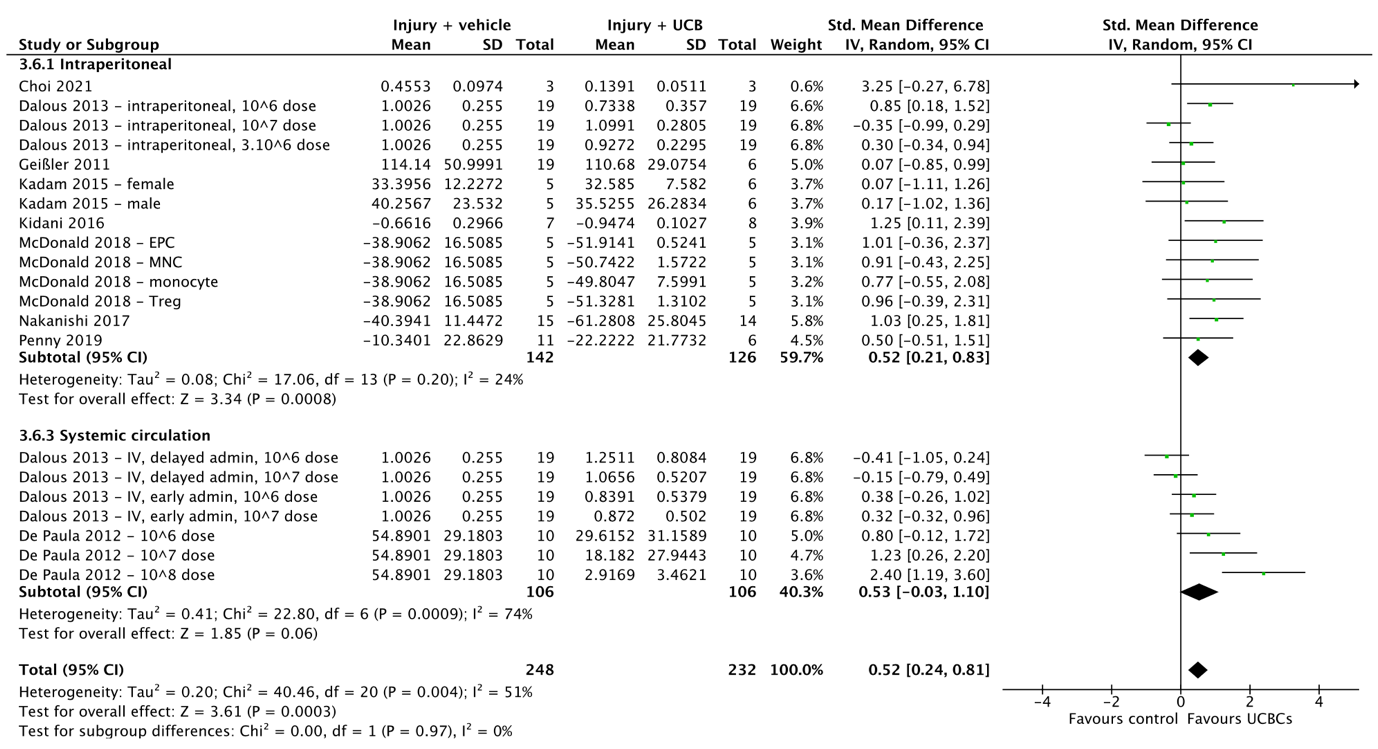

**(F)**

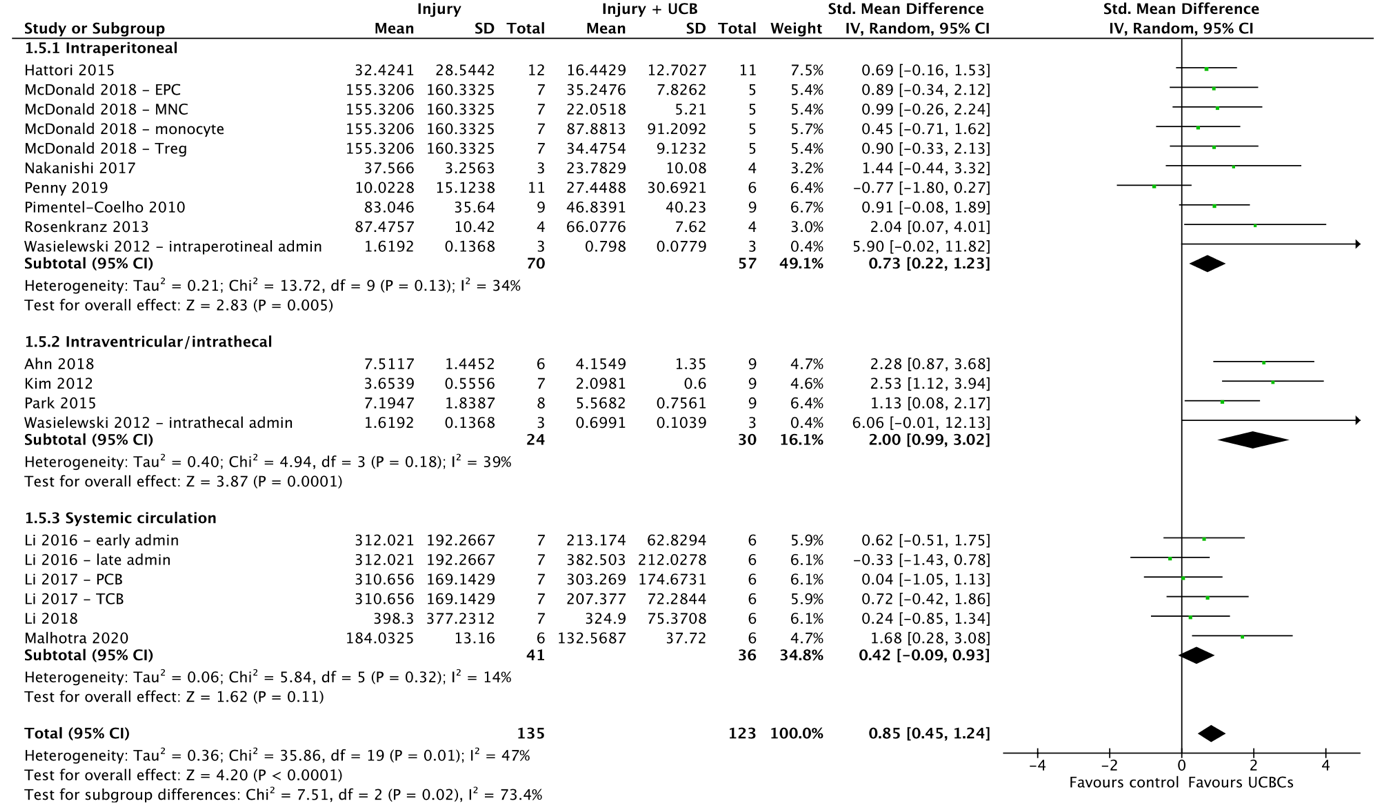

**(G)**

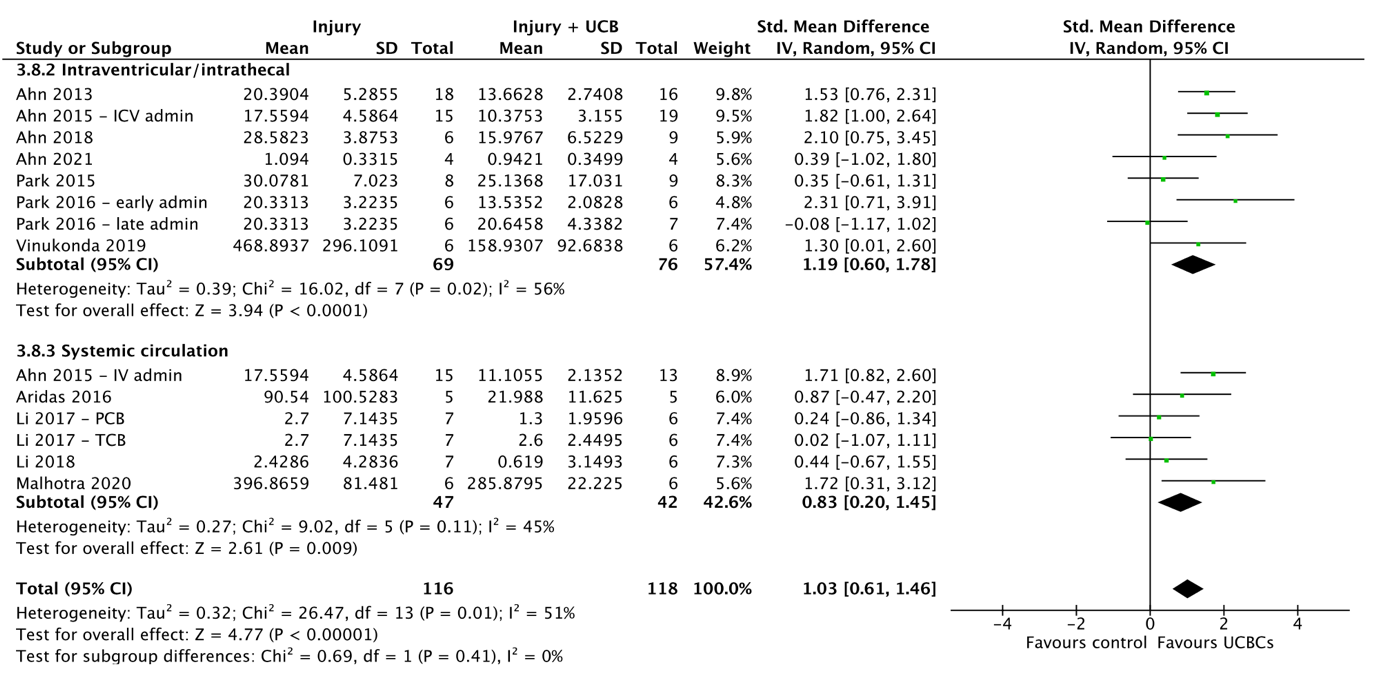

**(H)**
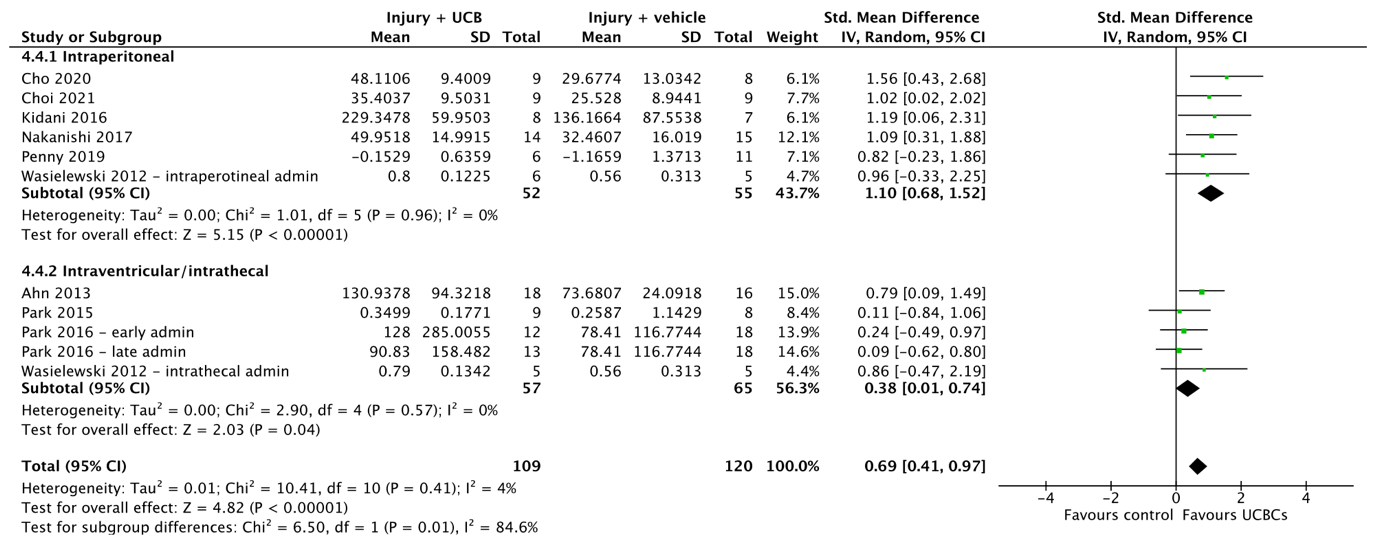

**Supplemental 7**. Forest plots demonstrating the effect of cell dosage on brain outcomes of **(A)** Apoptosis - grey matter; **(B)** Apoptosis - white matter; **(C)** Astrogliosis - grey matter; **(D)** Astrogliosis - white matter; **(E)** Infarct size; **(F)** Microglial activation - grey matter; **(G)** Neuron number; (**H)** Oligodendrocyte number – white matter; **(I)** Neuroinflammation - TNF-α; **(J)** Neuroinflammation - IL-1$\beta$; **(K)** Motor function – cylinder test. Abbreviations: admin, administration; ECFC, endothelial colony-forming cell; EPC, endothelial progenitor cell; ICV, intracerebroventricular; IV, intraventricular; MNC, mononuclear cell; million/kg, million cells per kilogram; PCB, preterm cord blood; TCB, term cord blood; Treg, T regulatory cell.

**(A)**

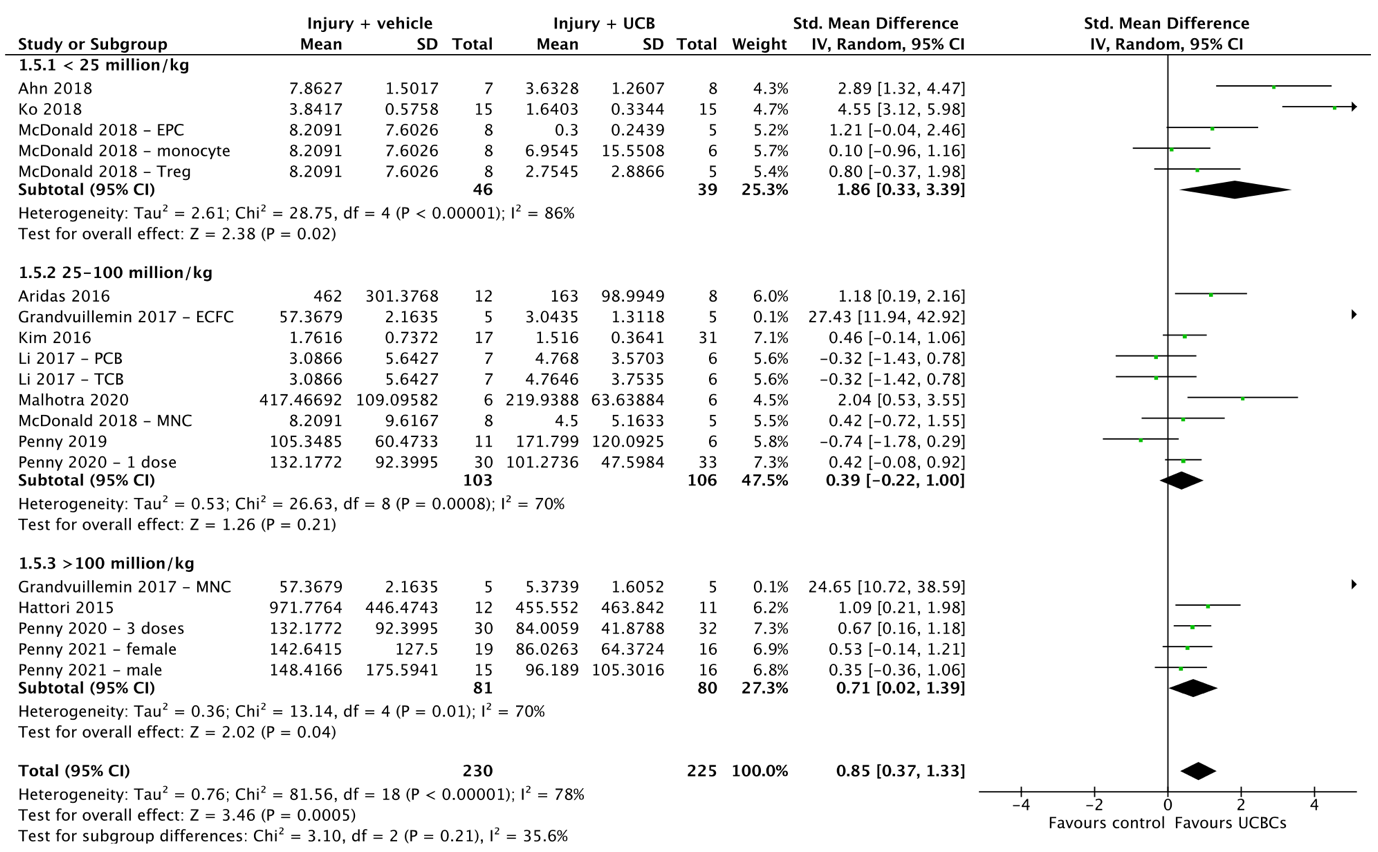

**(B)**

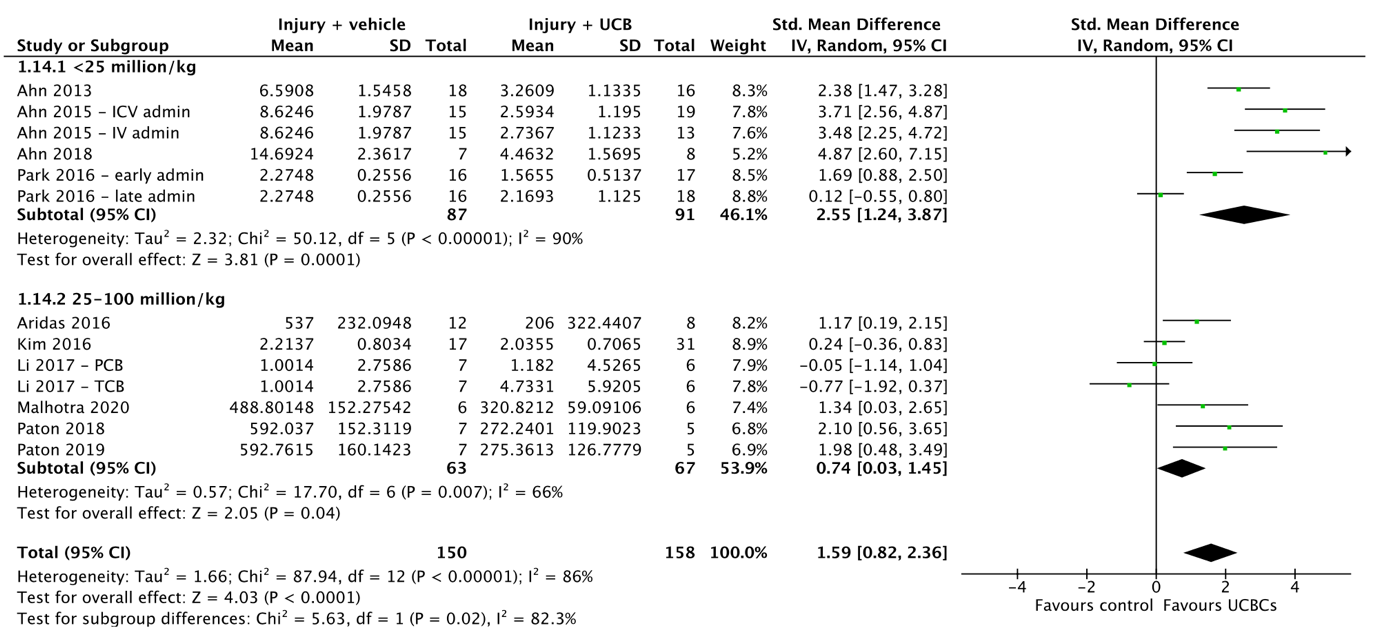

**(C)**

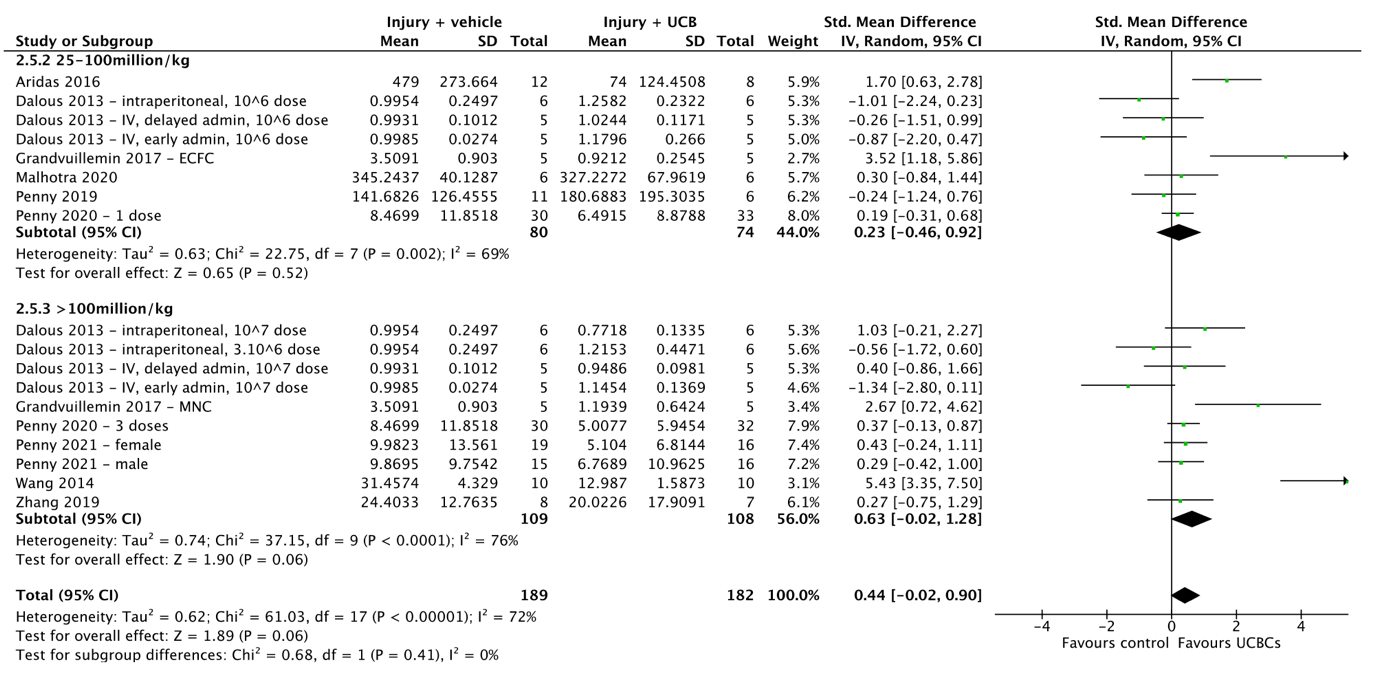

**(D)**

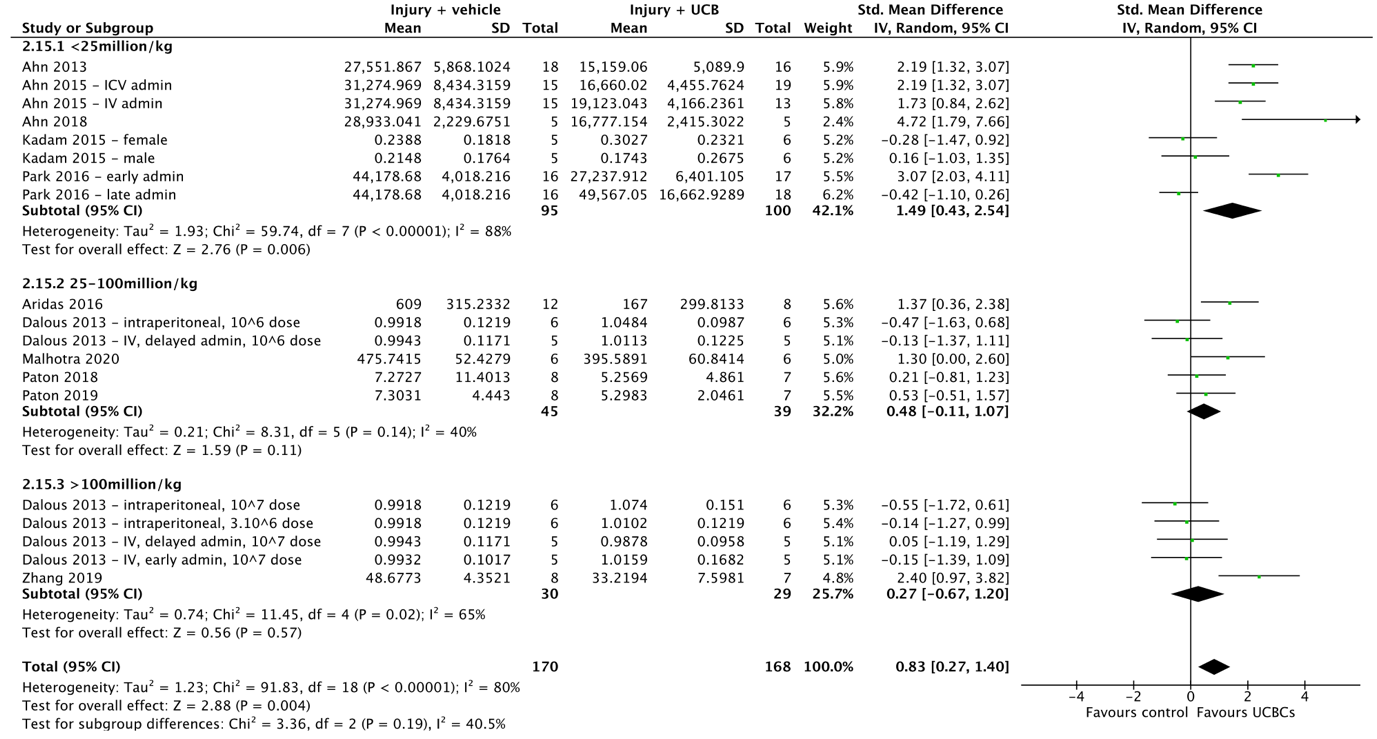

**(E)**

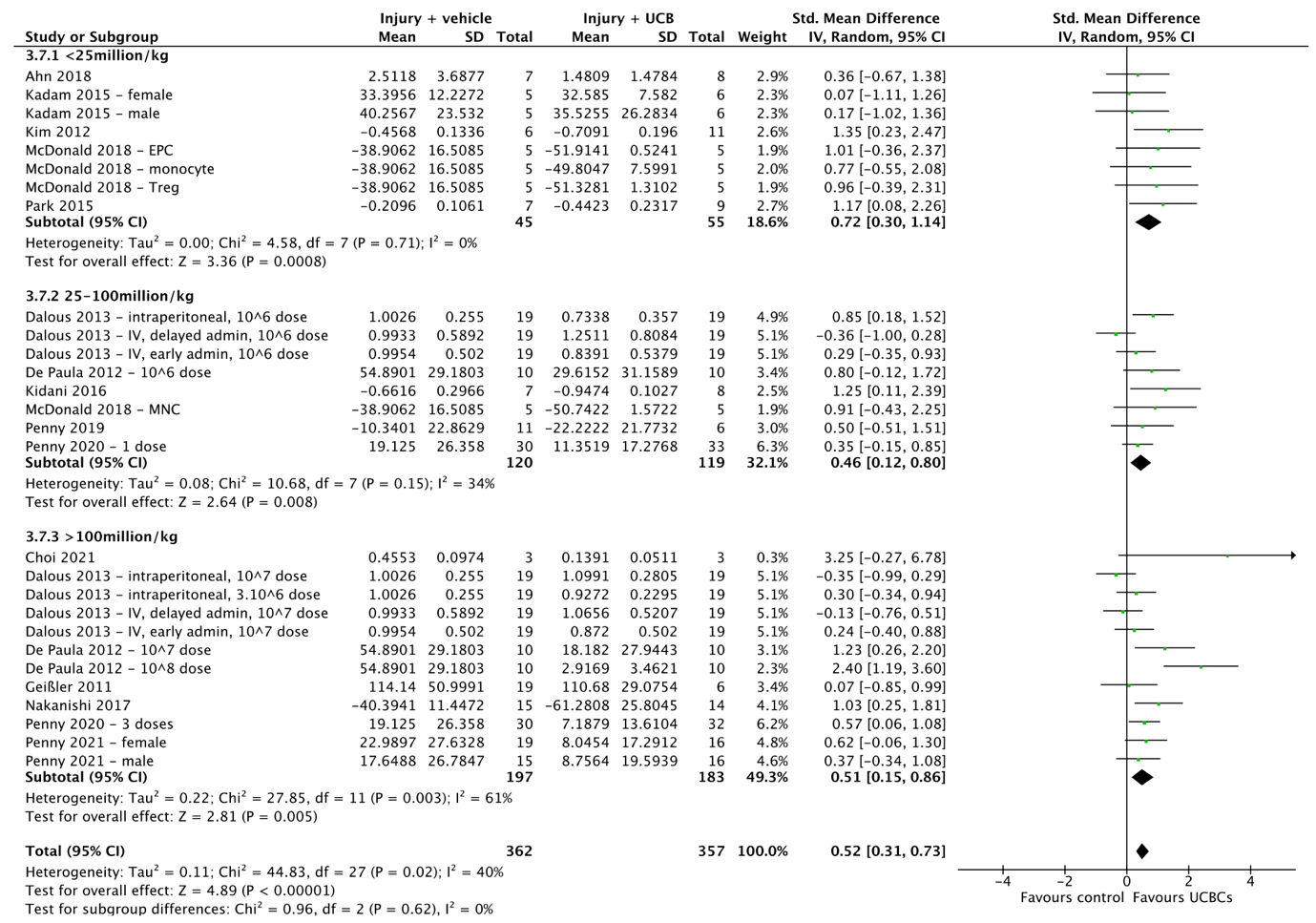

**(F)**
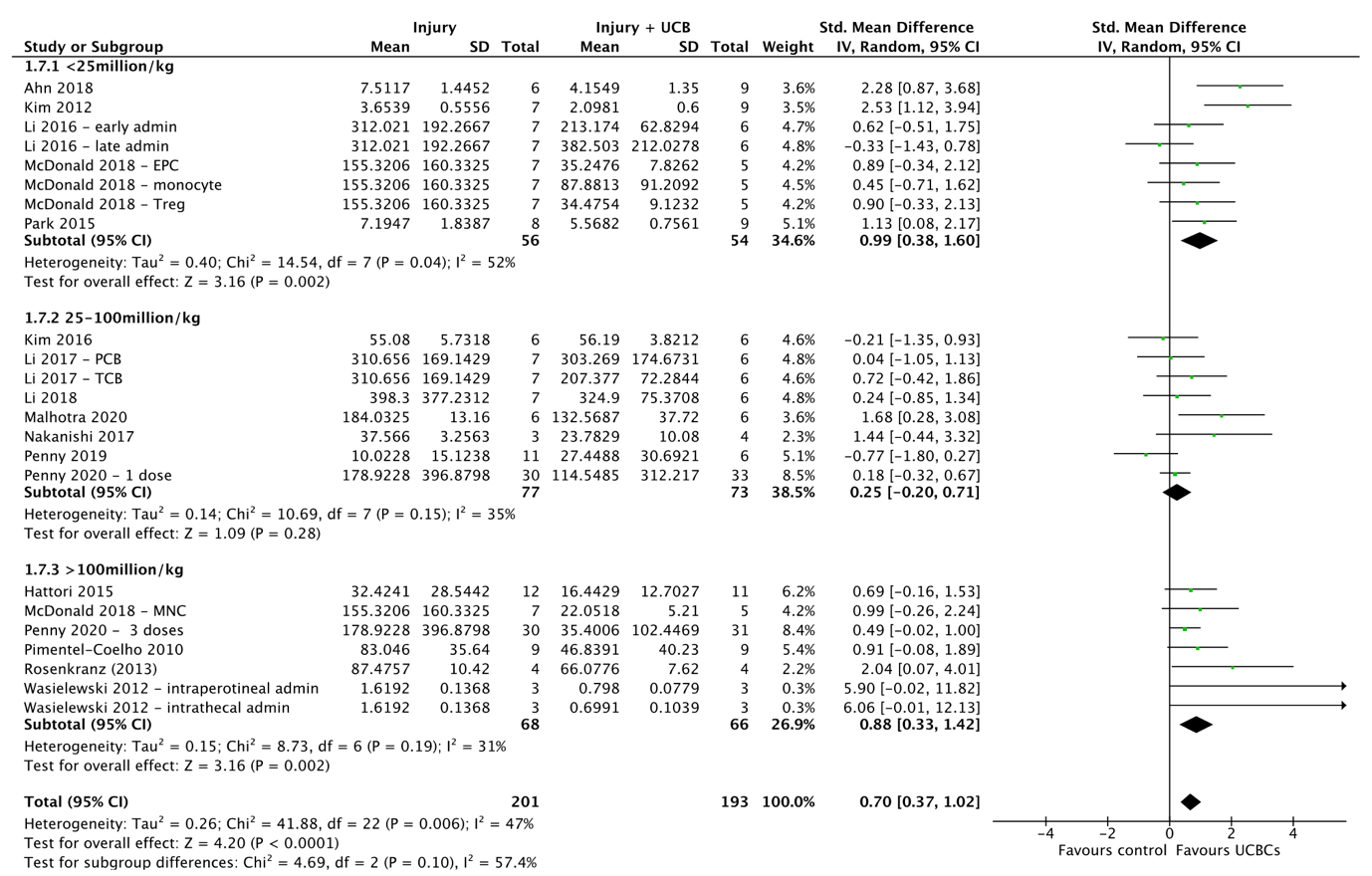

**(G)**

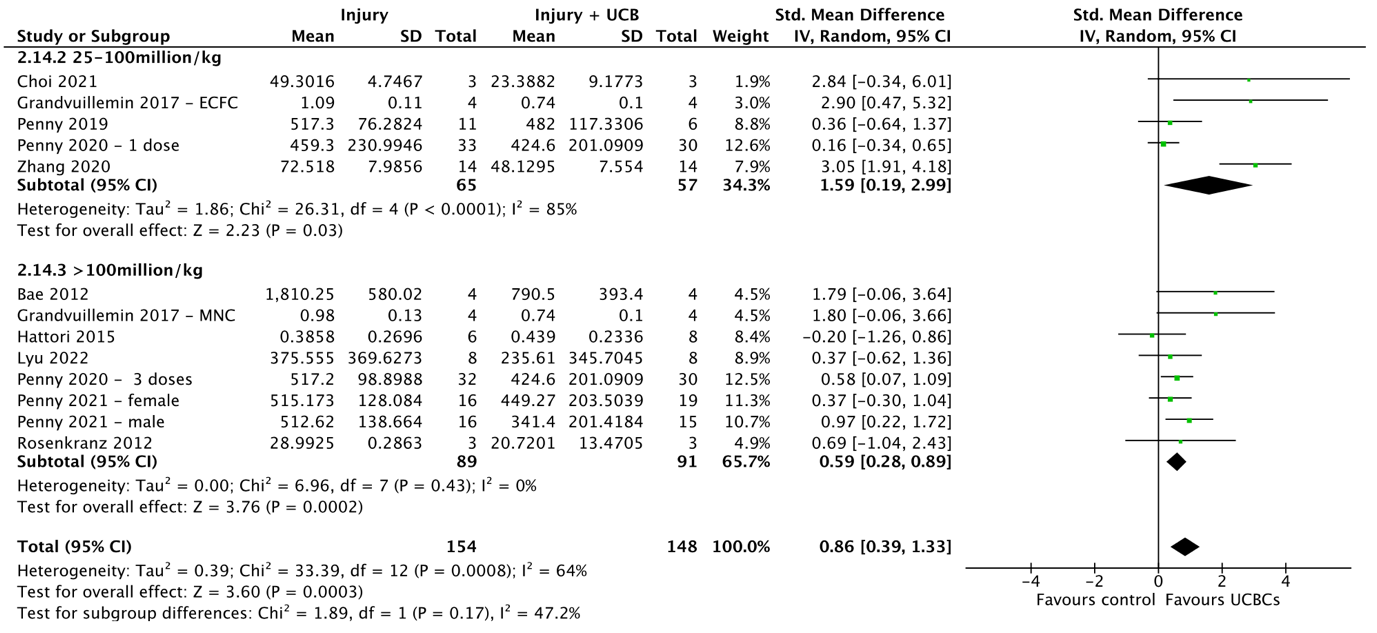

**(H)**
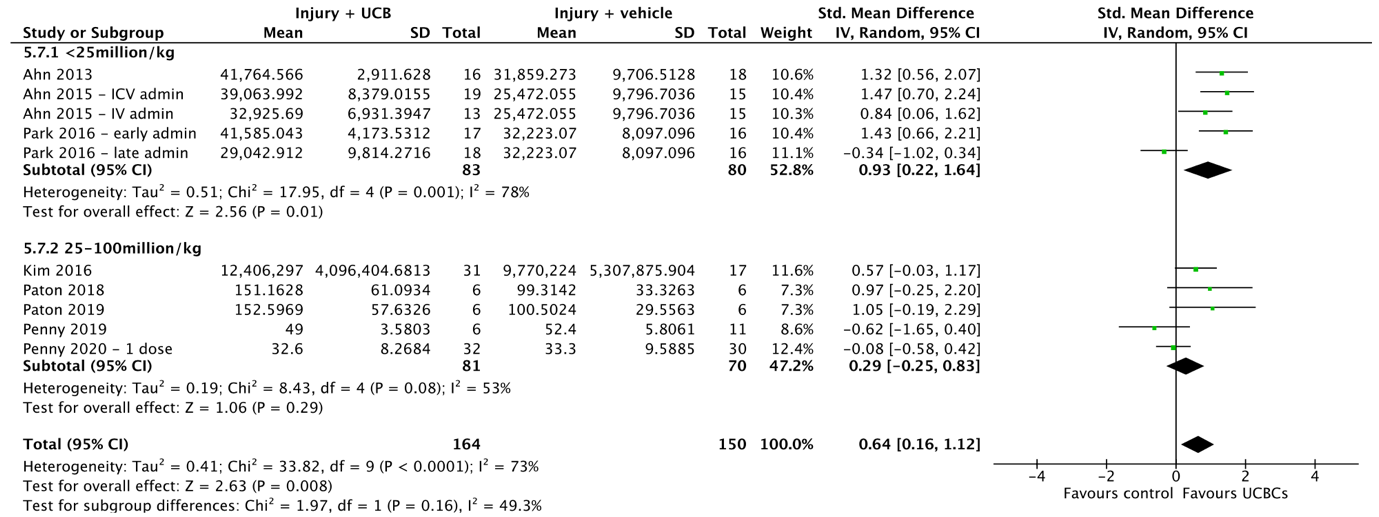

**(I)**

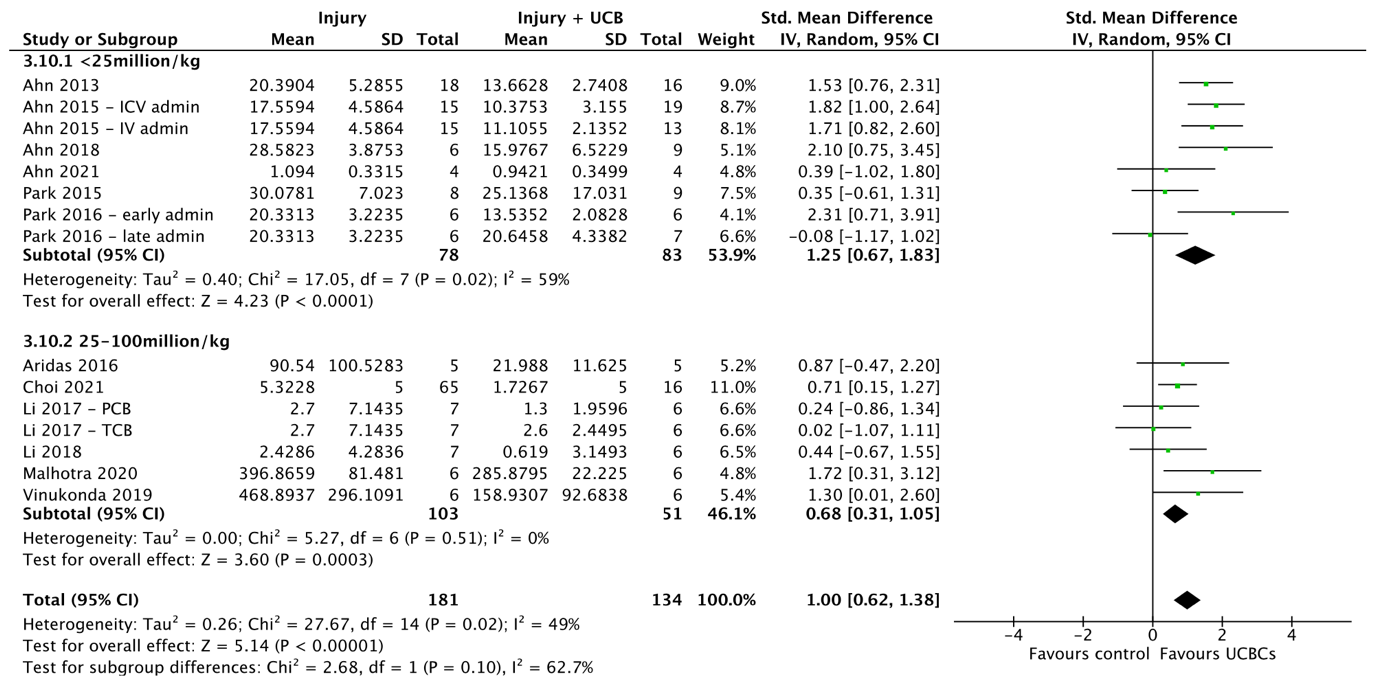

**(J)**

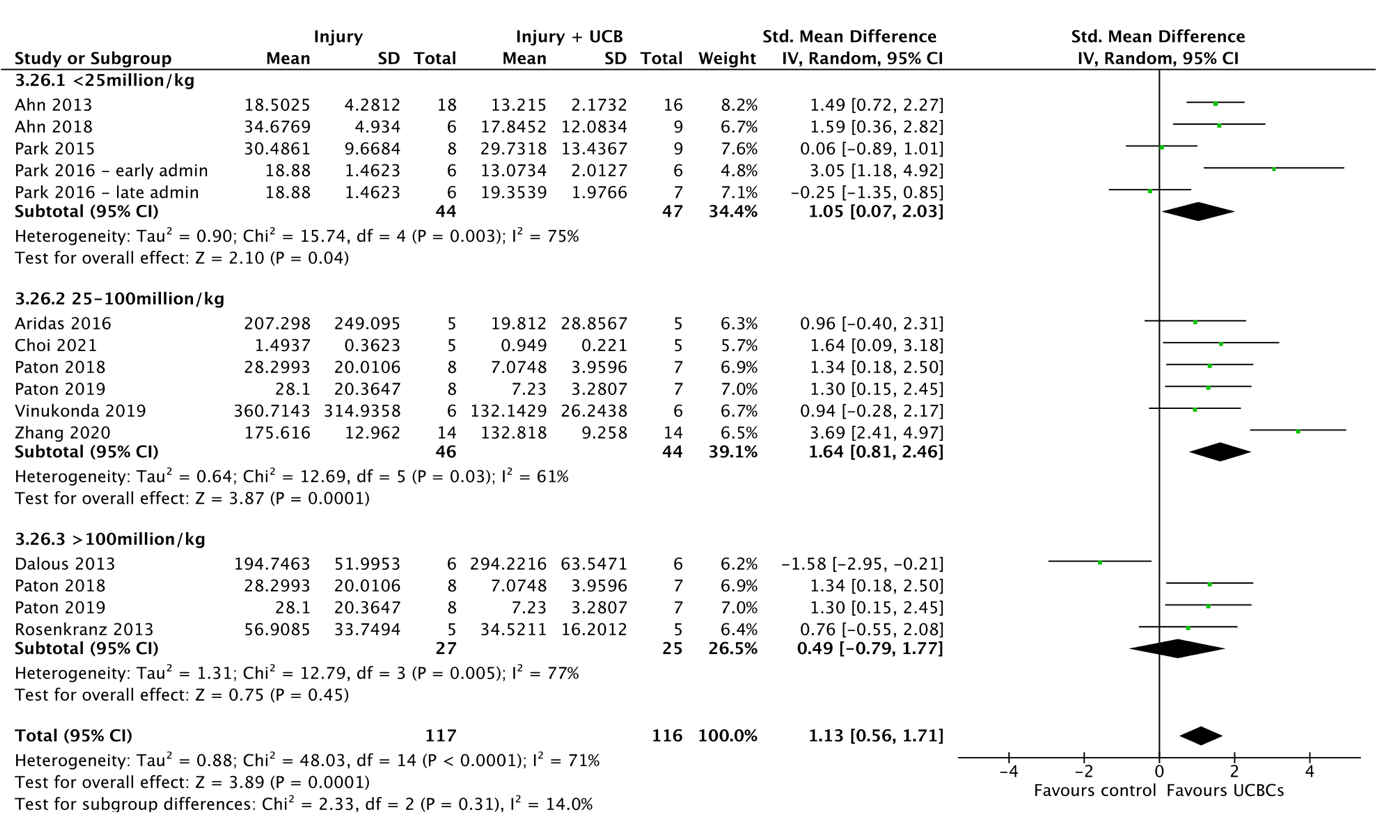

**(K)**

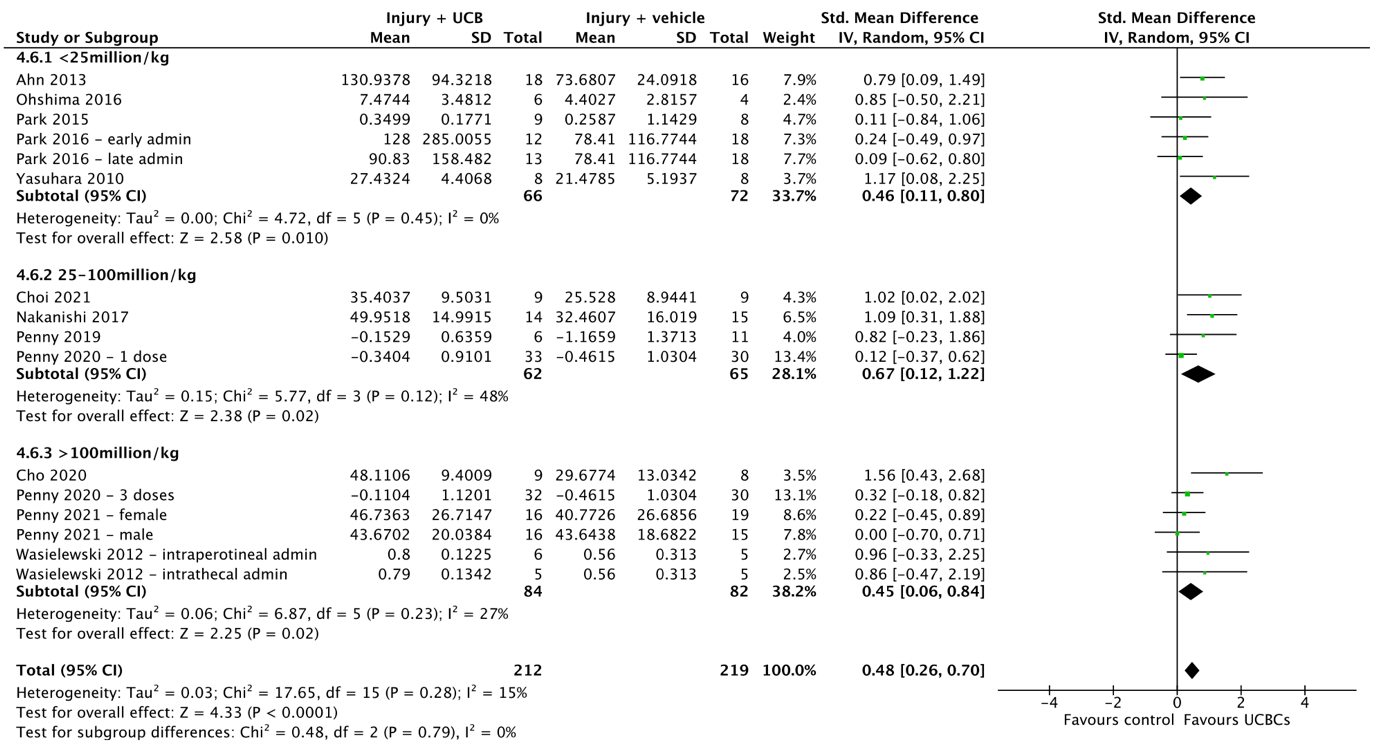

**Supplemental 8.** SYRCLE risk of bias assessment. Abbreviations: + = low risk of bias; ? = unclear risk of bias; - = high risk of bias.

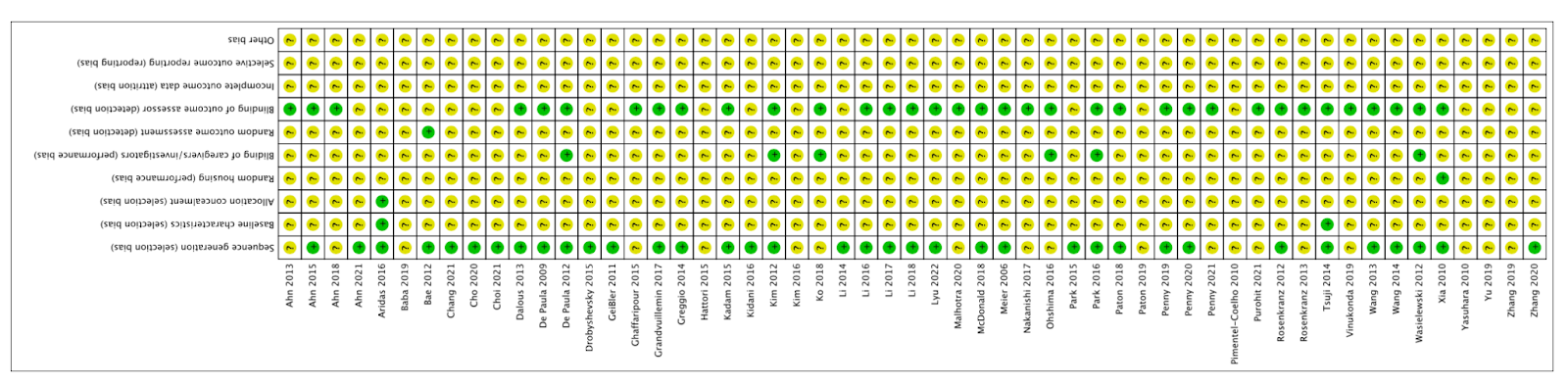
